# GeneWeld: a method for efficient targeted integration directed by short homology

**DOI:** 10.1101/431627

**Authors:** Wesley A. Wierson, Jordan M. Welker, Maira P. Almeida, Carla M. Mann, Dennis A. Webster, Melanie E. Torrie, Trevor J. Weiss, Macy K. Vollbrecht, Merrina Lan, Kenna C. McKeighan, Jacklyn Levey, Zhitao Ming, Alec Wehmeier, Christopher S. Mikelson, Jeffrey A. Haltom, Kristen M. Kwan, Chi-Bin Chien, Darius Balciunas, Stephen C. Ekker, Karl J. Clark, Beau R. Webber, Branden Moriarity, Staci L. Solin, Daniel F. Carlson, Drena L. Dobbs, Maura McGrail, Jeffrey J. Essner

## Abstract

Choices for genome engineering and integration involve high efficiency with little or no target specificity or high specificity with low activity. Here, we describe a targeted integration strategy, called GeneWeld, and a vector series for gene tagging, pGTag (<u>p</u>lasmids for <u>G</u>ene <u>Tag</u>ging), which promote highly efficient and precise targeted integration in zebrafish embryos, pig fibroblasts, and human cells utilizing the CRISPR/Cas9 system. Our work demonstrates that *in vivo* targeting of a genomic locus of interest with CRISPR/Cas9 and a donor vector containing as little as 24 to 48 base pairs of homology directs precise and efficient knock-in when the homology arms are exposed with a double strand break *in vivo*. Our results suggest that the length of homology is not important in the design of knock-in vectors but rather how the homology is presented to a double strand break in the genome. Given our results targeting multiple loci in different species, we expect the accompanying protocols, vectors, and web interface for homology arm design to help streamline gene targeting and applications in CRISPR and TALEN compatible systems.

## Introduction

Designer nucleases have rapidly expanded the way in which researchers can utilize endogenous DNA repair mechanisms for creating gene knock-outs, reporter gene knock-ins, gene deletions, single nucleotide polymorphisms, and epitope tagged alleles in diverse species (Bedell et al., 2012; Beumer et al., 2008; Carlson et al., 2012; Geurts et al., 2009; Yang et al., 2013). A single dsDNA break in the genome results in increased frequencies of recombination and promotes integration of homologous recombination (HR)-based vectors (Hasty et al., 1991; Hoshijima et al., 2016; Orr-Weaver et al., 1981; Rong and Golic, 2000; Shin et al., 2014; Zu et al., 2013). Additionally, *in vitro* or *in vivo* linearization of targeting vectors stimulates homology-directed repair (HDR) (Hasty et al., 1991; Hoshijima et al., 2016; Orr-Weaver et al., 1981; Rong and Golic, 2000; Shin et al., 2014; Zu et al., 2013). Utilizing HDR or HR at a targeted double-strand break (DSB) allows base-pair precision to directionally knock-in exogenous DNA, however, frequencies remain variable and engineering of targeting vectors is cumbersome.

Previous work has shown *Xenopus* oocytes have the ability to join or recombine linear DNA molecules that contain short regions of homology at their ends, and this activity is likely mediated by exonuclease activity allowing base pairing of the resected homology (Grzesiuk and Carroll, 1987). More recently, it was shown in *Xenopus*, silkworm, zebrafish, and mouse cells that a plasmid donor containing short (≤40 bp) regions of homology to a genomic target site can promote precise integration at the genomic cut site when the donor plasmid is cut adjacent to the homology (Aida et al., 2016; Hisano et al., 2015; Nakade et al., 2014). Gene targeting is likely mediated by the alternative-end joining/microhomology-mediated end joining (MMEJ) pathway or by a single strand annealing (SSA) mechanism (Ceccaldi et al., 2016). In contrast, in human cell culture, linear donors using a similar strategy with homologous ends have been reported to show inefficient integration until homology domains reach ~600 bp (Zhang et al., 2017), suggesting that different repair pathways may predominate depending on cell type. In the initial reports using short regions of homology for *in vivo* gene targeting in zebrafish, the level of mosaicism in F0 injected animals was high, resulting in inefficient recovery of targeted alleles through the germline (Aida et al., 2016; Hisano et al., 2015; Nakade et al., 2014).

Here, we present GeneWeld, a strategy for targeted integration directed by short homology, and demonstrate increased germline transmission rates for recovery of targeted alleles. We provide a detailed protocol and a suite of donor vectors, called pGTag, that can be easily engineered with homologous sequences (hereafter called homology arms) to a gene of interest, and a web interface for designing homology arms (www.genesculpt.org/gtaghd/). We demonstrate that 24 or 48 base pairs of homology directly flanking cargo DNA promotes efficient gene targeting in zebrafish, pig, and human cells with frequencies up to 10-fold higher than other HR strategies. Our results also suggest that longer homology arms up to 1 kb in length provide no advantage to knock-in frequencies over short homology, and the important aspect of knock-in design is the ability to expose the homology on the knock-in cassette ends. Using short homology-arm mediated end joining, we can achieve germline transmission rates averaging approximately 50% across several zebrafish loci. Southern blot analysis in the F1 generation reveals that the GeneWeld strategy can yield alleles with precise integration at both 5’ and 3’ ends, as well as alleles that are precise on just one end. Finally, we present a strategy to delete and replace up to 48kb of genomic DNA with a donor containing homology arms flanking two distal CRISPR/Cas9 sites in a gene. These tools and methodology provide a tractable solution to creating precise targeted integrations and open the door for other genome editing strategies using short homology.

## Results

The GeneWeld strategy takes advantage of two simultaneous actions to initiate targeted integration directed by short homology (Fig. 1a). First, a high efficiency nuclease introduces a DSB in the chromosomal target. Simultaneously, a second nuclease makes a DSB in the pGTag vector integration cassette exposing the short homology arms. The complementarity between the chromosomal DSB and the donor homology arms activates a MMEJ/SSA or other non-NHEJ DNA repair mechanism, together referred to as homology-mediated end joining (HMEJ). The reagents needed for this gene targeting strategy include *Cas9* mRNA to express the Cas9 nuclease, a guide RNA targeting the genomic sequence of interest, a universal gRNA (UgRNA) that targets two sites in the pGTag series donor vectors to expose the homology arms, and a pGTag/donor vector with gene specific homology arms (Fig. 1a). The universal gRNA (UgRNA) has no predicted sites in zebrafish, pig, or human genomes. Alternatively, a gene specific guide RNA can be used to expose homology arms in the donor vector. For simplicity we will refer to this set of reagents as ‘GeneWeld reagents’. Using GeneWeld reagents to target various loci, we demonstrate widespread reporter gene expression in injected F0 zebrafish embryos, porcine fibroblasts, and human K-562 cells that correlates with efficient and precise in-frame integration in multiple species and cell systems.

**Figure 1.**
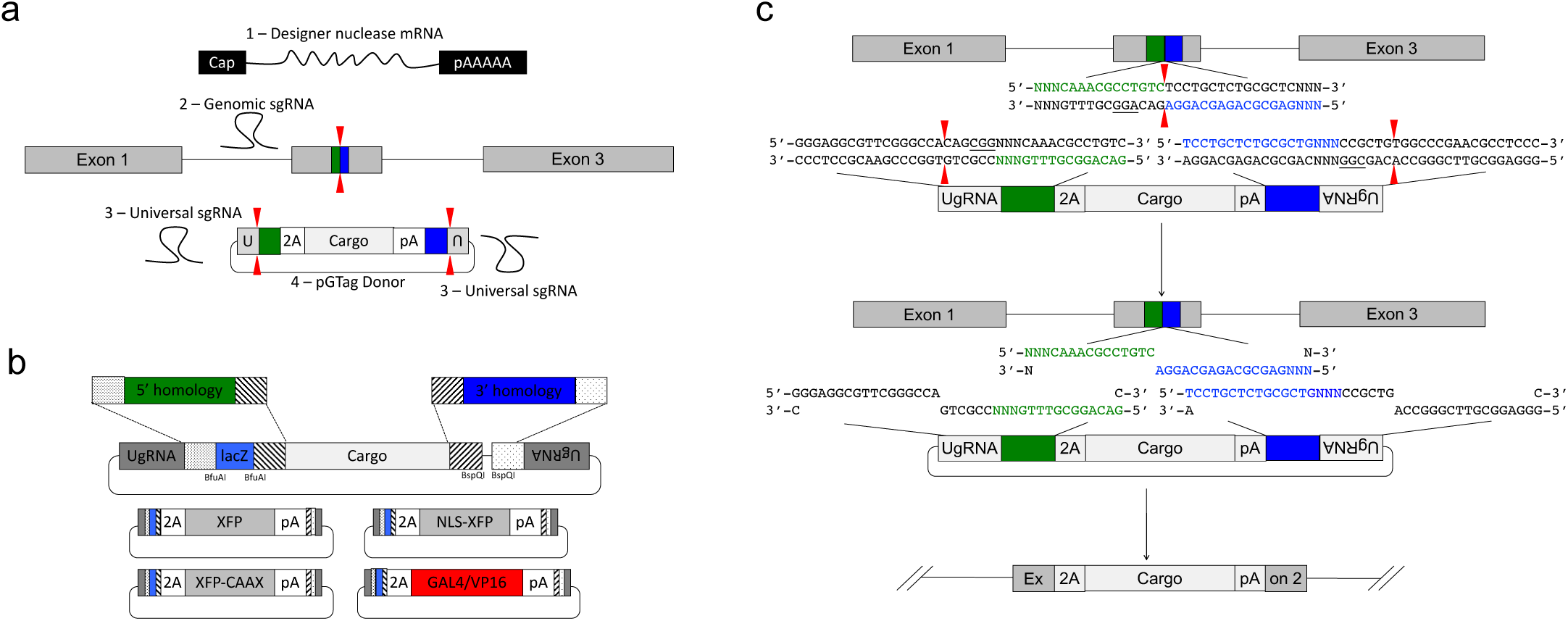
GeneWeld strategy and pGTag vector series. (a) GeneWeld reagent components are designed for simultaneous genome and donor nuclease targeting to reveal short regions of homology. Red arrowheads represent *in vivo* designer nuclease DSBs. Components include: 1 - Designer nuclease mRNA, either Cas9 to target both the genome and donor, or Cas9 to target the donor and TALEN to cut the genome; 2 - sgRNA for targeting Cas9 to genome; 3 - Universal sgRNA to liberate donor cargo and homologous ends; and 4 - pGTag donor of interest with short homology arms. (b) Type IIs restriction endonucleases *Bfu*AI and *Bsp*QI create incompatible ends outside of their recognition sequence, allowing digestion and ligation of both homology arms into the vector in a single reaction. Homology arm fragments are formed by annealing complementary oligonucleotides to form dsDNA with sticky ends for directional cloning into the vector. XFP = Green or Red Fluorescent Protein. pA = SV40 or β-actin 3’ untranslated region. Red and green fluorescent proteins were cloned into the pGTag vectors, and for each color, subcellular localization sequences for either nuclear localization (NLSs) and membrane localization (CAAX) are provided (c) Schematic of GeneWeld targeting *in vivo*. After designer nuclease creates targeted double-strand breaks in the genome and donor, end resection likely precedes homology recognition and strand annealing, leading to integration of the donor without vector backbone. Red arrowheads represent *in vivo* designer nuclease DSBs.

### A single 48bp short homology domain drives efficient CRISPR targeted integration

To develop baseline gene targeting data, we engineered variable length homology domains to target *noto*. Homology lengths were based on observations that DNA repair enzymes bind DNA and search for homology in 3 or 4 base pair lengths (Fig. S1a) (Conway et al., 2004; Singleton et al., 2002). Injection of a *noto* sgRNA that targets *noto* exon 1 and the 5’ homology domain cloned into a 2A-TagRFP-CAAX-SV40 donor vector, efficient targeted integration was observed as notochord-specific RFP expression (Fig. S1a-c; Table S1–S3). The frequency of embryos with notochord-specific RFP expression increased with the length of the homology arm up to 48 bp (Fig. S1b). Somatic junction fragment analysis revealed precise integration efficiencies reaching 95% of sequenced alleles (Fig. S1d), indicating a strong correlation between expression of the reporter gene and precise somatic targeting of *noto*.

Following these initial experiments, a 3 bp spacer sequence was included in all homology arm designs in order to separate the donor CRISPR/Cas9 target PAM and the homology domain (Fig. S2a). The spacer was included to prevent arbitrarily increasing the length of the targeting domain, since single base pair alterations in the homology region affected the tissue specific expression and presumed knock-in efficiency up to 2-fold in somatic tissue (Fig. S2b). Based on these observations the following experiments were carried out with either 24 or 48 base pairs of homology to the double strand break in the targeted gene.

### A universal guide RNA to liberate donor homology for targeted integration

To simplify donor design and liberate donor cargo *in vivo* with reproducible efficiency, a universal guide RNA sequence UgRNA, with no predicted targets in zebrafish, pig, or human genomes, was designed using optimal base composition in CRISPRScan (Fig. S3a) (Moreno-Mateos et al., 2015). To test the ability of the guide to direct a Cas9 double strand break and efficient targeted integration, the UgRNA and CGG PAM sequences were cloned 5’ to the *noto* homology arm in the 2A-TagRFP-CAAX-SV40 donor vector (Fig. S3b). Zebrafish embryos injected with Cas9 mRNA, UgRNA, *noto* sgRNA, and donor plasmid resulted in 21% of injected embryos showing notochord RFP, suggesting that Cas9 with the UgRNA efficiently exposes the donor vector 5’ homology arm and drive precise targeted integration (Fig. S3c). The high frequency of notochord-specific RFP-positive embryos following injection suggests repair of the DSB preferentially utilizes homology in the targeting construct over the NHEJ pathway.

### Dual homology arm liberation directs precise 5’ and 3’ integration in somatic tissue

We leveraged the activity of the UgRNA to develop GeneWeld, a strategy for targeted integration that results in high frequency precision repair at both 5’ and 3’ junctions at the target site. We built a series of vectors, pGTag, which contain sites on both sides of the cargo for cloning a short homology arm that is complementary to the 5’ or 3’ sequence flanking the genomic target site. The vectors also include the UgRNA sequence outside the sites for homology arm cloning (Fig. 1a, b). The final donor targeting vector will contain a cargo flanked by 5’ and 3’ homology arms with UgRNA sequences on both ends. Cleavage by Cas9 at the UgRNA sites liberates the DNA cargo from the plasmid backbone and exposes both 5’ and 3’ donor homology arms for interaction with DNA on either side of the genomic DSB (Fig. 1c).

We extended our analysis of the GeneWeld targeted integration strategy to four genes in zebrafish, *noto*, *tyrosinase (tyr)*, *esama (endothelial cell adhesion molecule a)*, and *connexin 43.4* (*cx43.4*), and measured the frequency of precise targeted integration in somatic tissue (Fig. 2 a-d). Injection of 24 or 48 bp homology arm *noto* 2A-eGFP-SV40 donors resulted in 24% of zebrafish embryos showing extensive reporter expression in the notochord (Fig. 2 a, e), suggesting a similar precise integration efficiency compared to targeting with the single 5’ homology arm 2A-TagRFP-CAAX-SV40 vector (Fig. S1, Table 1, 2). The results also suggest 24 bp of homology directs targeted integration as efficiently as 48 bp, further simplifying construction of GeneWeld vectors.

**Figure 2.**
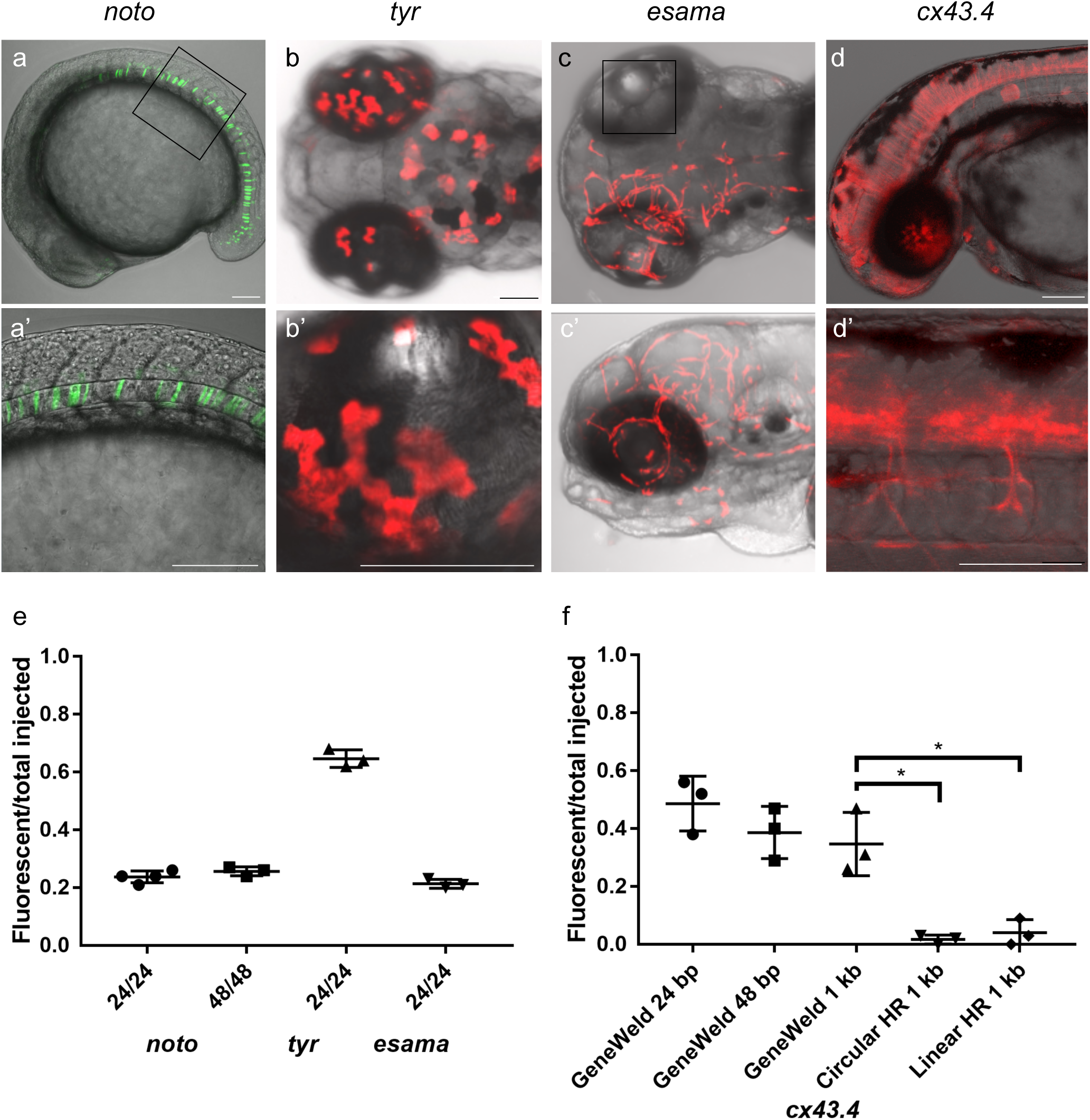
HMEJ strategy promotes efficient somatic targeting of knock-in cassettes in zebrafish. (a-d) Live confocal images of F0 injected embryos showing fluorescent reporter expression after GeneWeld targeted integration. *noto-2A-eGFP-SV40* at mid somite stage (a, a’). *tyr-2A-Gal4VP16-β-actin*;*Tg(UAS:mRFP*)^tpl2^ at 5 days post fertilization (dpf) (b, b’). *esama-2A-Gal4VP16-β-actin;Tg(UAS:mRFP*)^tpl2^ at 2 dpf (c) and 3 dpf (c’). *cx43.4-2A-tagRFP-CAAX-SV40* at 31 hours post fertilization (d, d’). (e) Frequency of embryos with reporter gene expression following GeneWeld targeting at *noto*, *tyr* and *esama*. 5’ and 3’ homology lengths flanking donor cargos indicated in base pairs as 24/24 or 48/48. (f) Comparison of the frequency of RFP expressing embryos after targeting *cx43.4* exon 2 using GeneWeld 24/24 bp homology, GeneWeld 48/48 bp homology, Geneweld 1kb/1kb homology, Circular HR 1kb/1kb (injection did not include UgRNA, \**p*=0.0067), Linear HR 1kb/1kb (donor was linearized before injections, \**p*=0.0111). Data represents mean +/− s.e.m. of 3 independent targeting experiments. *p* values calculated using Students *t* test. Scale bars, 100 μm.

**Table 1.**
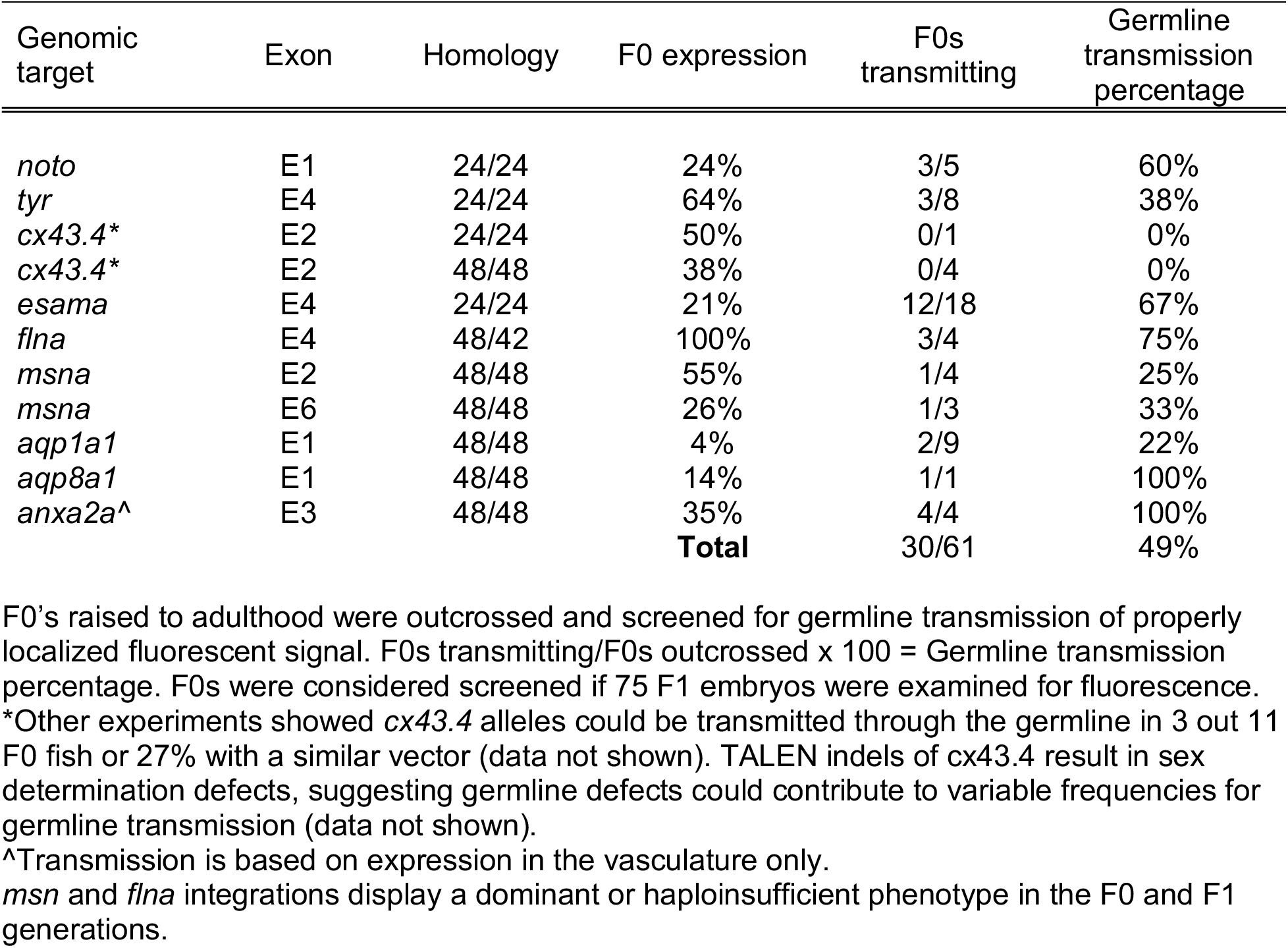
Germline transmission of zebrafish GeneWeld GTag integrations.

Targeting exon 4 of *tyr* or exon 2 of *esama* with a 24 bp homology arm 2A-TagRFP-CAAX-SV40 donor did not result in detectable RFP signal, similar to previous reports for *tyr* (Hisano et al., 2015). However, PCR junction fragments from injected embryos showed the donor was precisely integrated in frame into *tyr* exon 4 (Fig. S4), suggesting RFP expression was below the threshold of detection. To amplify the fluorescent signal, we built pGTag 24 bp homology arm 2A-Gal4VP16-β−actin3’UTR donors to integrate the Gal4VP16 trans-activator into the *tyr* and *esama* target sites, and injected into transgenic zebrafish embryos carrying a 14xUAS-RFP reporter, *Tg*(*UAS:mRFP*)^tpl2^ (Balciuniene et al., 2013). This resulted in strong RFP signal in 64% of *tyr* injected animals (Fig. 2b, e), however, the embryos were highly mosaic with only 9% of RFP embryos displaying extensive expression throughout most of the pigmented cells. Targeting *esama* exon 2 with 2A-Gal4VP16-β−actin3’UTR in the *Tg*(*UAS:mRFP*)^tpl2^ transgenic background resulted in 21% of embryos displaying extensive RFP expression specifically in the vasculature (Fig. 2c, e). This approach was further extended to five additional loci, targeting 2A-Gal4VP16 to *filamin a* (*flna*) exon 4, *moesin a* (*msna*) exon 2 and 6, *aquaporin 1a1* (*aqp1a1*) exon 1, *aquaporin 8a1* (*aqp8a1*) exon 1, and *annexin a2a* (*anxa2a*) exon 3. At these loci, transient expression of RFP was observed following injection in 4-55% of *Tg*(*UAS:mRFP*)^tpl2^ embryos (Table S1 and S2). Taken together, these results suggest that the GeneWeld targeting method promotes high efficiency targeted integration in zebrafish embryo somatic tissue.

We next compared the frequency of GeneWeld targeted integration in zebrafish embryos with previous methods that used TALEN targeting of long homology arm (~ 1 kb) donors in combination with restriction enzyme digestion to liberate the linear donor template (Hoshijima et al., 2016; Shin et al., 2014). Targeted integration of pGTag 24 and 48 bp homology arm 2A-TagRFP-CAAX-SV40 donors into exon 2 of *connexin43.4* (*cx43.4*) resulted in 38-56% and 29-47% of injected embryos showing broad RFP expression throughout the nervous system and vasculature (Fig. 2d, f). Increasing the length of the 5’ and 3’ homology arms to 1 kb did not significantly change the frequency of targeted integration compared to 24 bp (*p*=0.1693) or 48 bp (*p*=0.6520) (Fig. 2 f), with 26-47% of injected embryos showing the expected neuronal and vascular RFP expression pattern (Table S1, S2). Injection without the UgRNA leaves the 1 kb homology circular donor intact and reduced targeting to 0-3% (Fig. 2f Circular HR 1 kb; *p*=0.0067; Table S1, S2), as expected given the low frequency of homologous recombination in embryos. In comparison to the GeneWeld method, the frequency of RFP expressing embryos after injection of linear 1 kb homology arm donor template was significantly reduced to 2-9% (Fig. 2f Linear HR 1kb; *p*=0.0111; Table S1, S2). Together, these results suggest long regions of homology in the donor template do not enhance the frequency of integration at the genomic target site, compared with short 24 or 48 bp homology, when using the GeneWeld approach. The increased efficiency compared to linear donor injection may be that introduction of linear DNA into the embryo normally leads to concatemers through NHEJ. We propose that simultaneous targeting of DSBs at the genomic site and in the donor prevents early concatemer formation and favors homology directed repair.

The effect of homology length on GeneWeld integration efficiency was also tested with the 2A-Gal4VP16-β−actin3’UTR donor targeting *esama* exon 2. The *esama* gene is expressed primarily in the vascular system. Increasing homology length from 24 bp to 1 kb dramatically increased the percentage of RFP positive embryos, from 20-23% to 82-94% (*p*=0.0001 Fig. S5, Table S1, S2), but the majority of RFP was not vascular specific, suggesting off-target expression. A high frequency of RFP-positive embryos was also observed when the donor template was injected without UgRNA (27-53%) (Fig. S5 Circular HR 1kb) or the donor template was linearized before injection (83-94%) (Fig. S5 Linear HR 1 kb). Common repetitive elements, enhancers, or a cryptic promoter in the intronic sequence of the *esama* 1 kb homology arms together may lead to off target integration and ectopic RFP expression. These results underscore the utility of the GeneWeld short homology approach for simple donor vector construction and efficient precision targeted integration.

### Efficient germline transmission of GeneWeld precision targeted integration events

Three out of five (60%) *noto*-2A-TagRFP-CAAX-SV40 injected founder fish raised to adulthood transmitted *noto-2A-TagRFP-CAAX-SV40* tagged alleles through the germline (Fig. 3, Table 1, S3 and S4). Although RFP expression in *tyr*-2A-Gal4VP16-β−actin injected *Tg*(*UAS:mRFP*)^tpl2^ embryos was not uniform throughout melanophores, three out of eight RFP mosaic embryos raised to adulthood transmitted germline tagged alleles (37.5%) (Fig. 3, Table 1, S3 and S4). Similarly, for *esama-2A-Gal4VP16-β-actin*, 12/18 (66.7%) F0s displaying widespread vasculature RFP expression were raised to adulthood transmitted *esama-2A-Gal4VP16-β-actin* alleles to the F1 generation. We extended the germline transmission analysis to include 6 additional loci: *flna*, two target sites in *msna* (exon 2 and 6), *aqp1a1*, *aqp8a1*, and *anxa2a*. Overall, the data reveal a combined transmission rate to the F1 generation of 49% across all GeneWeld targeted loci (Fig. 3, Table 1, S3 and S4). Taken together, these results demonstrate the GeneWeld method promotes targeted integration that is efficiently transmitted through the germline in zebrafish.

**Figure 3.**
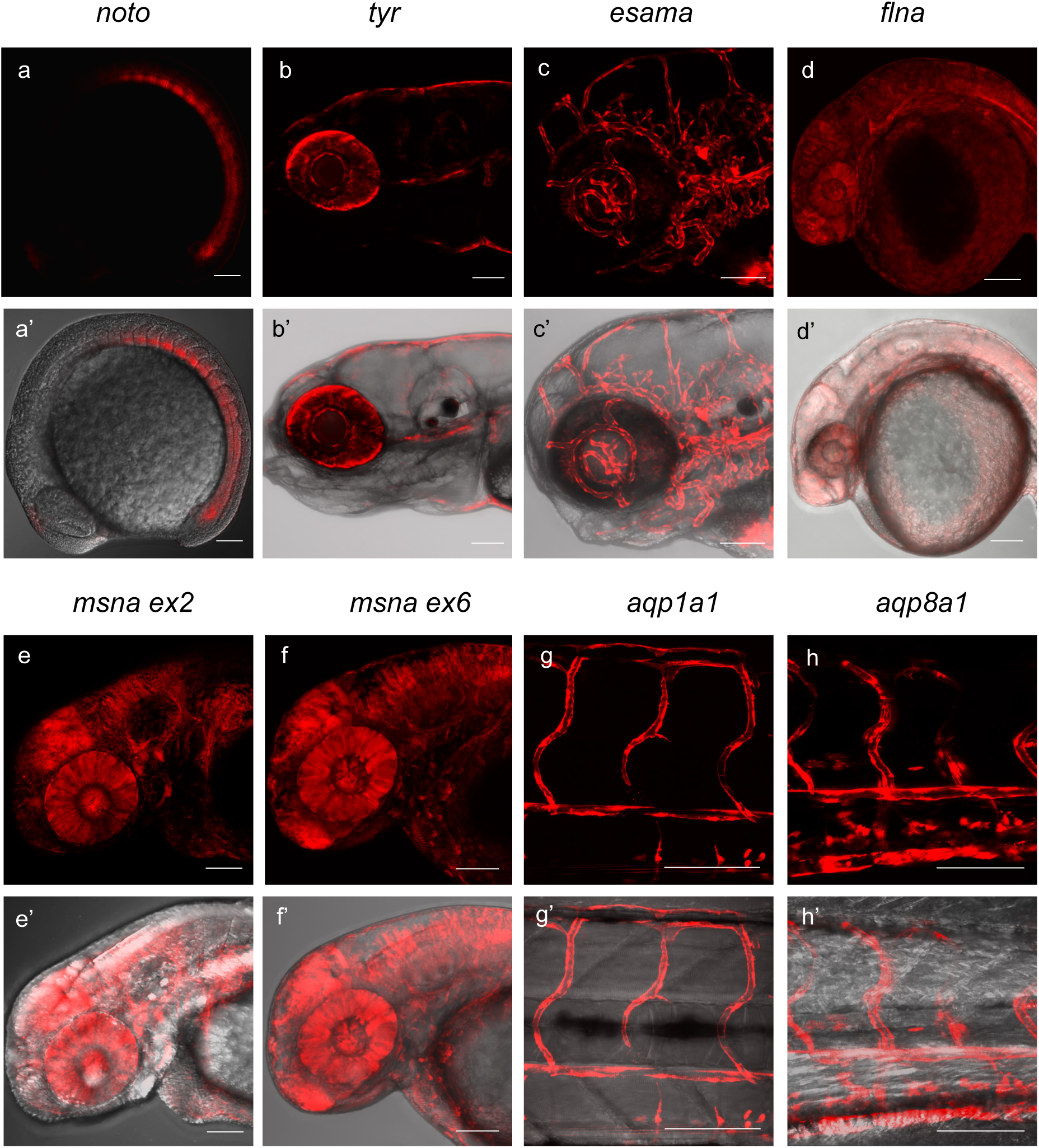
Live confocal images of F1 zebrafish with inherited germline alleles of integrated GTag reporters. (a, a’) *Tg*(*noto-2A-TagRFP-CAAX-SV40)* embryo at mid somite stage showing expression in the notochord and floor plate. (b, b’) *Tg*(*tyr-2A-Gal4Vp16-β-actin)* displaying expression in the melanocytes in a 5 dpf larva. (c, c’) *Tg*(*esama-2A-Gal4Vp16- β- actin)* larva showing expression in the vascular system at 4 dpf. (d, d’) *Tg*(*flna-2A-Gal4VP16- β- actin)* embryo at 1 dpf showing widespread expression. (e, e’ and f, f’) *Tg(msna-2A-Gal4VP16- β-actin)* targeted to either exon 2 or exon 6 showed expression in the central nervous system and vasculature at 2 dpf. (g, g’ and h, h’) *Tg(aqp1a1-2A-Gal4VP16- β-actin)* and *Tg(aqp8a1-2A-Gal4VP16- β-actin)* display expression in the trunk and tail vasculature at 2 dpf. All images are lateral views, and the *Gal4VP16* integrations have *Tg(UAS:mRFP*)^tpl2^ in the background for visualization of expression. Scale bars are 100 μm.

### Precise 5’ and 3’ junctions and single copy integration in F1 germline GTag alleles

We performed Genomic Southern blot analyses and PCR junction fragment sequencing of F1 GTag germline alleles to determine whether GeneWeld targeting lead to precise integration at the 5’ and 3’ sides of the genomic target site. Southern blot analysis and sequencing of *tyr-*2A-Gal4VP16-β-actin F1 progeny demonstrated a single copy integration of the Gal4VP16 cassette (Fig. 4 a-c) with precise sequence at both 5’ and 3’ ends of the integration site (Fig. S6). Analysis of four F1 progeny from two *noto*-2A-TagRFP-CAAX-SV40 founder adults confirmed a single copy integration in *noto* exon 1 (Fig. 4, d-f). However, sequencing of PCR junction fragments in F1 progeny revealed precise 5’ integration but imprecise integration at the 3’ ends that could represent NHEJ repair (Fig. S6).

**Figure 4.**
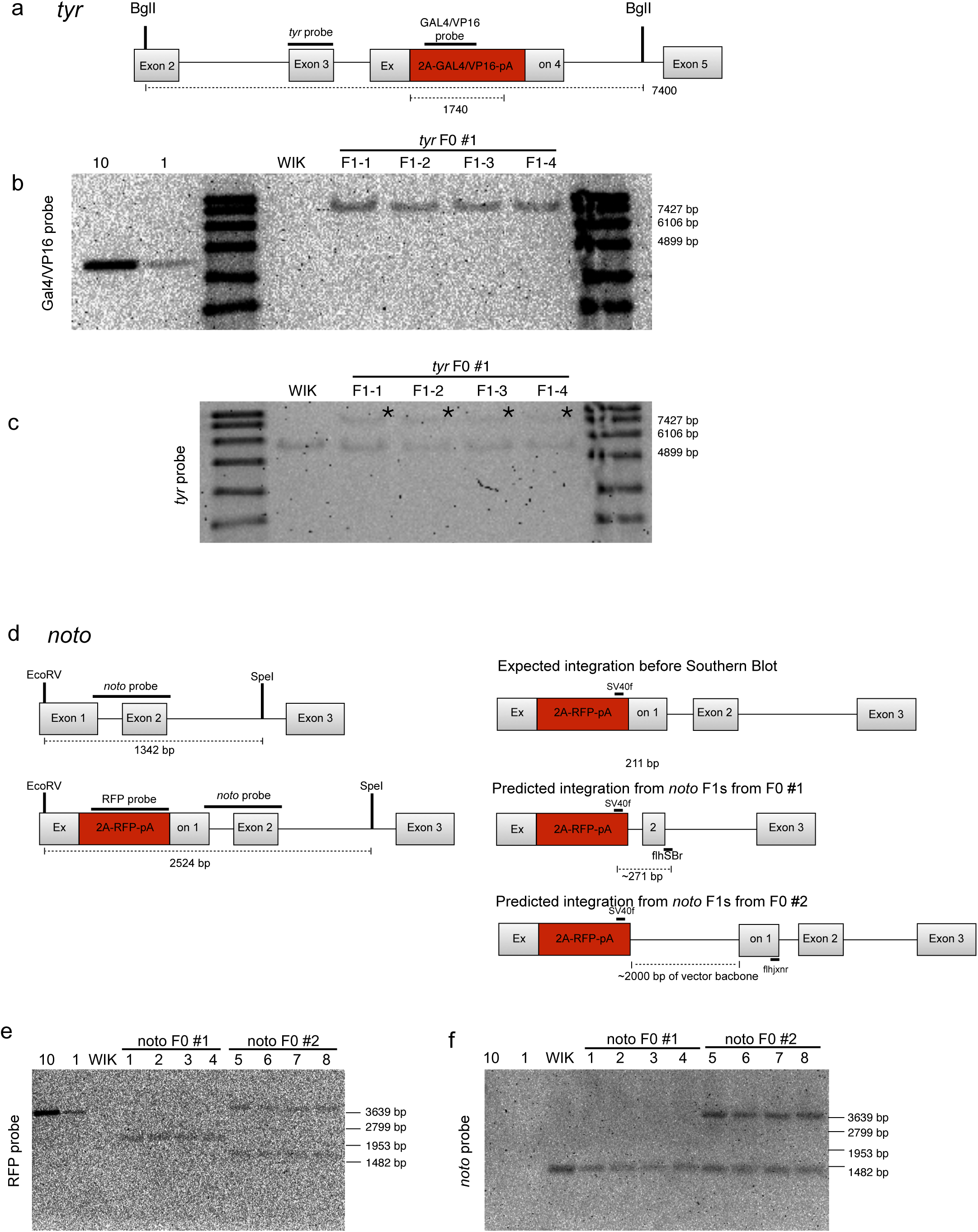
Molecular analysis of F1 GeneWeld GTag targeted alleles at *tyr* and *noto*. (a-c) Molecular analysis of *Tg(tyr-2A-GAL4/VP16)* F1 offspring from a single targeted F0 founder. (a) Schematic of expected integration pattern for *tyr* targeted with pGTag-2A-GAL4/VP16. 148 bp *tyr* probe in Exon 3 and 583bp probe in GAL4/VP16 are indicated. (b) GAL4/VP16 and (c) *tyr* probed Southern blots of genomic DNA from wild type (WIK) and 4 individual GAL4/VP16 positive F1s. The expected 7400 bp band is detected with both probes, suggesting a single copy integration. (d-f) *Tg(noto-2A-RFP)* F1 targeted integration alleles from 2 independent F0 founders. (d) *noto* gene model with location of restriction enzymes used for genomic Southern blot analysis. Location of the 513 bp *noto* probe is indicated (dark lines). The predicted and recovered alleles are shown. (e) Southern blots of F1 *Tg(noto-2A-RFP)* individuals hybridized with RFP probe. F1 from founder F0#1 contain a ~2100 bp band corresponding to integration plus deletion of ~400 bp in *noto*. F1 progeny from founder F0#2 show two bands: a ~3700 bp band corresponding to integration of the reporter plus 2000 bp of vector backbone, and a ~1500 bp band which may represent an off-target integration. Loading controls (10, 1) correspond to 10 copies or 1 copy of RFP containing plasmid. WIK, wild type control DNA. (f) Southern blot in (d) stripped and re-hybridized with the *noto*-specific probe. A 1342 bp band representing the wild type allele was detected in all individuals. The integration allele in F1s from F0 #1 was not detected due to deletion of the region containing the probe. F1s from F0 #2 contain the ~3700 bp band corresponding to the *noto-2A-RFP* integration allele.

Junction fragment analysis of F1s alleles from 5 additional targeted sites in *esama*, *flna*, *msna*, *aqp1a1*, and *aqp8a1* revealed precise events were primarily recovered at the 5’ for all genes examined (30/31 or 97% across seven genes) (Fig. S6). This result is expected, since screening for fluorescent reporter expression from the integrated donor cargo selects for precise 5’ integration. For *esama*, the 3’ junctions were also precise in 9/10 of the F1s examined from 6 different F0s, and both *aqp1a1* and *app8a1* had precise 3’ junctions. This is compared to *msna* E2 targeting with 2A-Gal4VP16-β-actin, where only one out of the 12 F1s examined had a precise 3’ junction. Together, these results indicate that GeneWeld reagents can promote with high frequency precise single copy integration at a genomic cut site without insertion of donor vector backbone sequences, although events involving NHEJ at the at the 3’ end are also recovered.

### Homology Engineered to Distal Genomic gRNA Sites Seeds Deletion Tagging in Somatic Tissue

To further demonstrate the utility of GeneWeld targeted integration, we tested whether the pGTag donor could function to bridge two CRISPR/Cas9 genomic cuts, resulting in simultaneous deletion of endogenous sequences and integration of exogenous DNA to create a “deletion tagged” allele. The pGTag-2A-Gal4VP16 donor was cloned with 48bp homology arms to two gRNA target sites in the zebrafish *retinoblastoma1* (*rb1*) gene. Guide RNAs were designed to sites in exons 2 and 4, which are located 394 bp apart, or exons 2 and 25 which are separated by ~48.4 kb (Fig. 5a). The 5’ homology arm contained sequence upstream of the cut site in exon 2, while the 3’ homology arm contained sequence downstream of the cut site in either exon 4 or exon 25. GeneWeld targeting with the corresponding exon 2-4 or exon 2-25 pGTag-2A-Gal4VP16-β-actin donor into Tg(*UAS:mRFP*)^tpl2^ embryos resulted in injected embryos showing broad and ubiquitous RFP expression (Fig. 5b-c). Using the same approach, we targeted the zebrafish gene *moesina* (*msna*) at exons 2 and 6, located 7.8 kb apart, with 2A-Gal4VP16 using 48 bp of homology (Fig. 5d), and found RFP expression in a pattern consistent with the expression of *msna* (Fig. 5 e, e’). The frequency of RFP positive embryos was similar after targeting *rb1* exon 2-4 (44-78%) and *msna* exon 2-6 (50-85%), and did not seem to be affected by increasing the size of the deleted region from 394 bp to 48.4 Kb in rb1 exon 2-25 (49-70%) (Fig. 5f). Somatic junction fragment analysis detected precise integration of 2A-Gal4VP16 in both genes at the 5’ upstream exon (*rb1* 97%; *msna* 85%) and 3’ downstream exon (*rb1* 67%; *msna* 45%) (Fig. S7). However, only one out of 16 (6%) *rb1* e2-25-2A-Gal4VP16 targeted F0 founders transmitted a precise 5’ junction through the germline, but the 3’ junction could not be amplified by PCR (Table S3, S4). None of the 10 *msna* e2-e6 2A-Gal4VP16 targeted F0 zebrafish transmitted a deletion tagged allele to the next generation. In contrast, targeting 2A-Gal4VP16-β−actin to exon 2 or 6 alone resulted in 2 out of 7 F0s transmitting a targeted allele to the next generation (Table S3, S4).

**Figure 5.**
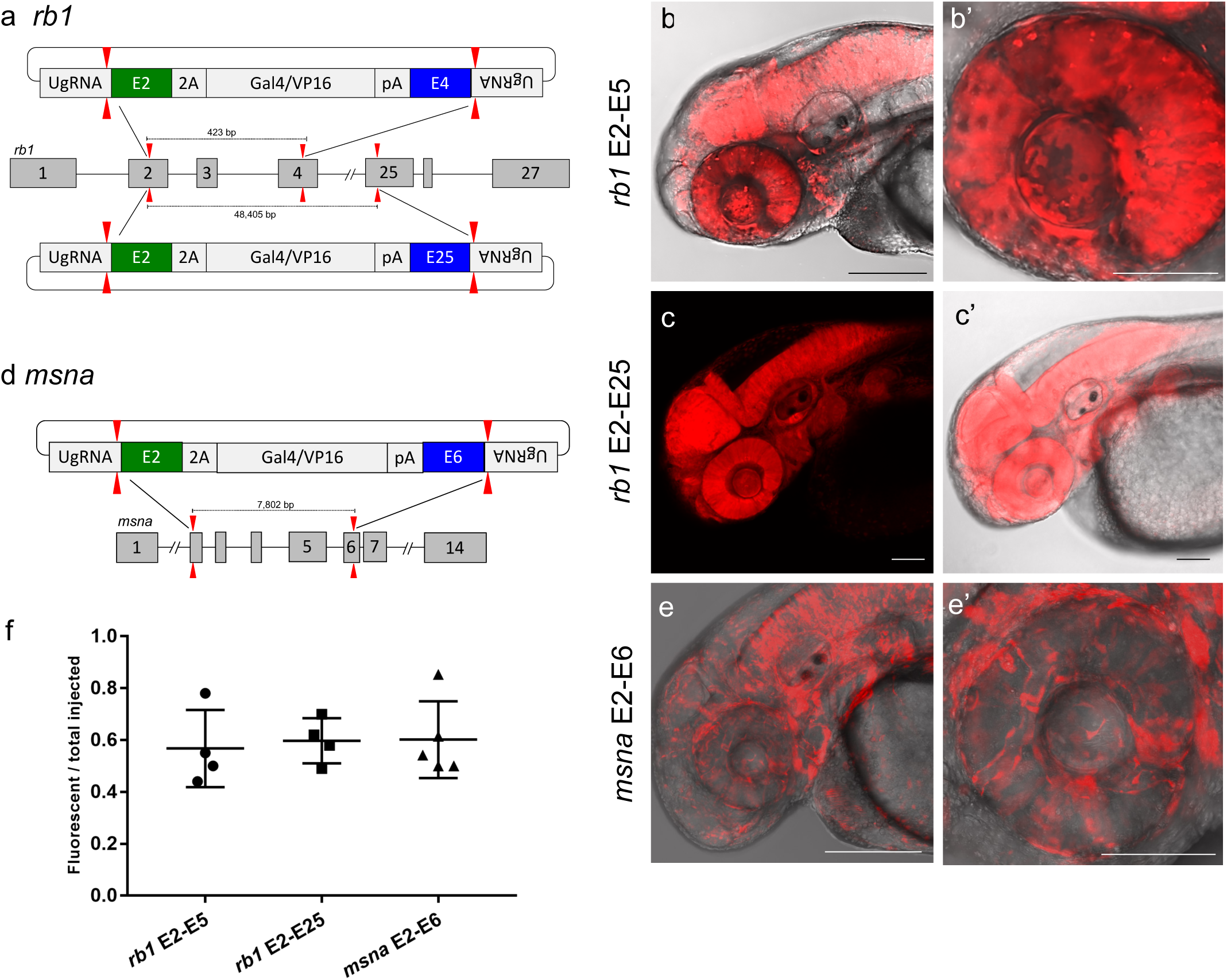
Deletion tagged alleles created with the GeneWeld strategy in zebrafish somatic tissue. (a) Schematic for *Gal4VP16* reporter integration to tag a deletion allele of *rb1* exons 2-4 (top) and *rb1* exons 2-25 (bottom). Arrowheads designate CRISPR/Cas9 DSBs. CRISPR gRNAs in two exons are expected to excise the intervening genomic DNA. The targeting vector contains a 5’ homology arm flanking the upstream exon target site and a 3’ homology arm flanking the downstream exon target site. (b-c) Live confocal image of F0 Tg(*UAS:mRFP*)^tpl2^ embryo after *2A-Gal4VP16* deletion tagging *rb1* exons 2-4 (b, b’) and *rb1* exons 2-25 (c, c’). (d) Schematic for *2A-Gal4VP16* deletion tagging of *msna* exons 2-6. (e, e’) Live confocal image of F0 Tg(*UAS:mRFP*)^tpl2^ embryo after *2A-Gal4VP16* deletion tagging at *msna* exons 2-6. (f) Somatic reporter efficiency of targeted deletion tagging using 48 bp homology arms for *rb1* exons 2-4, *rb1* exons 2-25, and *msna* exons 2-6. Data represents mean +/− s.e.m. of 4 (*rb1*) and 5 (*msna*) independent targeting experiments. Scale bars 200 μm (b, c, c’, e); 100 μm (b’, e’).

Together, these results demonstrate simultaneous targeting of two distal genomic cut sites can create precise HMEJ integration at both ends of a pGTag reporter cassette in somatic tissue, but these events are not easily passed through the germline. This was reinforced by attempting deletion tagging at additional loci, including *kdrl*, *s1pr1*, and *vegfaa*, which showed 32-81% expression in F0 embryos, but no germline transmission to the F1 generation (Table S2, S3).

### Integration of Exogenous DNA Using HMEJ in Porcine and Human Cells is More Efficient than HR

To determine if HMEJ integration directed by short homology functions efficiently in large animal systems, we tested the GeneWeld targeting strategy in *S. scrofa* fibroblasts. The *ROSA26* safe harbor locus was targeted with a cassette that drives ubiquitous eGFP expression from the UbC promoter (Fig. 6a-c). GeneWeld reagents, where the genomic sgRNA was replaced with mRNAs encoding a TALEN pair to generate a genomic DSB in the first intron of *ROSA26*, were delivered to pig fibroblasts by electroporation. This strategy was compared to cells electroporated with just the TALEN pair and a HR donor containing approximately 750 bp of homology flanking the genomic target site. GFP expression was observed in 23% of colonies using GeneWeld reagents, compared to 2% of colonies using the HR donor with ~750 bp homology arms. Co-occurring precise 5’ and 3’ junctions were observed in over 50% of the GFP+, GeneWeld engineered colonies while none of the GFP+, HR colonies contained both junctions. Sequencing of junctions from 8 GFP+, GeneWeld engineered colonies that were positive for both junctions showed precise integration in 7/8 colonies at the 5’ junction and 8/8 colonies at the 3’ junction.

**Figure 6.**
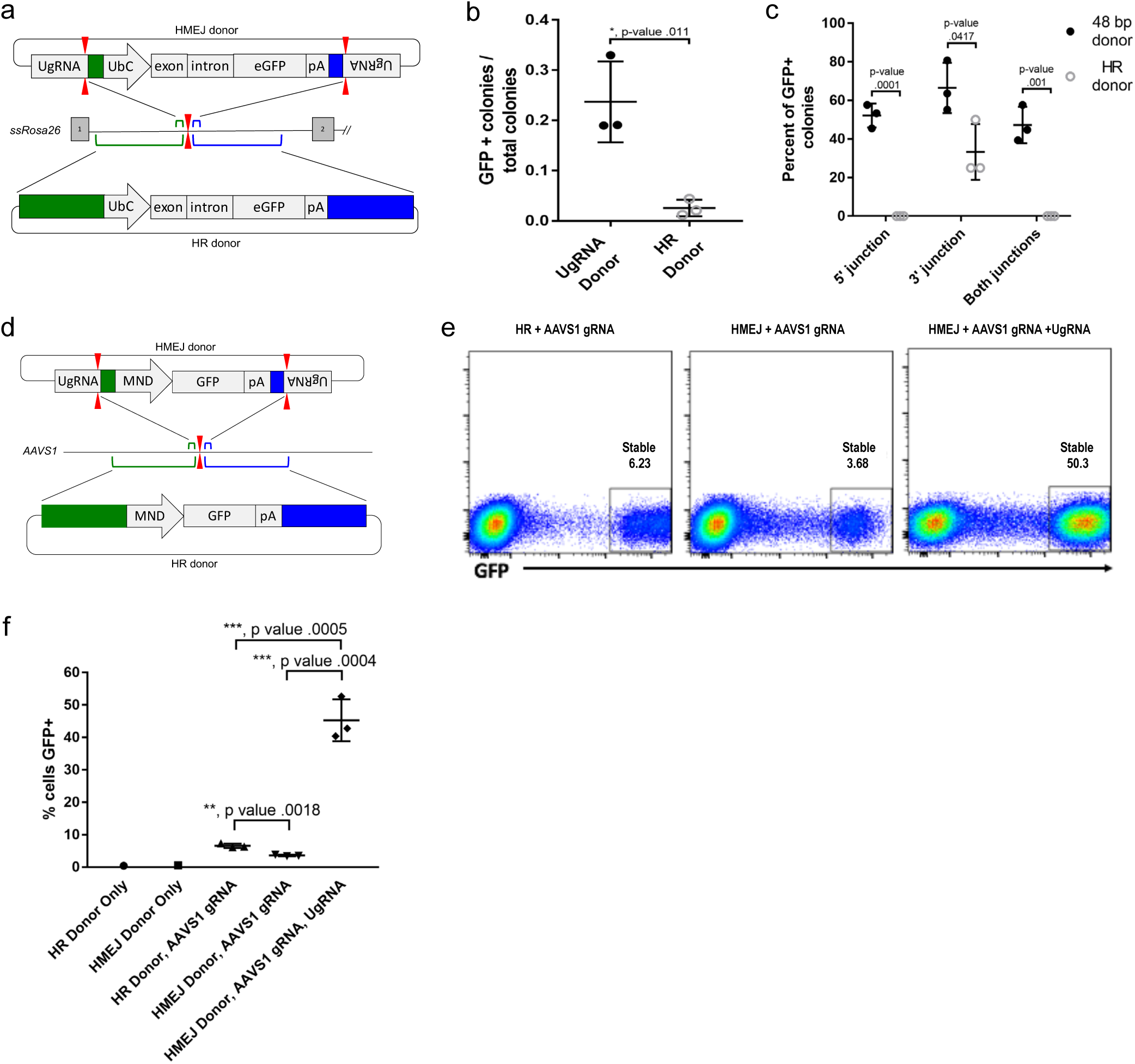
HMEJ-based targeted integration with UgRNA-based vectors promotes efficient knock-in in porcine fibroblasts and human K-562 cells. (a) Strategy for integration using HMEJ and HR donors into intron 1 of *S. scrofa ROSA26* locus. Arrowheads CRISPR/Cas9 (for HMEJ donor) and TALEN (genome) DSBs. (b) Targeting efficiency of the HMEJ donor vs the HR donor as reported by GFP positive colonies out of total colonies. (c) Percent of GFP positive colonies analyzed containing properly sized junction fragments, comparing HMEJ and HR donors. Data are from three independently targeted cell populations. Data represents mean +/− s.e.m. of 3 independent targeting experiments. (d) Diagram of HR and HMEJ strategies for targeted integration of a MND:GFP reporter cassette into the human *AAVS1* locus. (e) Flow cytometry analysis of GFP expression 14 days post-electroporation for each targeting modality: HR (left), HMEJ without universal sgRNA (middle), and HMEJ with universal sgRNA (right). Stable gate was drawn to measure the uniformly expressing population formed by targeted integration and was set based on episome only controls. (f) Quantitation of stable GFP expressing population as measured by flow cytometry at day 14. Data are from three independently targeted cell populations. Data represents mean +/− s.e.m. of 3 independent targeting experiments. p values calculated using two-tailed unpaired t-test.

The GeneWeld strategy was also used to target integration of a MND:GFP reporter (Halene et al., 1999) into the *AAVS1* safe harbor locus in human K-562 cells (Fig. 6d-f). Integrations were attempted with either GeneWeld reagents or an HR donor targeting the *AAVS1* cut site by electroporation of K-562 cells. Cells were FACs sorted by GFP at day 14 following electroporation. With GeneWeld reagents over 50% of cells were GFP positive, compared to only 6% of cells electroporated with the HR donor. This suggests the GeneWeld strategy promoted efficient integration and stable expression of the MND:GFP cassette at the *AAVS1* locus (Fig. S8). Expression was maintained over 50 days, and 5’ precise junction fragments were observed following PCR amplification in bulk cell populations (Fig. S9). The results above demonstrate that the GeneWeld strategy outperforms traditional HR techniques in mammalian cell systems and is effective without antibiotic selection.

## Discussion

The results described here demonstrate the utility of short homology-based gene targeting for engineering precise integration of exogenous DNA and expand the potential of efficient tagging to diverse loci with differing endogenous expression levels. Key to our strategy was to observe reporter gene expression that correlated with precise integration (Fig. 2 and Fig. S1) and selecting these animals for examination of germline transmission to the F1 generation. However, we have observed that restricted patterns of gene expression such as *noto* do not necessary correlate with the degree of precise integration throughout the embryo following injection (data not shown). Moreover, we show that using short homology to bridge distal ends together simultaneously creates a deletion and a reporter integration, however, these events are not easily passed through the germline. This emphasizes that the events observed in the somatic tissue of the zebrafish embryo following injection do not necessarily predict transmission through the germline. We demonstrate efficient integration of cargos up to approximately 2 kb in length in zebrafish, pig fibroblasts, and human cells, and in zebrafish, these events can be passed through the germline in nearly 50% of the injected embryos. Both CRISPR/Cas9 and TALENs are effective as GeneWeld genomic editors, providing flexibility in deployment and genome-wide accessibility.

Several components of the GeneWeld strategy may lead to enhanced somatic and germline targeting efficiencies in zebrafish as compared to previous reports (Hisano et al., 2015). Canonical NHEJ is highly active during rapid cell divisions in early zebrafish embryogenesis (Bedell et al., 2012). However, given the correct sequence context surrounding the dsDNA break, MMEJ is the preferred method of non-conservative repair (Ata et al., 2018; He et al., 2015; Kent et al., 2015). GeneWeld homology arms are rationally designed based on the known homology searching activity of RAD51 and strand annealing activity of RAD52 (Conway et al., 2004; Singleton et al., 2002). In our experiments at *noto*, gene targeting is significantly reduced when 48 bp homology arms are altered by 1 bp to 47 or 49 bp (Fig. S3). This suggests that optimal short homology arms should be designed in groups of 3 and/or 4 bp increments. We are currently testing this hypothesis further. Additionally, Shin et al., 2014 showed the highest rates of somatic targeting when their donor was linearized *in vitro* inside a ~1 kb 5’ homology arm, leaving 238 bp of homology flanking the knock-in cargo. Thus, it is tempting to speculate that gene targeting in these experiments proceeded not through HR, but through other related HMEJ DNA repair pathway more similar to the findings presented here.

Using GeneWeld reagents we observed that the length of the homology arm does not influence the observed somatic frequency of reporter gene expression when examined between 24 bp and 1 kb in length. However, there are distinct disadvantages to using long homology arms in experimental design and execution. First, longer homology arms often require the use of intronic sequences which are highly enriched of repetitive sequences that can potentially mediate random integrations across the genome. Second, alternative promoters and enhancer sequences present in intronic regions can also affecting the efficiency of reporter gene expression, as we likely observed with the *esama* gene. Lastly, cloning of the homology arms can be arduous, since intronic sequences are highly variable across individuals, and therefore, it is recommended that the longer homology arms are cloned from individuals used for gene targeting. Taken together, we suggest that using the GeneWeld method with short homology arms is a preferable choice for efficient precise exogenous DNA integration.

The dramatic shift of DNA repair at genomic DSBs from cis-NHEJ to trans-HMEJ using GeneWeld donors likely also influences enhanced editing of the germline. Across all zebrafish experiments with germline transmission, 49% of founders transmitted tagged alleles, with 17.4% of gametes carrying the edited allele of interest (Table S3, S4), demonstrating decreased germline mosaicism and increased germline transmission from previous reports. Given that our somatic knock-in and germline transmission rates are higher than most published reports, we conclude that GeneWeld is a more effective homology-based method for generating precisely targeted knock-in alleles in zebrafish.

While targeting *noto* with 5’ only homology shows an increase in targeting efficiency with longer homology, increasing homologies on both ends of the cargo DNA did not increase targeting efficiency (Fig. 2e). Positive events are selected only by fluorescently tagged alleles, indicating precise 5’ integration patterns. We speculate that inclusion of homology at the 3’ end of our cargo creates competition for the donor DNA ends, as not all editing events are precise at both 5’ and 3’ junctions (Fig. 4 and Fig. S4). Thus, it is conceivable that precise events at the 3’ end could preclude precise integration at the 5’ end during some editing events, and vice versa. It is tempting to speculate that this data hints at synthesis dependent strand annealing (SDSA) as a possible DNA repair mechanism for pGTag donor integration (Ceccaldi et al., 2016). After strand invasion using either of the homology domains and replication through the reporter, second DNA end capture may abort before or after replication through the opposing homology domain, resulting in imprecision, as greater than or equal to 150 bp is required for proper second end capture in yeast (Mehta et al., 2017). Experiments to address this hypothesis by varying homology arm lengths flanking the donor cassettes and including negative selection markers are of note for future work in determining the genetic mechanisms that promote efficient integration.

Timing and turnover of Cas9 during the genomic editing event can influence cut efficiency and somatic mosaicism/germline transmission rates (Clarke et al., 2018; Zhang et al., 2018), increasing the interest of using RNP during all precision gene editing applications. However, we were unable to observe fluorescence following injection of GeneWeld components with RNPs or detect targeted integrations at a high frequency (unpublished data). We hypothesize this is due to Cas9 and UgRNA locating and binding to the UgRNA sites on the pGTag donors during dilution of the injection mixture. This heteroduplex either activates DSBs on the donor *in vitro*, or directly after injection, before the genomic gRNA can locate and cut the genome. Thus, the stochastics of DNA end availability are altered using RNPs and integration activity is greatly reduced. Injection of the GeneWeld plasmid donor and RNPs in separate injection mixtures does not produce integration in zebrafish embryos (unpublished data). Further experiments could address these limitations through the use of inducible nuclease systems.

Targeting genes with lethal phenotypes, such as tumor suppressors or other genes required for embryogenesis, is of interest to the zebrafish community. However, using fluorescence to screen for targeted events can be misleading. For example, the RFP signal is dramatically reduced or lost upon biallelic inactivation of *noto*, likely when notochord cells transfate to muscle cells (Melby et al., 1996; Talbot et al., 1995). Additionally, though deletion tagging using two target sites in the genome seems to be robust in somatic tissue (Fig. 5), germline transmission of deletion tags is rare. This suggests that edited germ cells may be lost to apoptosis due to the additional cut in the genome, or that heterozygous deletion tagged alleles are recognized during homologous chromosome pairing and are repaired or lost as germ cells mature. Similar susceptibility of stem cells to apoptosis following gene editing has been previously observed (Ihry et al., 2018; Li et al., 2018). In both of these cases, it may be necessary to modulate GeneWeld reagent concentrations in order to avoid biallelic inactivation of the genomic target, or to ensure homozygous deletion tagging.

Amplification of the fluorescence signal using GAL4/VP16 allowed us to target several genes for which we did not observe a fluorescence report from integration of a fluorescent protein directly. While this approach is advantageous for selecting correctly targeted embryos to examine for germline integration, GAL4/VP16 may have toxic effects as reported previously (Ogura et al., 2009). For example, we found dominant phenotypes in the F1 generation for both *msna* and *flna* which could reflect toxicity from high levels of expression of GAL4/VP16. Alternatively, these gene could also display haploinsufficiency or express a partial protein product that functions in a dominant manner. Heterozygous *msna* mutants targeting exon 5 in the F1 generation display phenotypes similar to morpholino targeting of this gene (Wang et al., 2010) (data not shown), suggesting haploinsufficiency or a dominant negative peptide is a likely explanation.

GeneWeld is also an effective strategy to precisely control exogenous DNA integration in mammalian cell lines. While our data shows an approximate 10-fold increase in targeted integration using 48 bp of homology to drive HMEJ versus HR, Zhang et al. (2017) concluded that targeted integration did not appreciably increase until homology arms of ~600 bp were used (Zhang et al., 2017). However, this could reflect differences in the experimental design or cell types used and suggest different DNA repair pathways may be more prevalent in certain conditions. Deciphering the DNA repair pathway used for HMEJ in zebrafish and mammalian cells is paramount to increasing editing efficiencies in basic research and for gene therapy.

Given the high efficiency and precision of GeneWeld, additional applications to efficiently introduce other gene modifications, such as single or multiple nucleotide polymorphisms, by exon or gene replacement is possible using the deletion tagging method. Further, GeneWeld could be used to create conditional alleles by targeting conditional gene break systems into introns (Clark et al., 2011). In conclusion, our suite of donor vectors with validated integration efficiencies, methods, and web interface for pGTag donor engineering will serve to streamline experimental design and broaden the use of designer nucleases for homology-based gene editing at CRISPR/Cas9 and TALEN cut sites in zebrafish. We also demonstrate an advanced strategy for homology-based gene editing at CRISPR/Cas9 and TALEN cut sites in mammalian cell lines. Our results open the door for more advanced genome edits in animal agriculture and human therapeutics.

## Methods

### Contact for reagent and resource sharing

Further information and requests for resources and reagents should be directed to and will be fulfilled by the Lead Contact, Jeffrey Essner.

### Experimental model and subject details

Zebrafish were maintained in Aquatic Habitats (Pentair) housing on a 14 hour light/10 hour dark cycle. Wild-type WIK were obtained from the Zebrafish International Resource Center. The Tg(*miniTol2/14XUAS:mRFP, γCry:GFP*)^tpl2^, shortened to Tg(*UAS:mRFP*)^tpl2^, was previously described (Balciuniene et al., 2013). All experiments were carried out under approved protocols from Iowa State University IACUC.

The human K-562 chronic myelogenous leukemia cell line (ATCC CCL-243) used in gene targeting experiments was cultured at 37°C in 5% CO_2_ in RPMI-1640 (Thermo Fisher Scientific) supplemented with 10% fetal bovine serum (FBS) and Penicillin/Streptomycin. Electroporation was conducted with 1.5 x 10^5^ cells in a 10 μl tip using the Neon electroporation device (Thermo Fisher Scientific) with the following conditions: 1450V, 10ms, 3x pulse. Nucleic acid dosages were as follows: 1.5 μg Cas9 mRNA (Trilink Biotechnologies), 1 μg each chemically modified sgRNA (Synthego), and 1 μg donor plasmid.

Fibroblasts were cultured in DMEM (high glucose) supplemented to 10% vol/vol FBS, 20 mM L-glutamine and 1X Pen/ Strep solution and transfected using the Neon™ system (Invitrogen). Briefly, 1 x 10^6^ fibroblasts were transfected with 1 ug of polyadenylated ROSA TALEN mRNA, 1 μg of universal gRNA mRNA, 1 μg of polyadenylated Cas9 mRNA and 1 μg of donor plasmid. Transfected cells were cultured for 3 days at 30°C before low density plating, extended culture (10 days) and colony isolation. Individual colonies were aspirated under gentle trypsanization, replated into 96-well plates and cultured for 3-4 days.

### pGTag series vectors

To build the pGTag vector series, 2A-TagRFP, 2A-eGFP, and 2A-Gal4/VP16 cassettes were assembled from a 2A-TagRFP-CAAX construct, p494. To clone the eGFP cassette, the plasmid p494 was amplified with primers F-p494-XhoI and R-p494-SpeI to generate unique enzyme sites in the backbone. The eGFP coding sequence (Clontech Inc.) was amplified with the primers F-eGFP-SpeI and R-eGFP-XhoI to generate the corresponding enzyme sites on the eGFP coding sequence. Fragments were digested with SpeI-HF and XhoI (NEB) and following column purification with the Qiagen miniprep protocol, were ligated to the plasmid backbone with T4 ligase (Fisher).

The Gal4/VP16 coding sequence and zebrafish β-actin 3’ untranslated region was amplified from vector pDB783 (Balciuniene et al., 2013) with primers F-2A-Gal4-BamHI and R-Gal4-NcoI to add a 2A peptide to the 5’ end of the Gav4Vp16 cDNA. The resulting PCR product was then cloned into the intermediate Topo Zero Blunt vector (Invitrogen) and used for mutagenesis PCR with primers F and R ‘-gal4-Ecofix’ to disrupt the internal EcoRI restriction site. The resulting Gal4/VP16 sequence was cloned into the BamHI and NcoI sites in the p494 backbone.

The 5’ universal/optimal guide site and *lac*Z cassette were added to pC-2A-TagRFP-CAAX-SV40, pC-2A-eGFP-SV40, and pC-2A-Gal4VP16-β-actin with the following steps. The *lac*Z was first amplified with primers F-lacZ and R-lacZ, which add the type IIS enzyme sites to either end of the *lac*Z. The resulting PCR product was then cloned into an intermediate vector with the Zero Blunt® TOPO® PCR Cloning Kit (Invitrogen). This intermediate was used as a template in a nested PCR to add the Universal guide sequence GGGAGGCGTTCGGGCCACAGCGG to the end of the *lac*Z sequence. The nested PCR used primers F-lacZ-universal-1 and R-lacZ-universal-BamHI to add the first part of the universal guide to one end and a BamHI site to the other. This was used as template for PCR with the primers F-lacZ-universal-EcoRI and R-lacZ-universal-BamHI to add the remainder of the universal guide and an EcoRI site. The fragment was column purified as above, digested with EcoRI-HF and BamHI-HF and cloned into the appropriate sites in the three vectors.

The 3’ universal guide and type 2 restriction enzyme sites were cloned into each vector in two steps. A segment from a Carp beta-actin intron containing a 99 bp spacer flanked by two BspQI sites was amplified using the primers F-3’-uni-1 and R-3’-uni-1 to add the universal site to one side of the spacer. This product was column purified as above and used as template for the second amplification with primers F-3’-uniNco1 and R-3’-uniEagI to add cloning sites. This product was column purified and cloned using the Topo zero blunt kit. This intermediate was digested with NcoI-HF and EagI, and the BspQI fragment purified and cloned into the three vectors as above. Ligations were grown at 30°C to reduce the possibility of recombination between the two universal guide sites.

Correct clones for pU-2A-TagRFP-CAAX-U, pU-2A-eGFP-U, and pU-2A-Gal4/VP16-U were selected and used as template for mutagenesis PCR with KOD to remove extra BspQI sites from the backbone with primers F/R-BBfix, digested with DpnI (NEB), and ligated with T4 ligase. A correct pU-2A-TagRFP-CAAX-U clone was used as template for PCR with F/R-TagRFPfix to interrupt the BspQI site in the TagRFP coding sequence as above. A correct clone of pU-2A-Gal4/VP16-U was selected and used as template with primers F/R-Bactfix to remove the BspQI site in the Beta-actin terminator, the product was re-cloned as above. All constructs were sequence verified.

### Homology arm design and donor vector construction

For detailed methods, see Supplementary gene targeting protocol. In brief, homology arms of specified length directly flanking a genomic targeted double strand break were cloned into the pGTag vector, in between the UgRNA sequence and the cargo. A three nucleotide buffer sequence lacking homology to the genomic target site was engineered between the donor UgRNA PAM and the homology arms, in order to maintain the specified homology arm length. To generate 1kb homology arms for zebrafish genes *cx43.4* and *esama*, ~2kb of genomic DNA surrounding the CRISPR target site was PCR amplified from adult WIK finclips using the proofreading enzyme KOD, and then sequenced to identify polymorphisms. pGTag 1kb homology arm vectors were injected into embryos from adults with the matching genomic sequence. See Table S4 for all homology arms, gRNA target sites, and spacers.

### Zebrafish embryo injection

pT3TS-nCas9n was a gift from Wenbiao Chen (Addgene plasmid #46757). XbaI linearized pT3TS-nCas9n was purified under RNase-free conditions with the Promega PureYield Plasmid Miniprep System. Linear, purified pT3TS-nCas9n was used as template for *in vitro* transcription of capped, polyadenylated mRNA with the Ambion T3TS mMessage mMachine Kit. mRNA was purified using Qiagen miRNeasy Kit. The genomic and universal sgRNAs were generated using cloning free sgRNA synthesis as described in (Varshney et al., 2015) and purified using Qiagen miRNeasy Kit. Donor vector plasmid DNA was purified with the Promega PureYield Plasmid Miniprep System. *noto, cx43.4, tyrosinase, and moesina*, were targeted by co-injection of 150 pg of nCas9n mRNA, 25 pg of genomic sgRNA, 25 pg of UgRNA (when utilized), and 10 pg of donor DNA diluted in RNAse free ddH_2_O. The *rb1* targeting mixture contained 300 pg nCas9n mRNA. 2 nl was delivered to each embryo.

### Recovery of zebrafish germline knock-in alleles

Injected animals were screened for fluorescence reporter expression on a Zeiss Discovery dissection microscope and live images captured on a Zeiss LSM 700 laser scanning confocal microscope. RFP/GFP positive embryos were raised to adulthood and outcrossed to wildtype WIK adults to test for germline transmission of fluorescence in F1 progeny. *tyr, esama*, *rb1* and *msna* embryos targeted with Gal4VP16 were crossed to *Tg*(*UAS:mRFP*)*^tpl2^*.

### DNA isolation and PCR genotyping

Genomic DNA for PCR was extracted by digestion of single embryos in 50mM NaOH at 95°C for 30 minutes and neutralized by addition of 1/10^th^ volume 1M Tris-HCl pH 8.0. Junction fragments were PCR-amplified with primers listed in Table S6 and the products TOPO-TA cloned before sequencing.

### Southern blot analysis

Genomic Southern blot and copy number analysis was performed as described previously (McGrail et al., 2011). PCR primers used for genomic and donor specific probes are listed in Table S6.

### Junction fragment analysis in pig fibroblasts

Individual colonies were scored for GFP expression and prepared for PCR by washing with 1X PBS and resuspension in PCR-safe lysis buffer (10 mM Tris-Cl, pH 8.0; 2 mM EDTA; 2.5% (vol/vol) Tween-20; 2.5% (vol/vol) Triton-X 100; 100 μg/mL Proteinase K followed by incubation at 50°C for 60 min and 95°C for 15 min. PCR was performed using 1X Accustart Supermix (Quanta) with the primers: 5’ junction F-5’ TAGAGTCACCCAAGTCCCGT-3’, R-5’-ACTGATTGGCCGCTTCTCCT-3’; 3’ junction F-5’-GGAGGTGTGGGAGGTTTTT-3’, R-5’-TGATTTCATGACTTGCTGGCT-3’. ROSA TALEN sequences are: TAL FNG NI NI HD HD NG NN NI NG NG HD NG NG NN NN; TAL RHD NN NG NI HD NI HD HD NG NN HD NG HD NI NI NG.

### K-592 Flow Cytometry

K-562 cells were assessed for GFP expression every 7 days for 28 days following electroporation. Flow cytometry was conducted on an LSRII instrument (Becton Dickinson) and data was analyzed using FlowJo software v10 (Becton Dickinson). Dead cells were excluded from analysis by abnormal scatter profile and exclusion based on Sytox Blue viability dye (Thermo Fisher Scientific).

Junction PCR to detect targeted integration was conducted using external genomic primers outside of the 48bp homology region and internal primers complementary to the expression cassette. PCR was conducted using Accuprime HIFI Taq (Thermo Fisher Scientific). PCR products from bulk population were sequenced directly.

### Quantification and statistical analysis

Statistical analysis was performed using GraphPad Prism software. Data plots represent mean +/− s.e.m. of n independent experiments, indicated in the text. *p* values were calculated with two-tailed unpaired *t*-test. Statistical parameters are included in the Figure legends.

### Data and software availability

The webtool ***GTagHD*** was developed to assist users in designing oligonucleotides for targeted integration using the pGTag vector suite. GTagHD guides users through entering: 1) the guide RNA for cutting their cargo-containing plasmid; 2) the guide RNA for cutting their genomic DNA sequence; (3) the genomic DNA sequence, in the form of a GenBank accession number or copy/pasted DNA sequence; and 4) the length of microhomology to be used in integrating the plasmid cargo. If the user is utilizing one of the pGTag series plasmids, GTagHD can also generate a GenBank/ApE formatted file for that plasmid, which includes the user’s incorporated oligonucleotide sequences. GTagHD is freely available online at http://genesculpt.org/gtaghd/ and for download at https://github.com/Dobbs-Lab/GTagHD.

### Key resources table

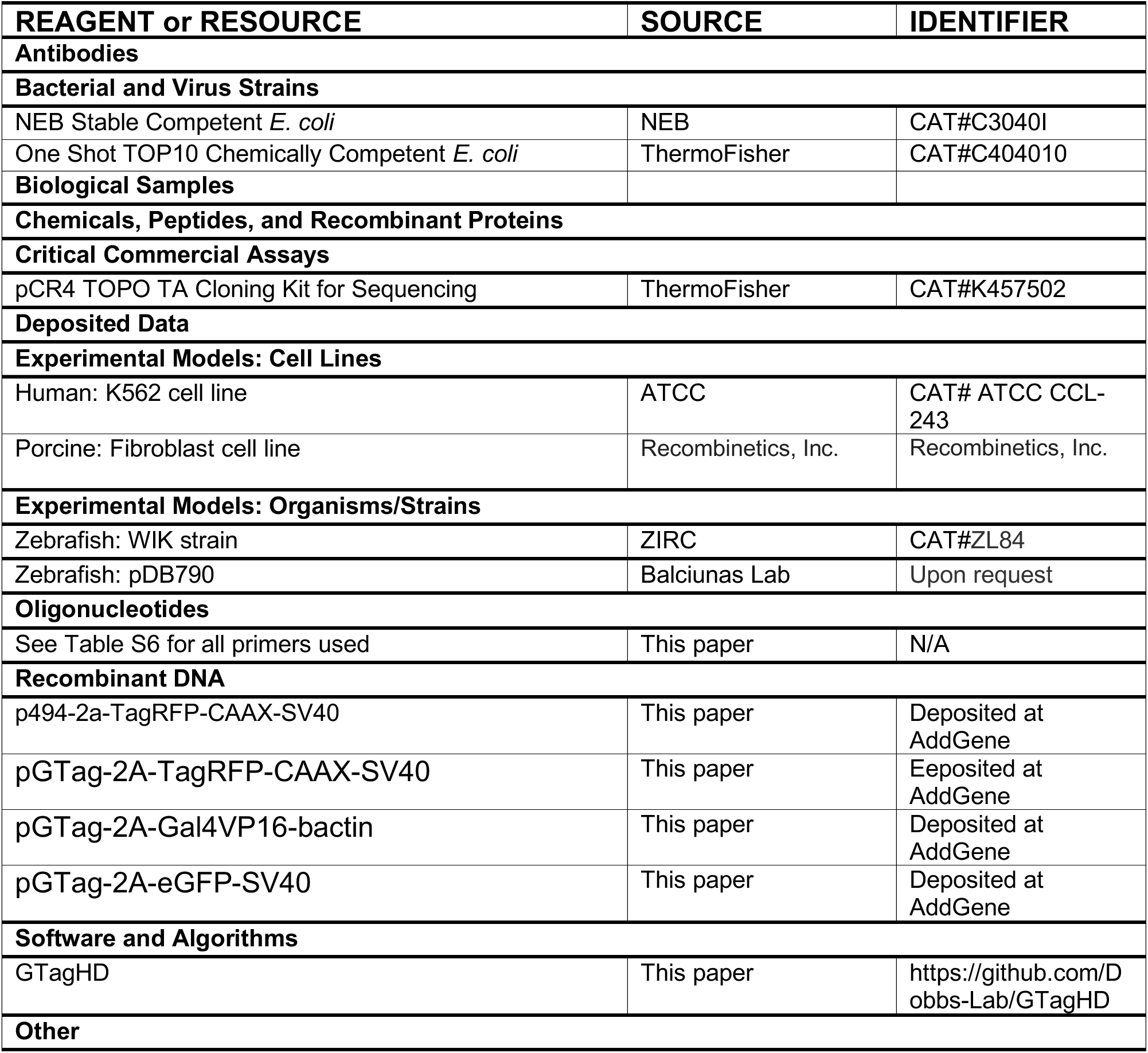

## Acknowledgements

This work was supported by NIH grants R24OD020166 (JJE, MM, DLD, KJC, SCE), GM088424 (JJE), and GM63904 (SCE).

## Author Contributions

WAW, JMW, MM and JJE conceived the study; WAW, JMW, MM and JJE wrote the manuscript with input from SCE and KJC; WAW, JMW, MPA, MET, KCM, ML, ZM, AW and JAH designed and performed the zebrafish experiments; JMW and TJW designed and created the vector suite with input from WAW, KJC, SCE, KMK, C-BC, DB, MM, and JJE; CMM created the web interface with input from JMW, WAW, CSM and DLD; SLS, DAW, MKV and DFC designed and performed the pig fibroblast experiments. BSM and BRW designed and performed the human cell line experiments.

---

**Supplementary Figure 1.**
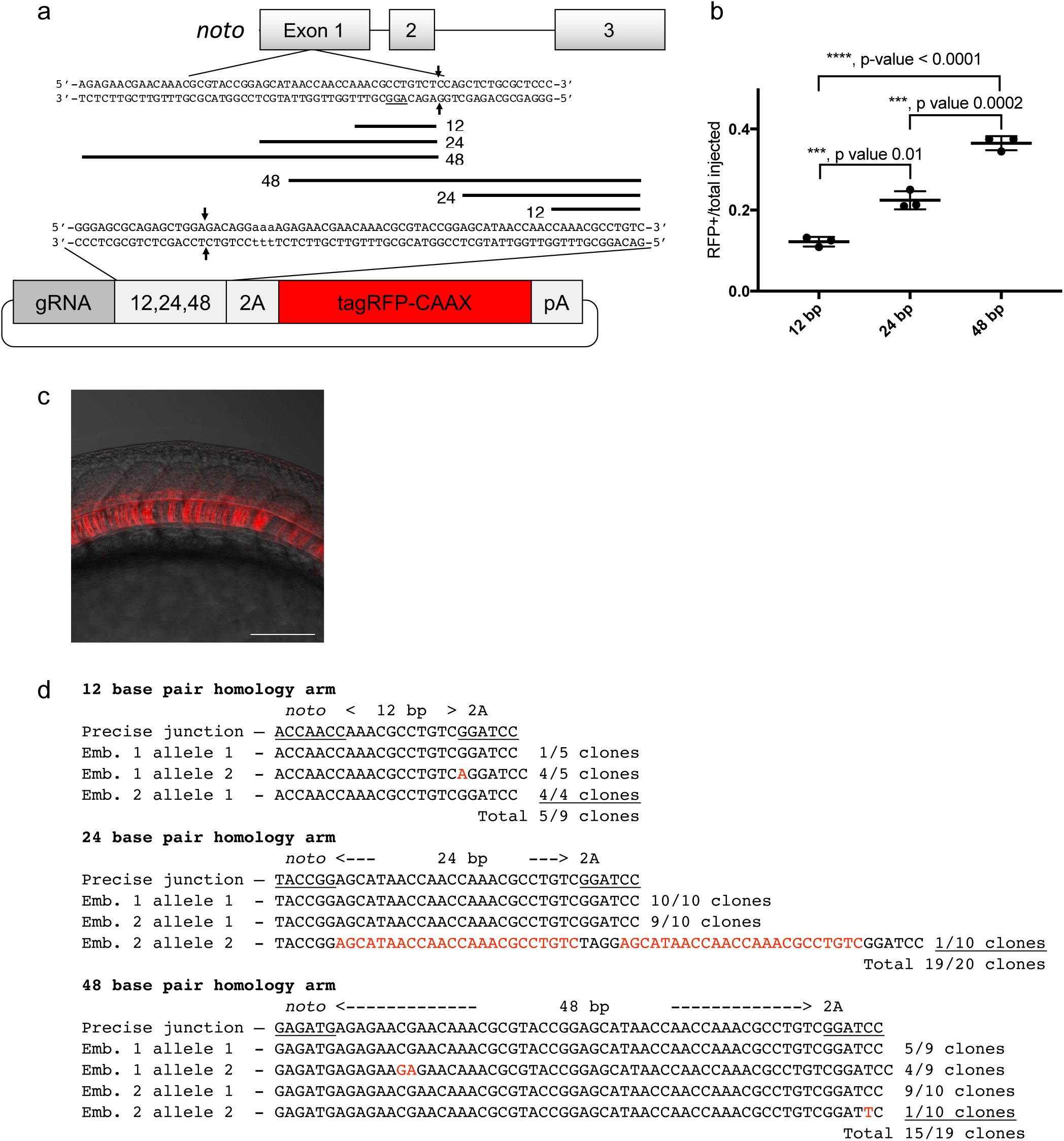
Short homology to the *noto* gene from a single homology arm 5’ to the gRNA target site targets integration in zebrafish embryos. (a) Schematic for *noto* homology arm and donor vector design. gRNA is the *noto* non-coding template strand. Black bars represent 12, 24, and 48 bp homology arms. PAM sequences are underlined. (b) Targeting efficiency of 12, 24, and 48 bp *noto* 5’ only donors. Data represents mean +/− s.e.m. of 3 independent targeting experiments. p values calculated using two-tailed unpaired t-test. (c) Live confocal image of *noto-2A-TagRFP-CAAX-SV40* targeted embryo showing specific RFP expression in the notochord. Scale bar is 100 μm. (d) Sanger sequencing of cloned 5’ junction fragments from RFP positive F0 embryos, aligned to the expected sequence from a precise integration event.

**Supplementary Figure 2.**
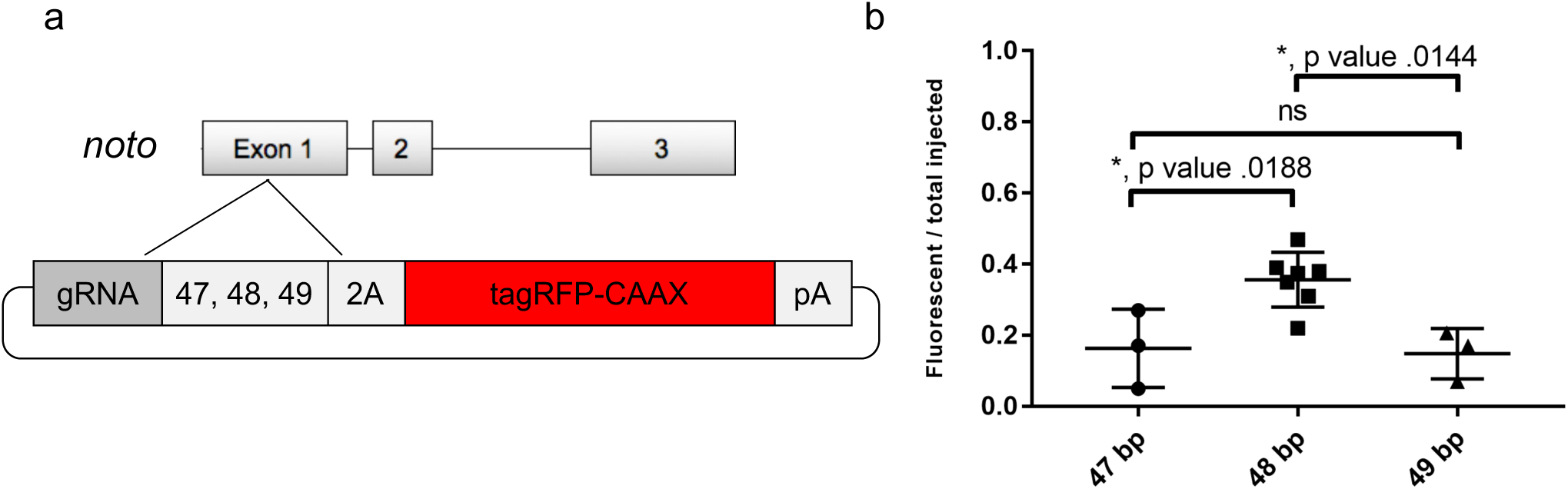
Single base pair differences in homology arm length 5’ to the Cas9/gRNA cut site influence integration frequencies in zebrafish embryos. (a) Schematic for targeting 2A-TagRFP-CAAX-SV40 into *noto* exon 1 with 5’ homology to the Cas9/gRNA cut site containing 47, 48, or 49 bp of homology. (b) The frequency of injected zebrafish embryos displaying notochord RFP expression after targeting *noto* exon 1 with donors containing 47, 48, or 49 bp of 5’ homology. Data represents mean +/− s.e.m. of 3 (47 bp, 49 bp) or 7(48 bp) independent targeting experiments. p values calculated using two-tailed unpaired t-test.

**Supplementary Figure 3.**
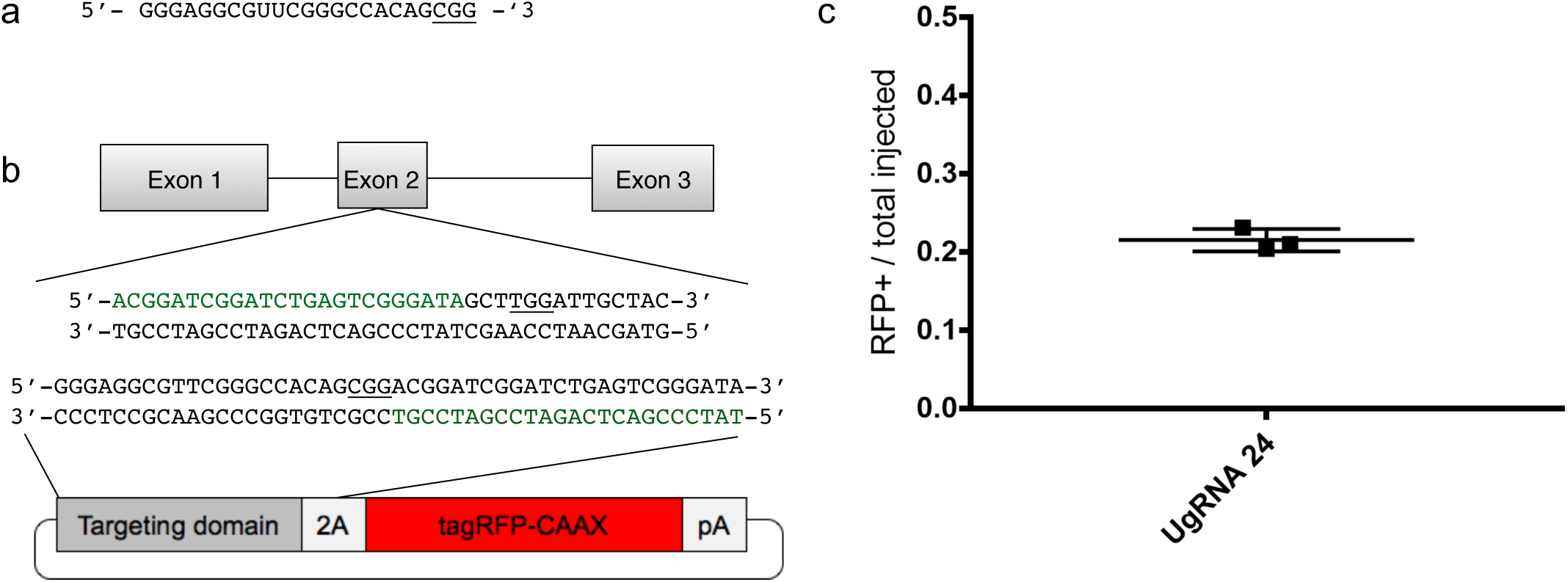
The Universal gRNA (UgRNA) promotes high efficiency targeted integration. (a) Universal gRNA (UgRNA) sequence. Cas9 PAM underlined. (b) Schematic showing the sequence of UgRNA in the targeting domain of the knock-in cassette. Sequence in green from the *noto* gene is the engineered homology in the donor vector for HMEJ. The Cas9 Wierson, Welker, Almeida et al. GeneWeld: a method for efficient targeted integration directed by short homology PAM is underlined. (b) Frequency of injected embryos displaying RFP expression in the notochord compared to total injected embryos following *noto* targeting using UgRNA to liberate the homology in the donor.

**Supplementary Figure 4.**
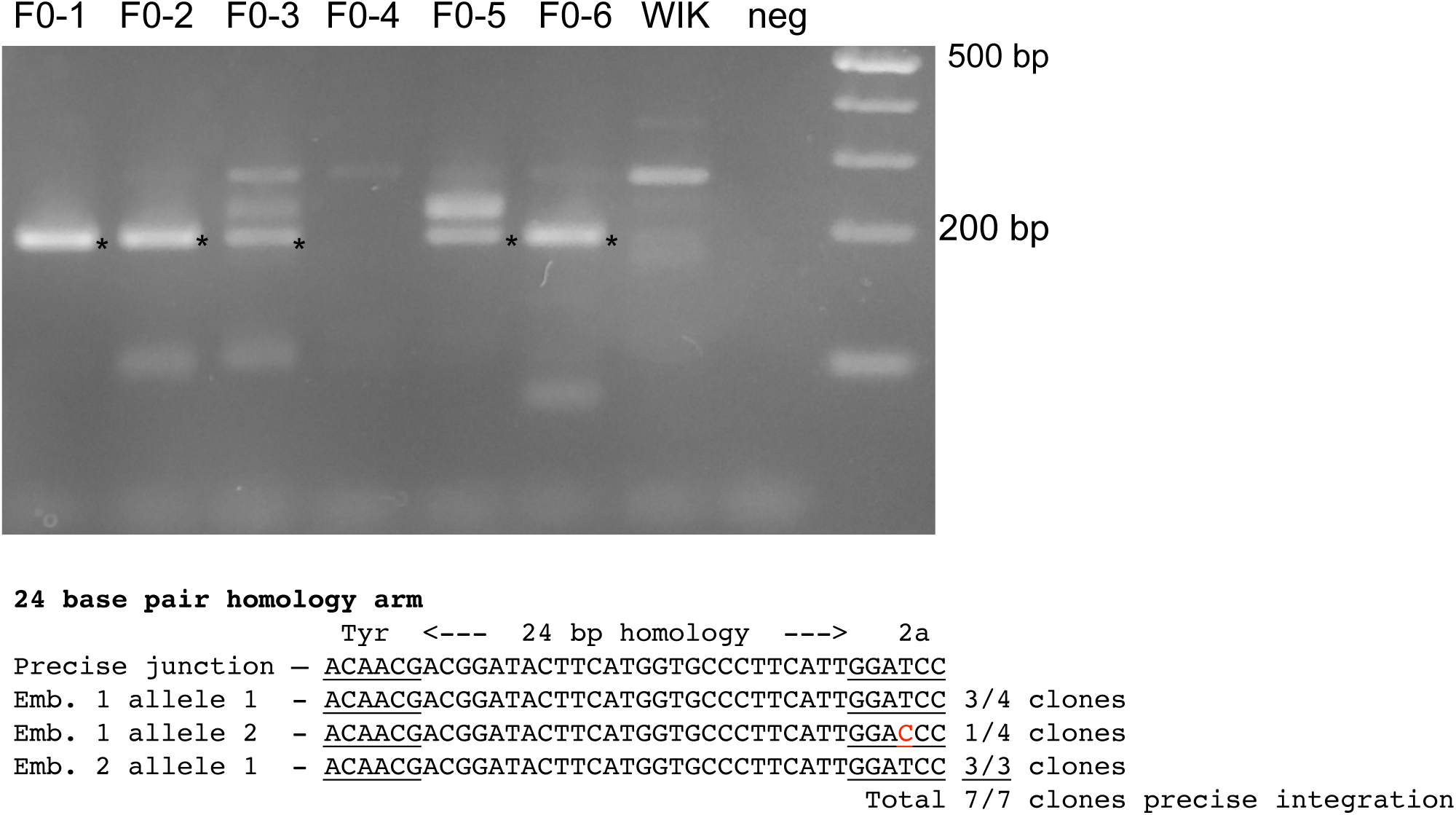
Integration of Gal4/VP16 amplifies signal of targeted *tyr*. (a) PCR of 5’ junction fragments and sequencing results from junction fragments between the pGTag vector and the *tyr* locus amplified from randomly selected RFP negative embryos after injection with GeneWeld reagents for targeting *tyr* with 2A-tagRFP-CAAX-SV40pA. Most F0 injected zebrafish contain the expected 5’ junction fragment (marked with an ‘*’). The junction fragments from F0-1 and −2 were isolated for sequencing, and precise integrations were observed (b) Efficiency of 5’ homology integration to target RFP or GAL4/VP16 into *tyr* and detect RFP expression. Data represents mean +/− s.e.m. of 3 independent targeting experiments. p values calculated using two-tailed unpaired t-test.

**Supplementary Figure 5.**
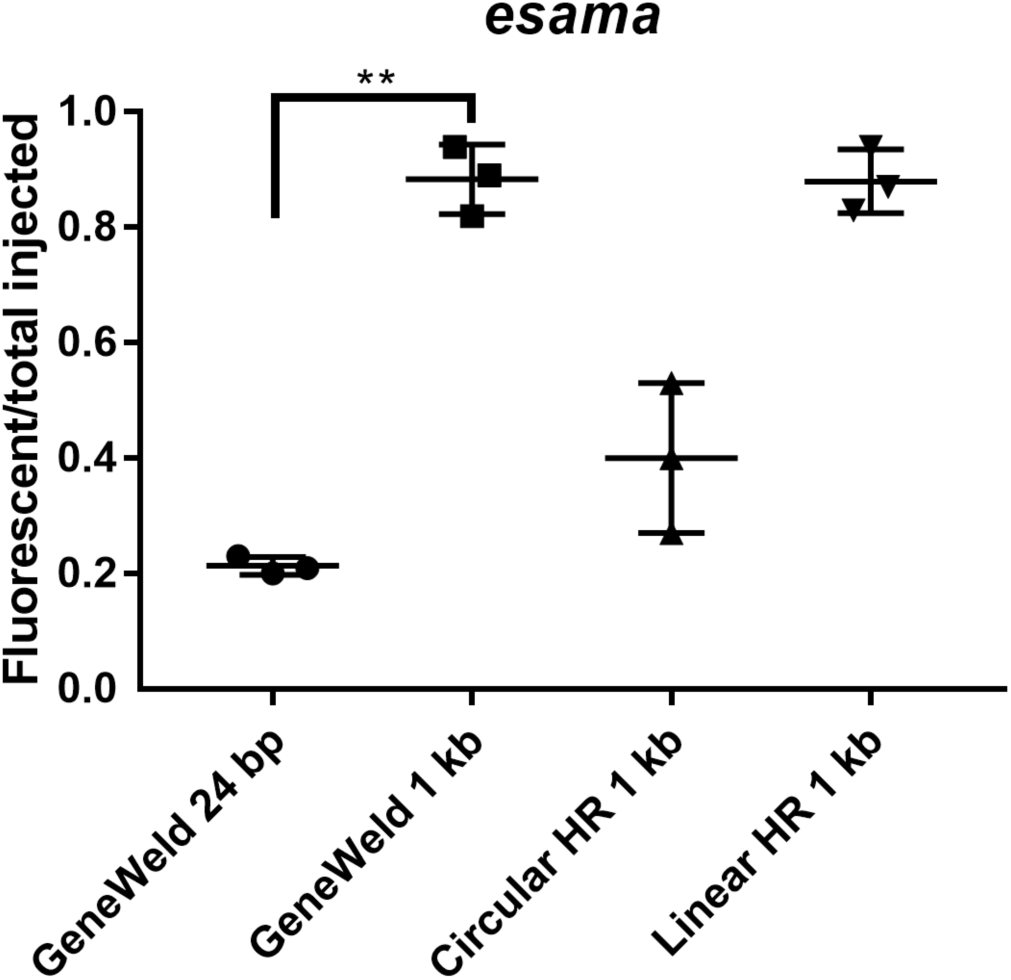
GeneWeld targeting efficiency comparison. Comparison of the frequency of RFP expressing embryos after targeting *esama* exon 2 using GeneWeld 24/24 bp homology, Geneweld 1kb/1kb homology, Circular HR 1kb/1kb (injection did not include UgRNA), Linear HR 1kb/1kb (donor was linearized before injections). Increasing the length of the homology arm to 1kb significantly increased the frequency of RFP expressing embryos using GeneWeld (*p*=0.0001), Circular, or Linear template. Data represents mean +/− s.e.m. of 3 independent targeting experiments. *p* value calculated using Students *t* test.

**Supplementary Figure 6.**
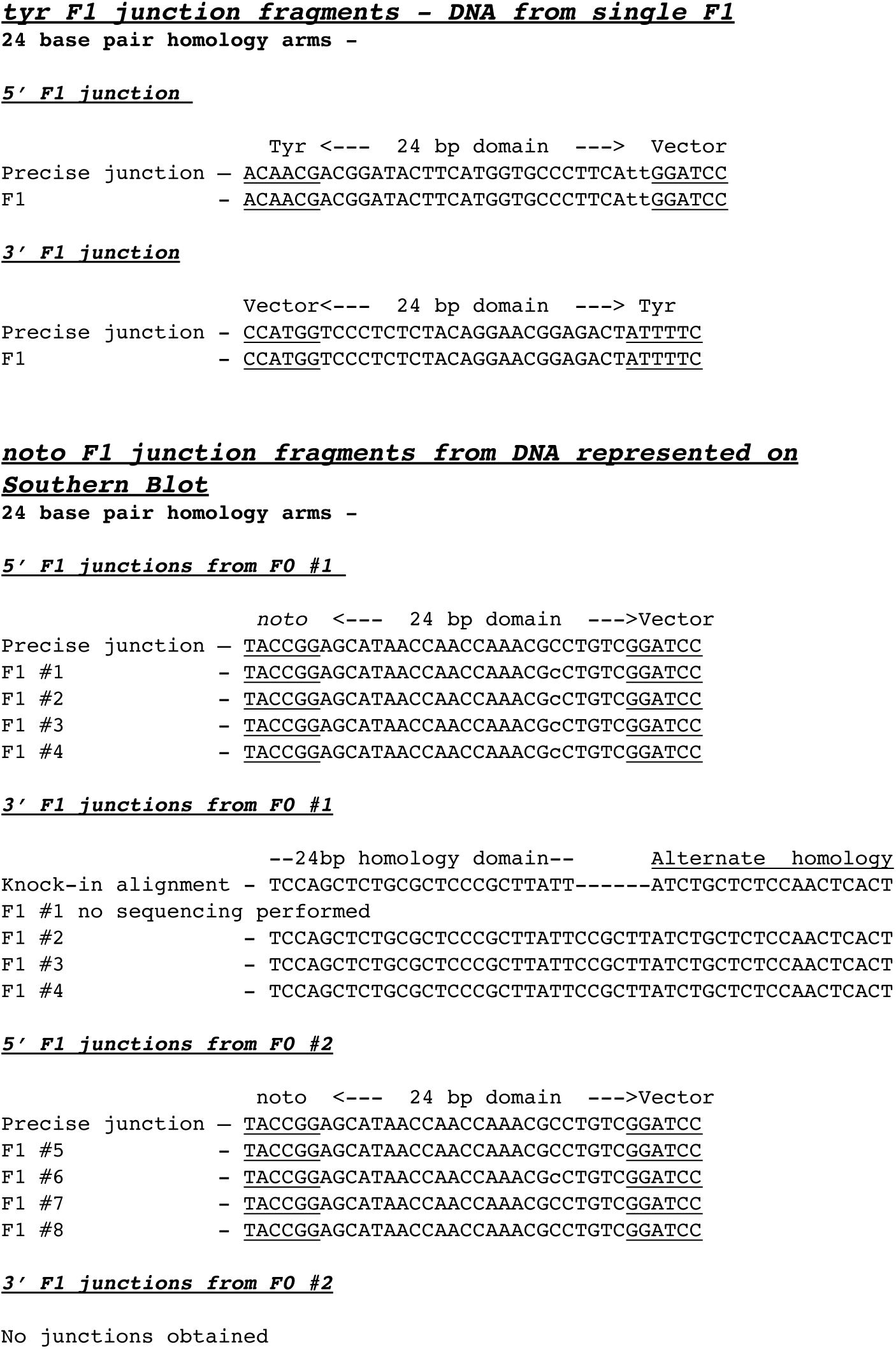
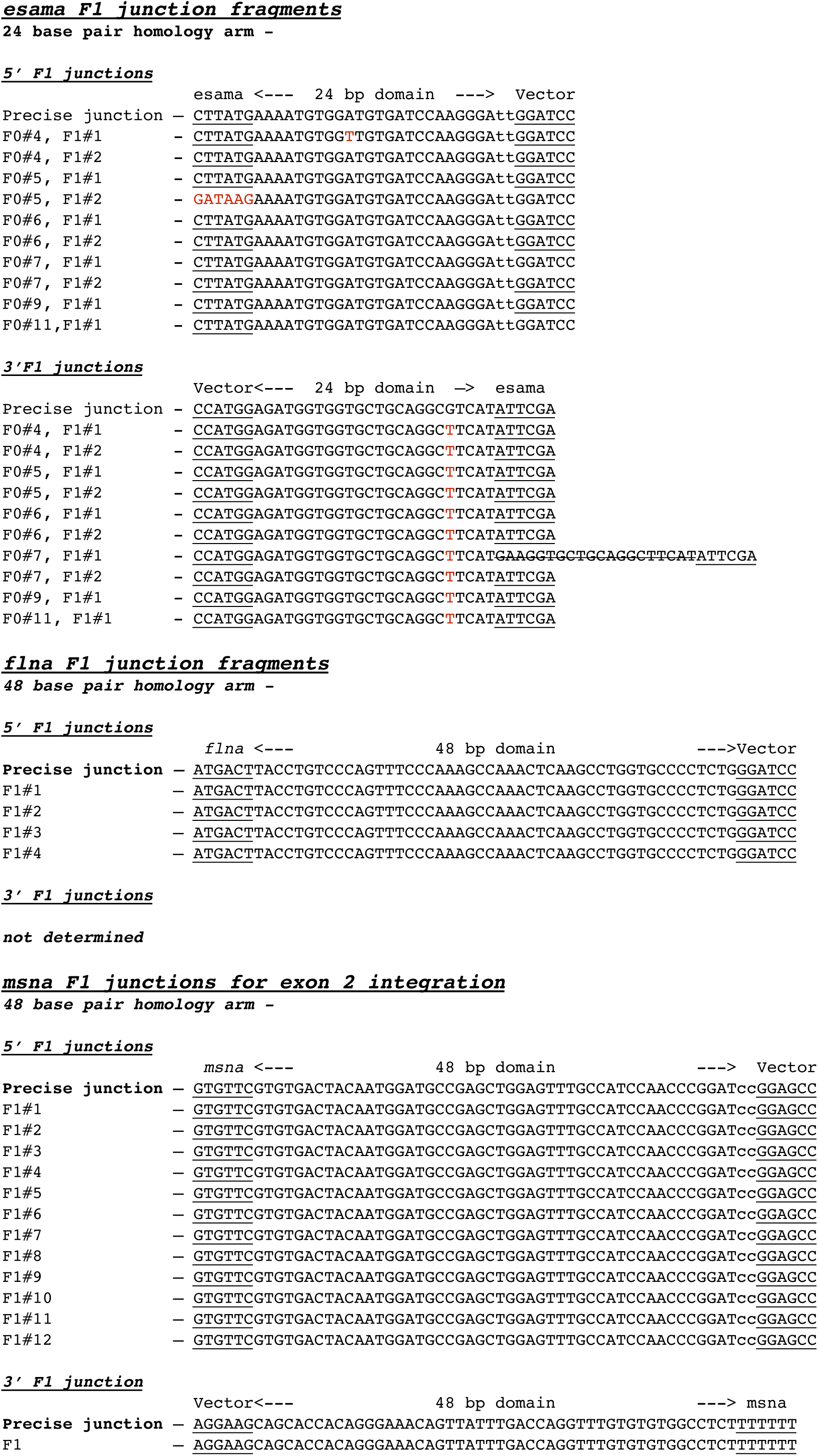
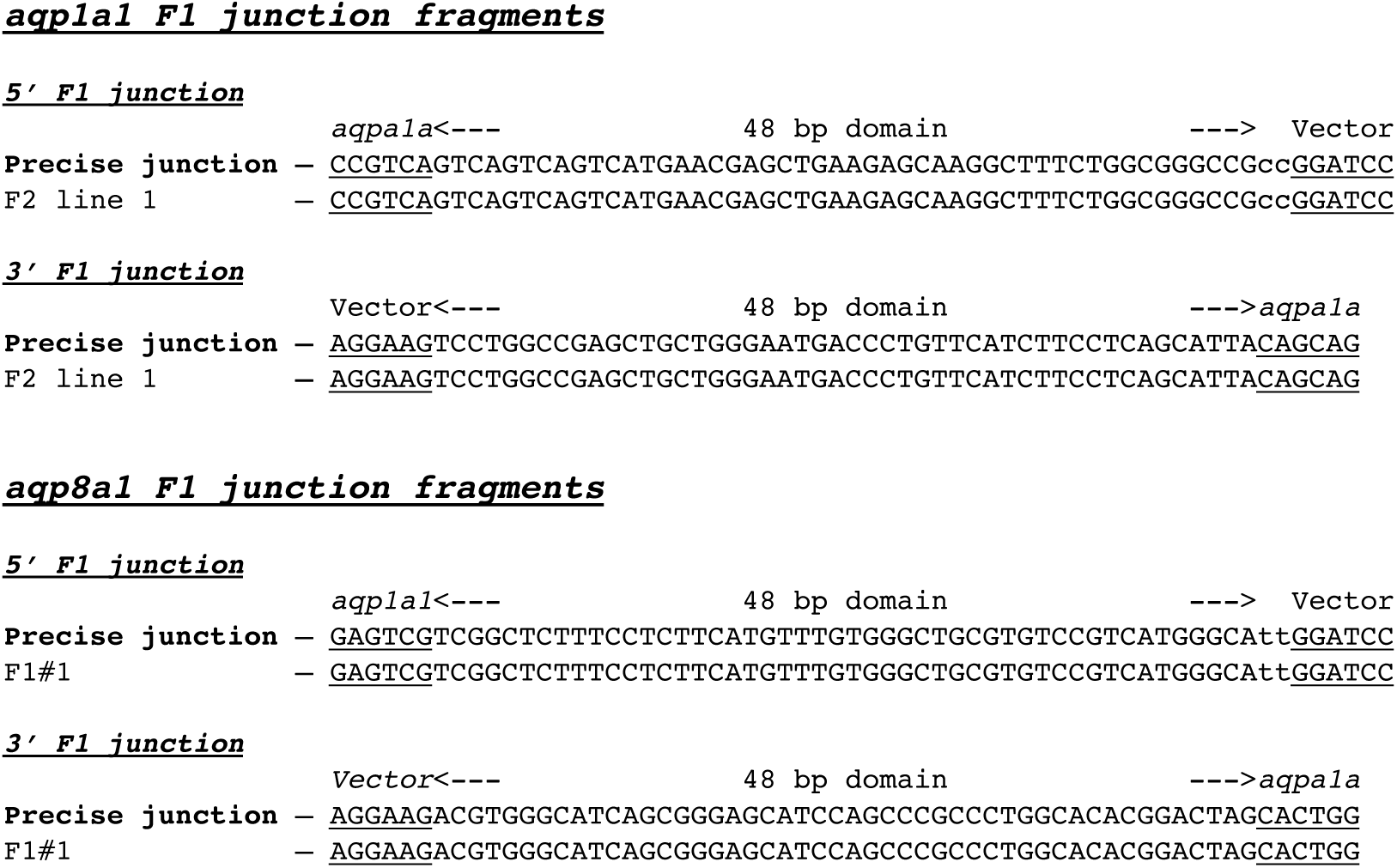
Sequence of PCR junction fragments amplified from genomic DNA from F1 transgenic zebrafish adults generated by GeneWeld targeted integration. Precise integration at the 5’ and 3’ ends in F1 progeny from F0 founder fish targeted at *tyr*, Wierson, Welker, Almeida et al. GeneWeld: a method for efficient targeted integration directed by short homology *esama*, *flna*, *msna*, *aqp1a1*, and *aqp8a1*. *noto* F1 progeny from founder #1 had a precise 5’ junction and imprecise 3’ junction. *noto* F1 progeny from founder #2 had 5’ precise junction; no 3’ junction was amplified by PCR. Lowercase letters represent “padding” nucleotides used to bring homology in frame of the coding region based on Cas9 cut site. Red letters represent mismatches unless otherwise noted below. *esama* F1 3’ junctions contain a single nucleotide variant shown in red. One exama F1 3’ junction included a 20 bp insertion (strike-through).

**Supplementary Figure 7.**
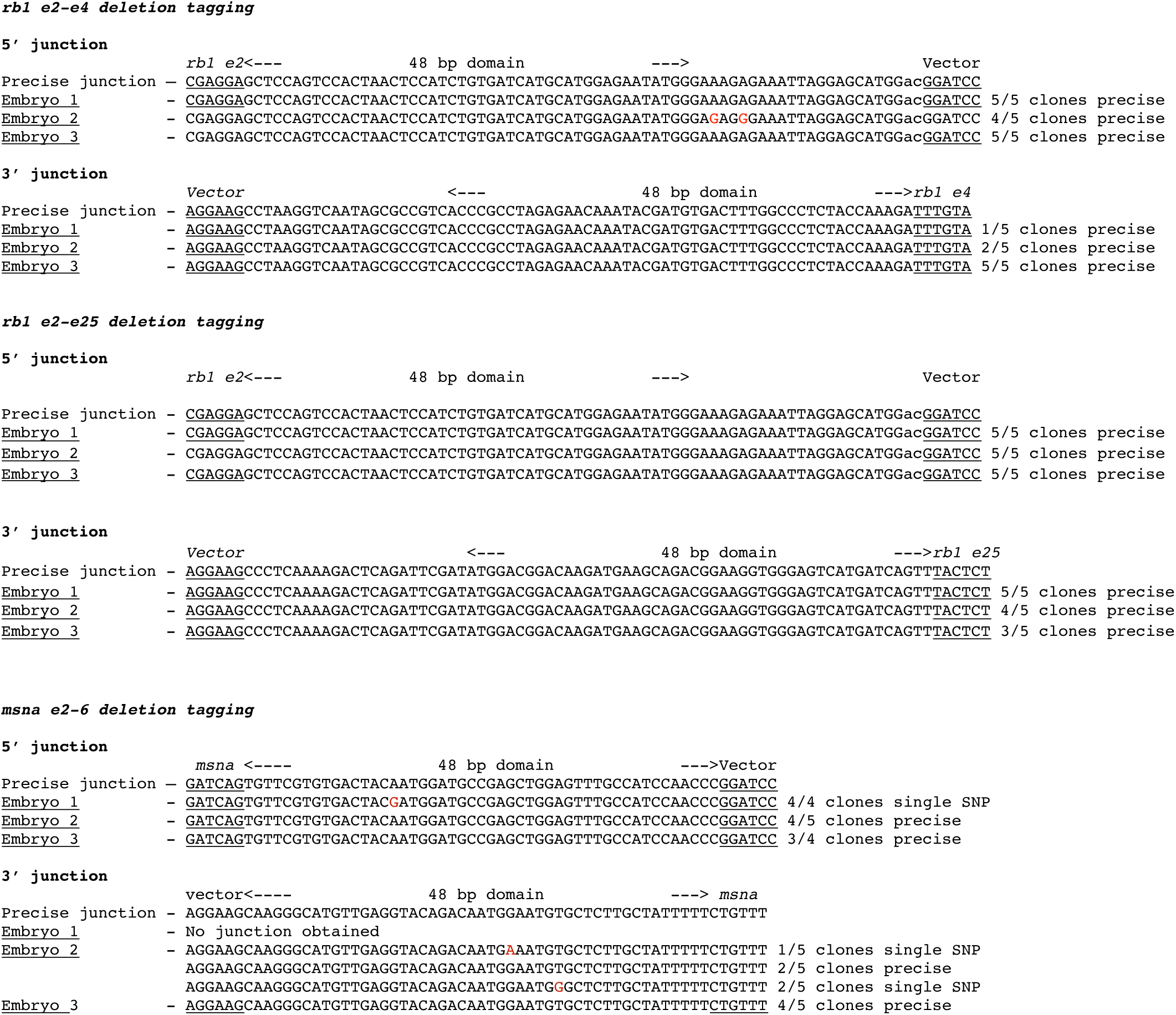
Sequences of 5’ and 3’ junction fragments from *rb1* exon 2-4, *rb1* exon 2-25, and *msna* exon 2-6 deletion tagged alleles in F0 injected embryos. Detection of precise and imprecise 5’ and 3’ junction fragments in somatic tissue of F0 embryos targeted with two guides targeting two exons and a pGTag-Gal4VP16 donor with 5’ and 3’ homology arms corresponding to the 5’ exon and 3’ exon target sites. Cloned PCR amplicons were sequenced from 3 individual embryos for each targeted deletion tagging experiment.

**Supplementary Figure 8.**
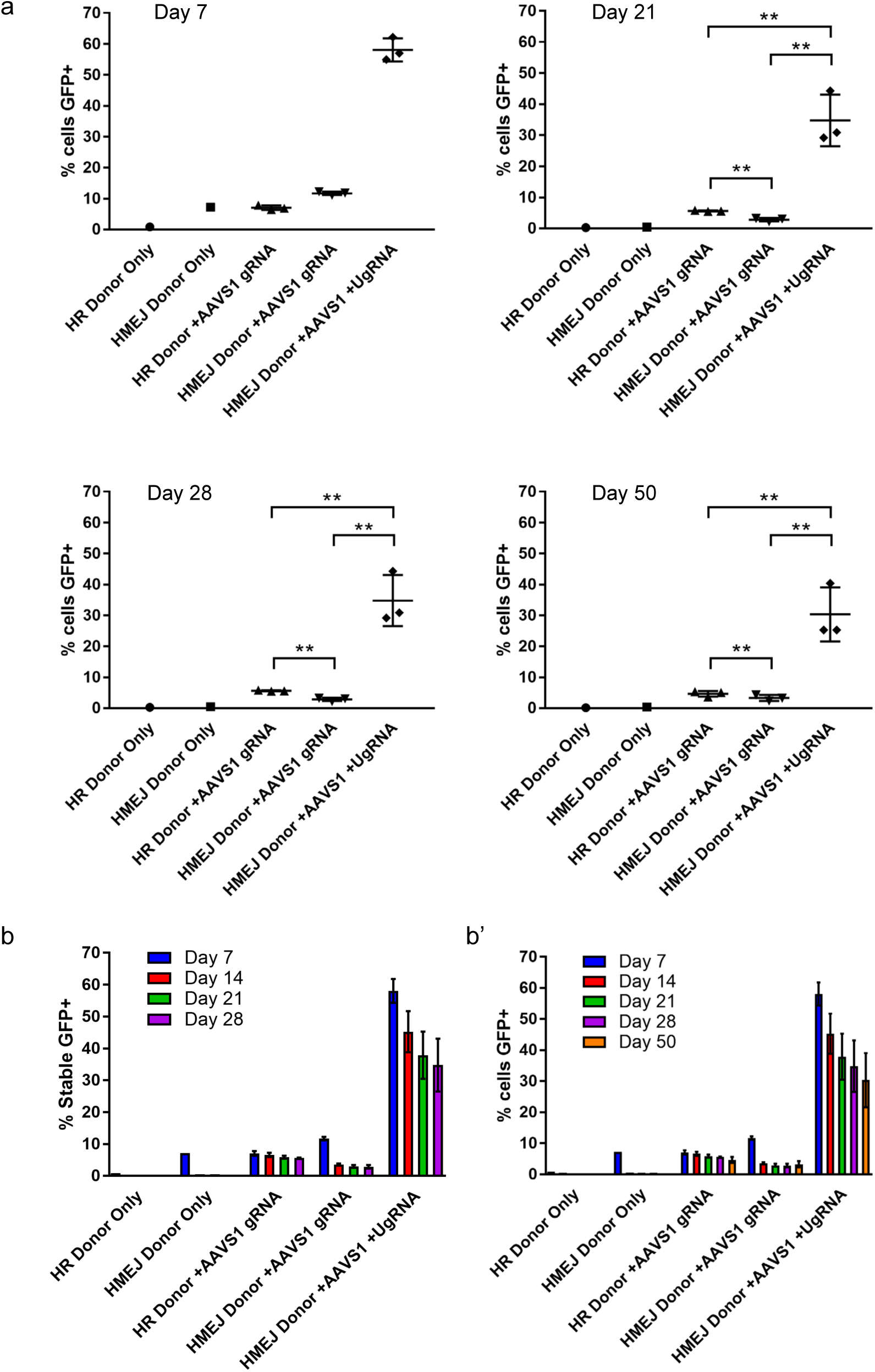
HMEJ-mediated targeted integration of an MND:GFP reporter at the *AAVS1* locus in human K-562 cells. FACs sorted percent of GFP+ cells out of total K-562 cells at day 7, 21, 28, and 50. (b) Summary data for percent of stable GFP+ K-562 cells from day 7, 14, 21, and 28. (b’) Summary data for percent of total cells GFP+ from day 7, 14, 21, 28, and 50. Data represents mean +/− s.e.m. of 3 independent targeting experiments. p values calculated using two-tailed unpaired t-test.

**Supplementary Figure 9.**
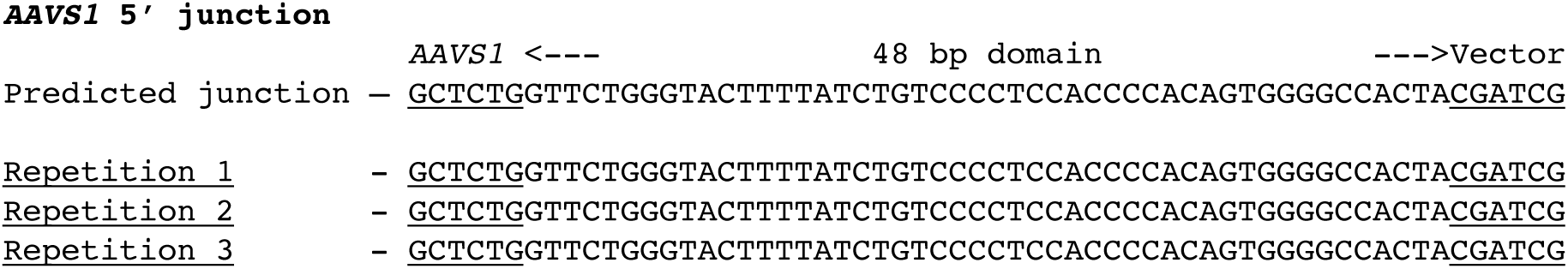
Direct sequencing of 5’ junction PCR products derived from three independently targeted bulk cell populations. (a) Direct sequencing of 5’ junction PCR products derived from three independently targeted bulk cell populations. 48 bp HMEJ homology region and remainder of genomic *AAVS1* gRNA are indicated. Genomic sequence is Wierson, Welker, Almeida et al. GeneWeld: a method for efficient targeted integration directed by short homology directly left of the 48 bp HMEJ region and vector sequence is directly to the right of the *AAVS1* gRNA cut site.

**Supplementary Table 1.**
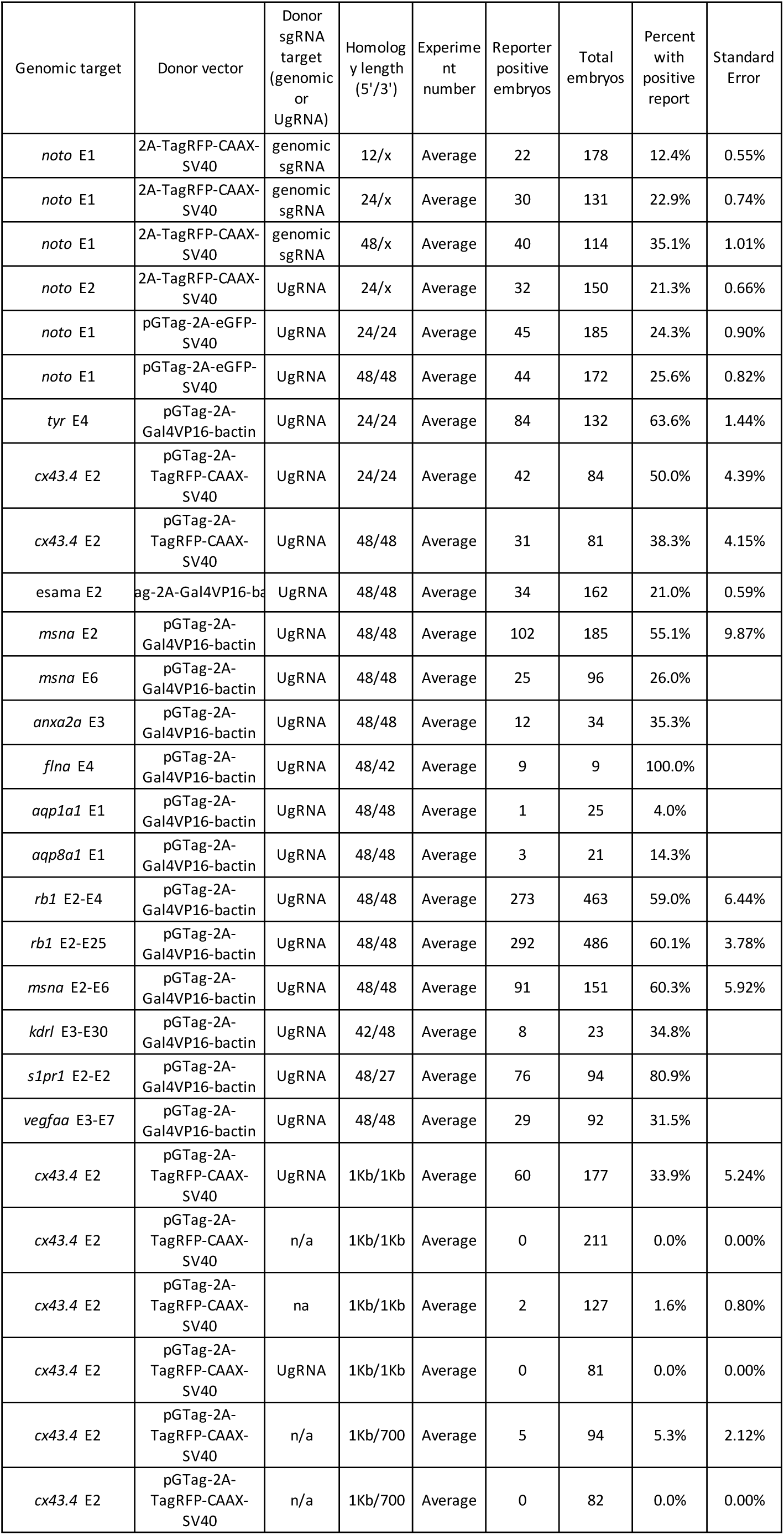
Gene targeting experiments and knock-in averages.

**Supplementary Table 2.**
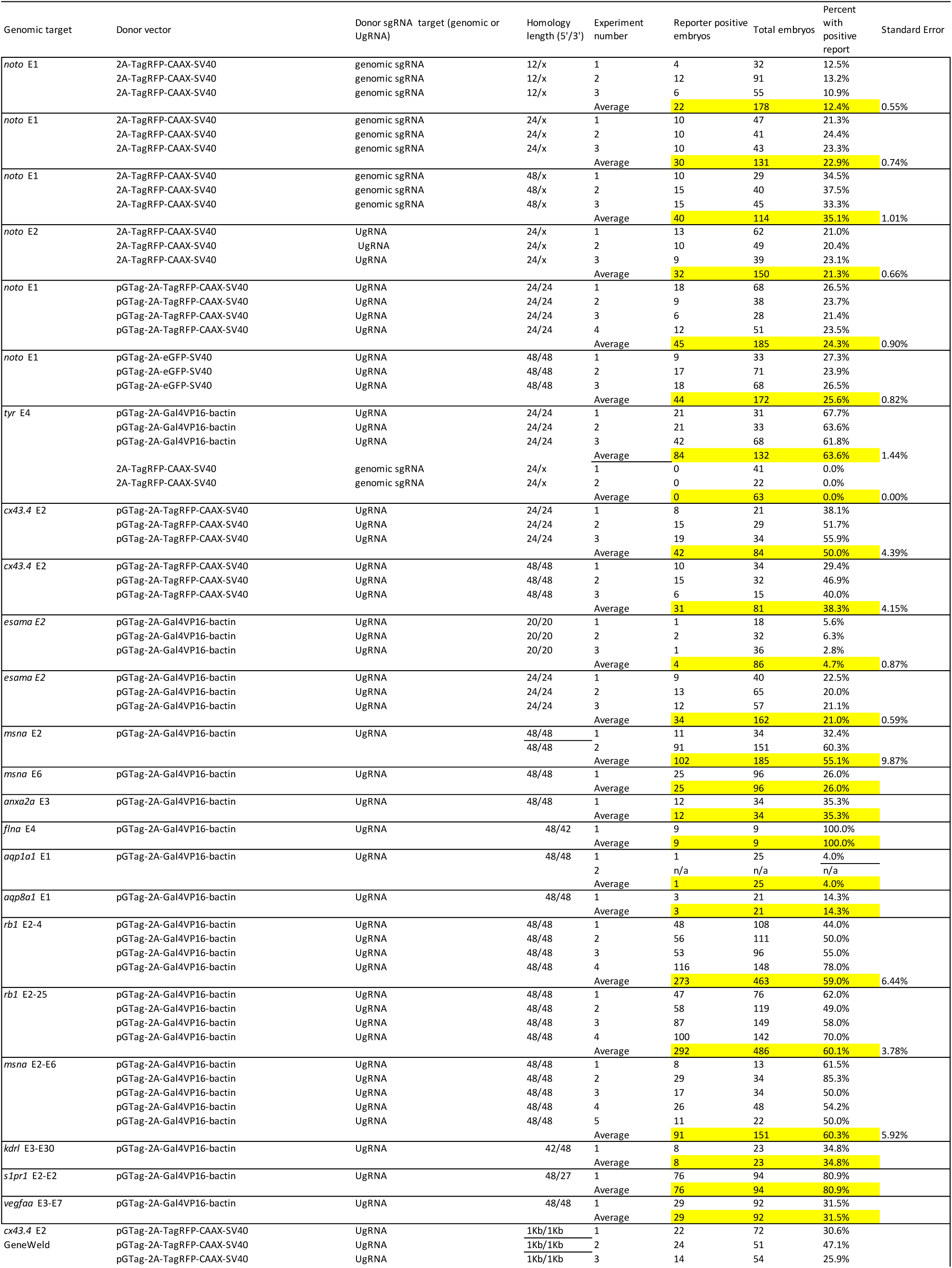
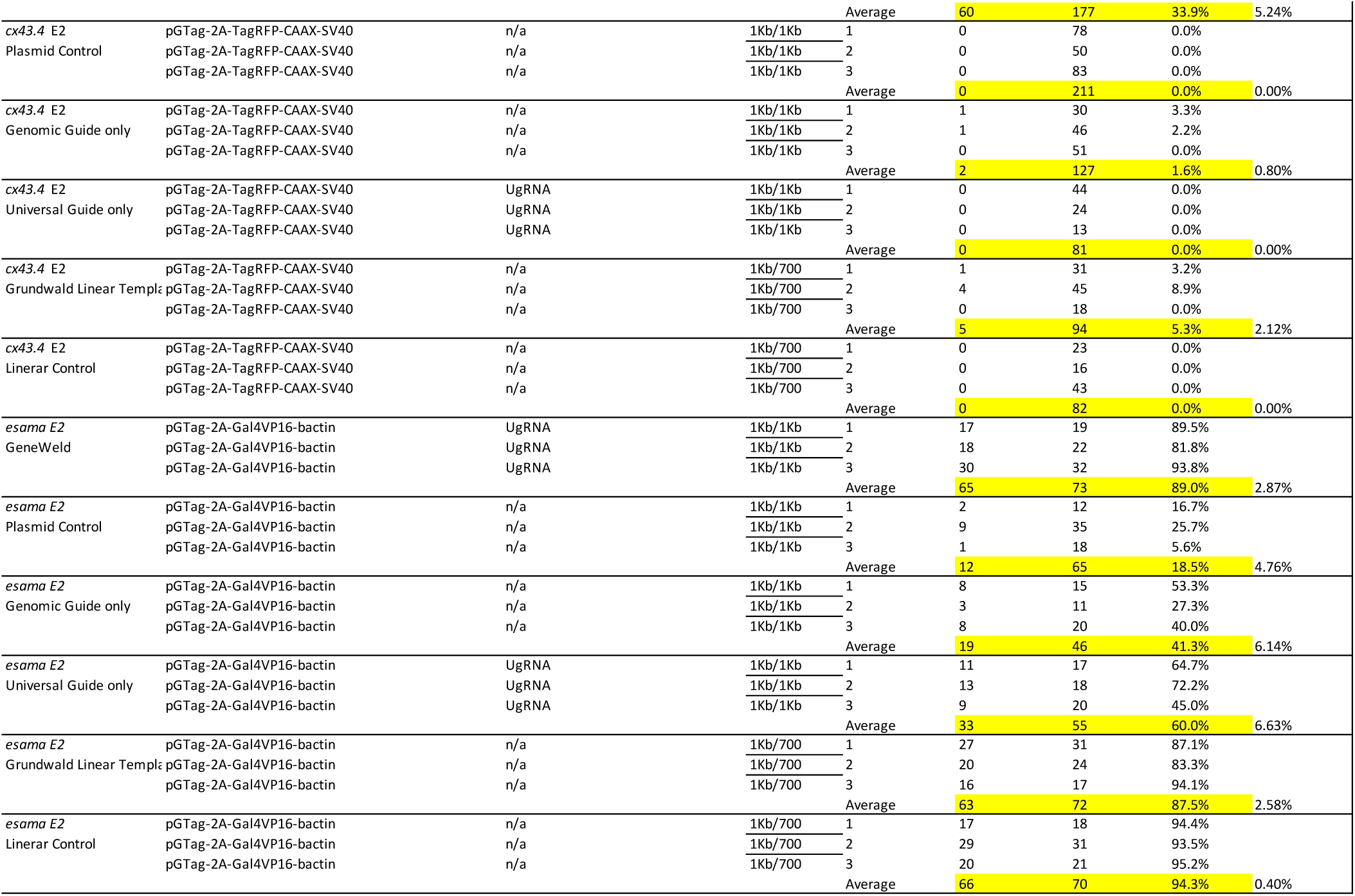
Gene targeting experiments and knock-in percentages.

**Supplementary Table 3.**
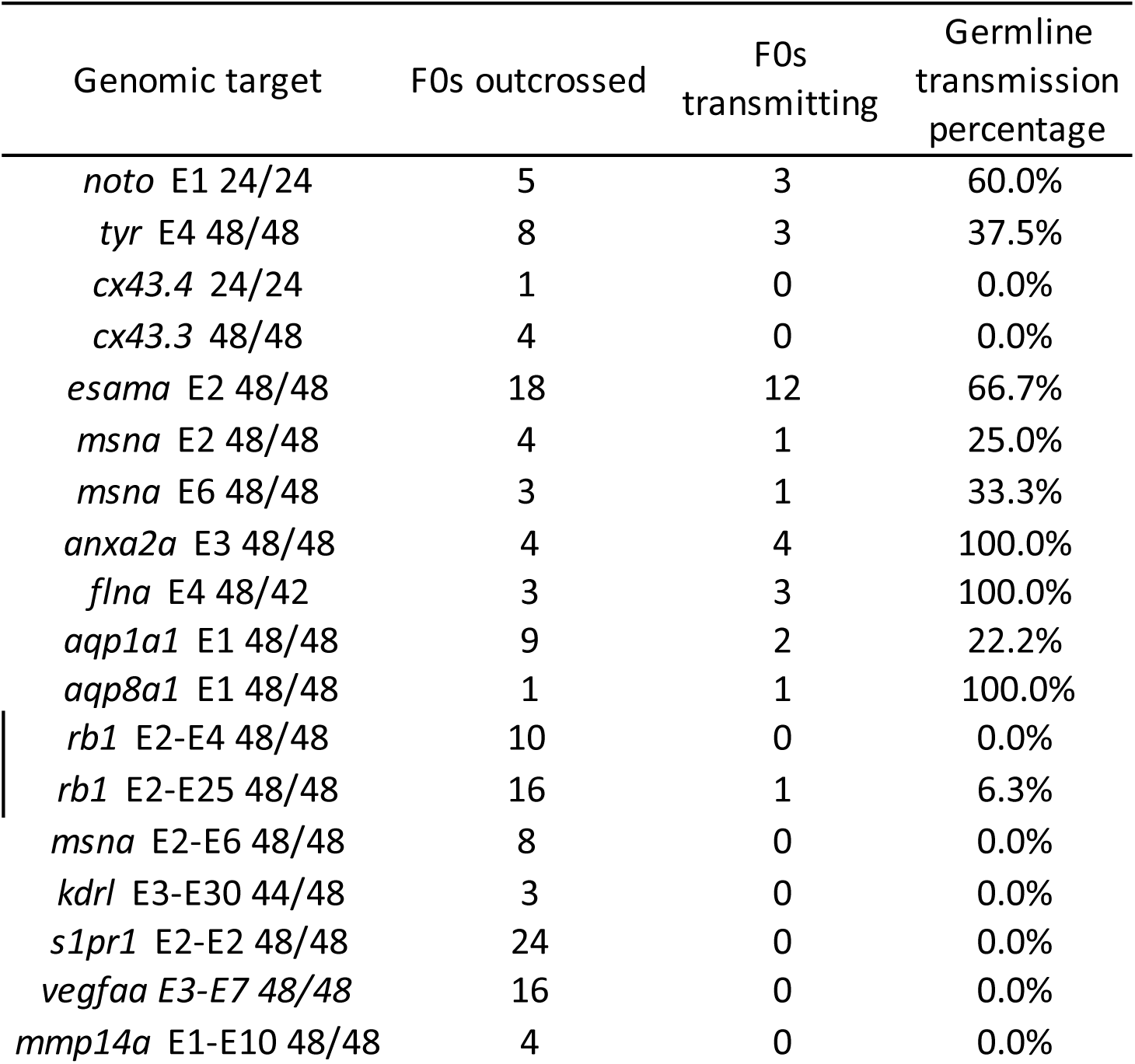
Germline Transmission

**Supplementary Table 4.**
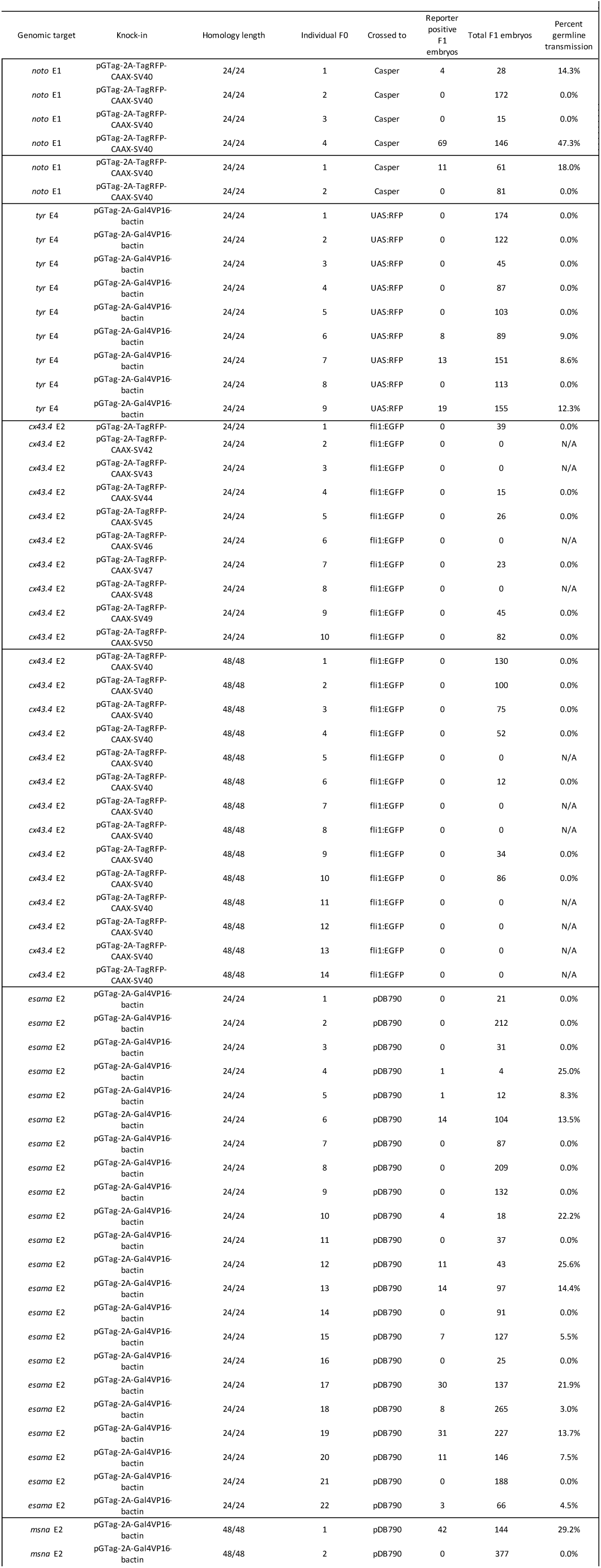
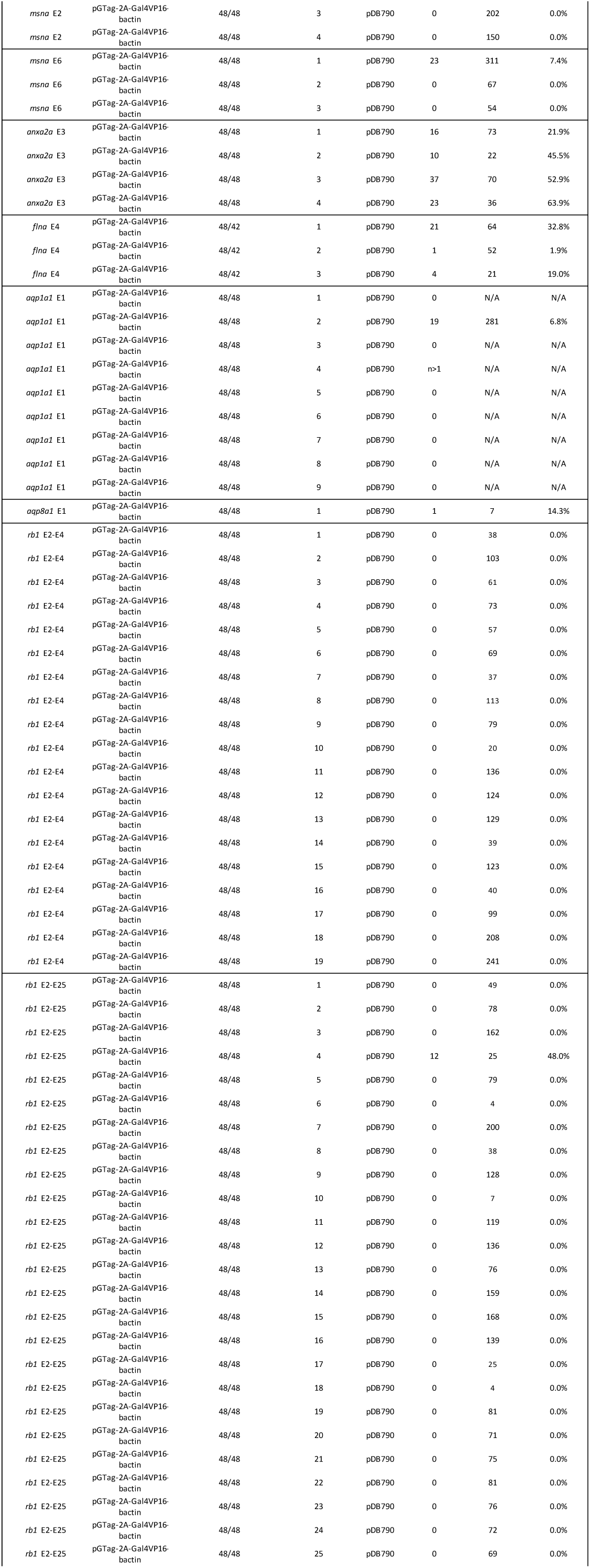
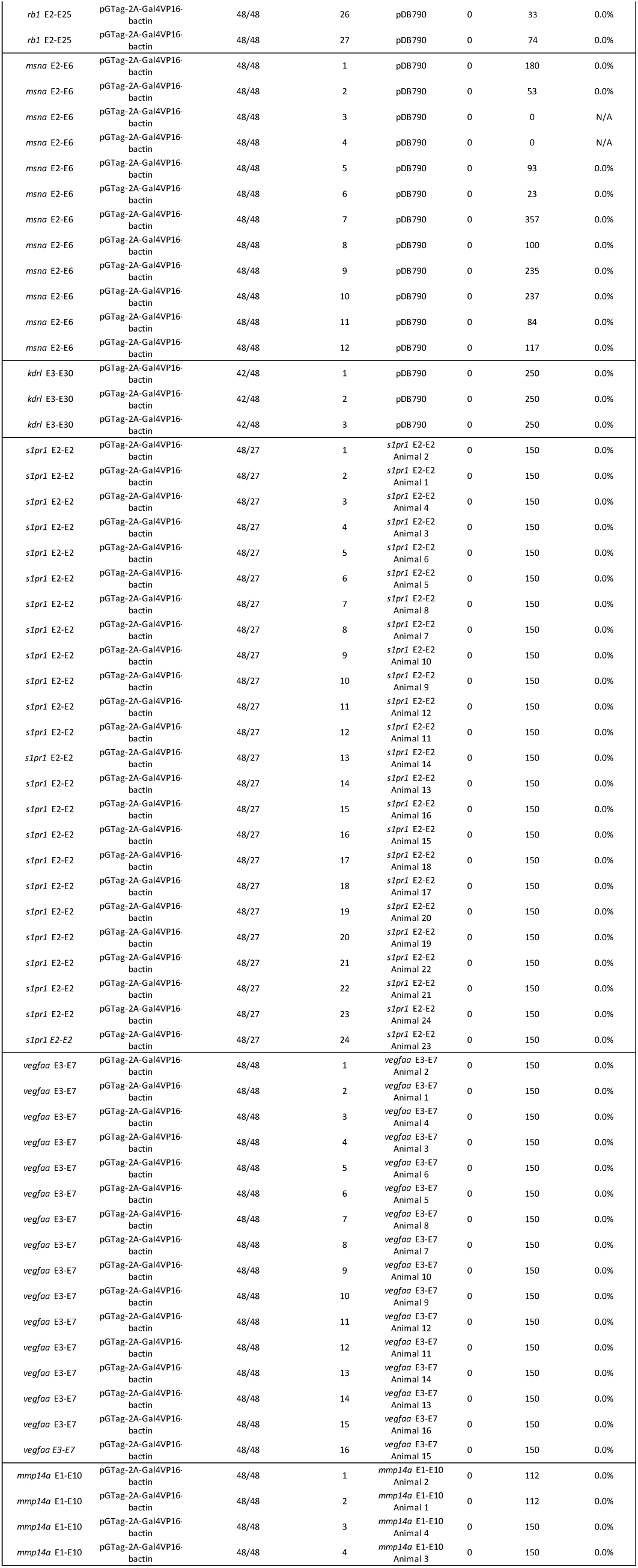
F0 outcrosses for transmission; asterisk denotes in-cross

**Supplementary Table 5.**
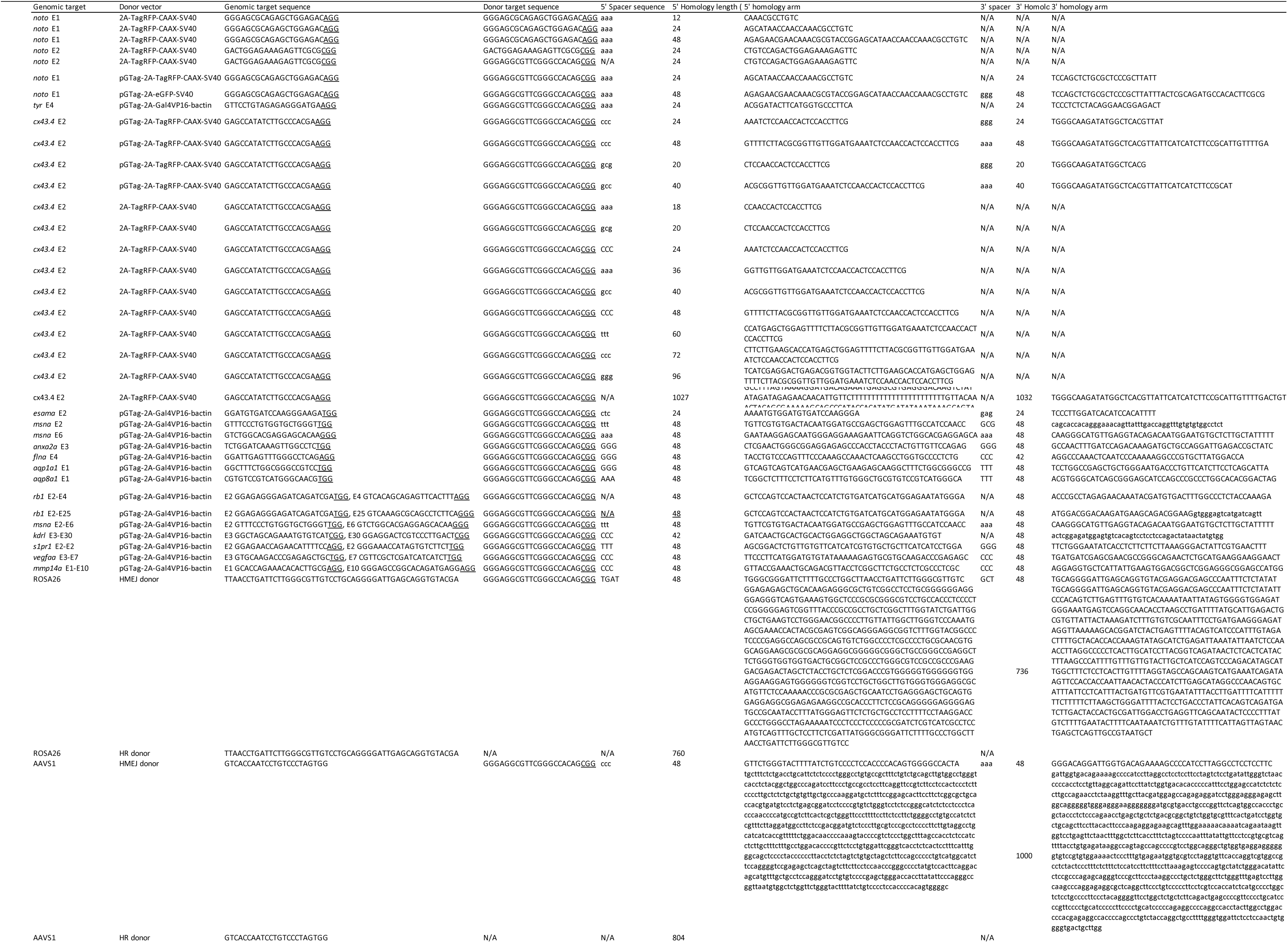
Targeting domain information for all gene targeting experiments. N/A means not applicable

**Supplementary Table 6.**
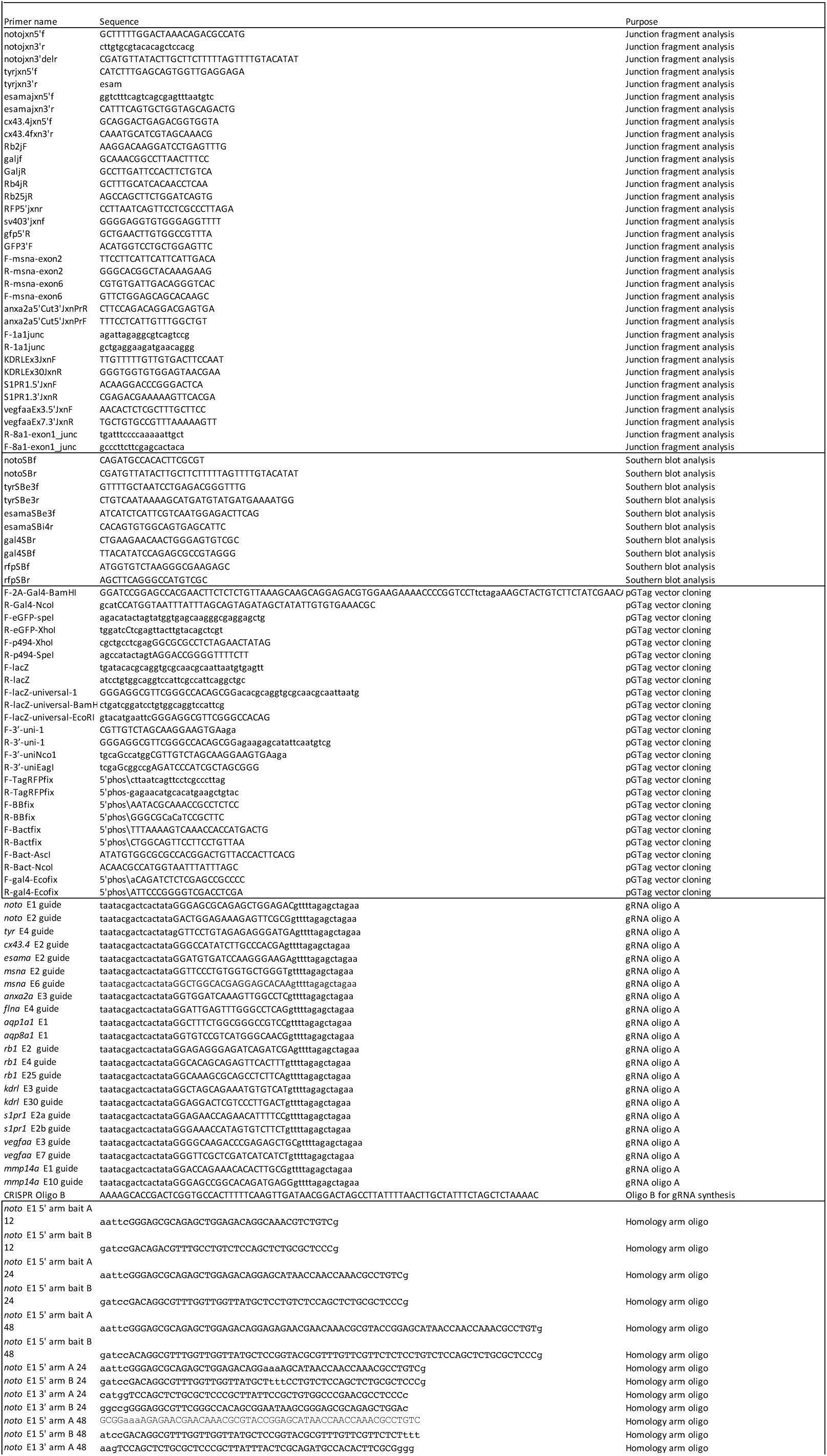
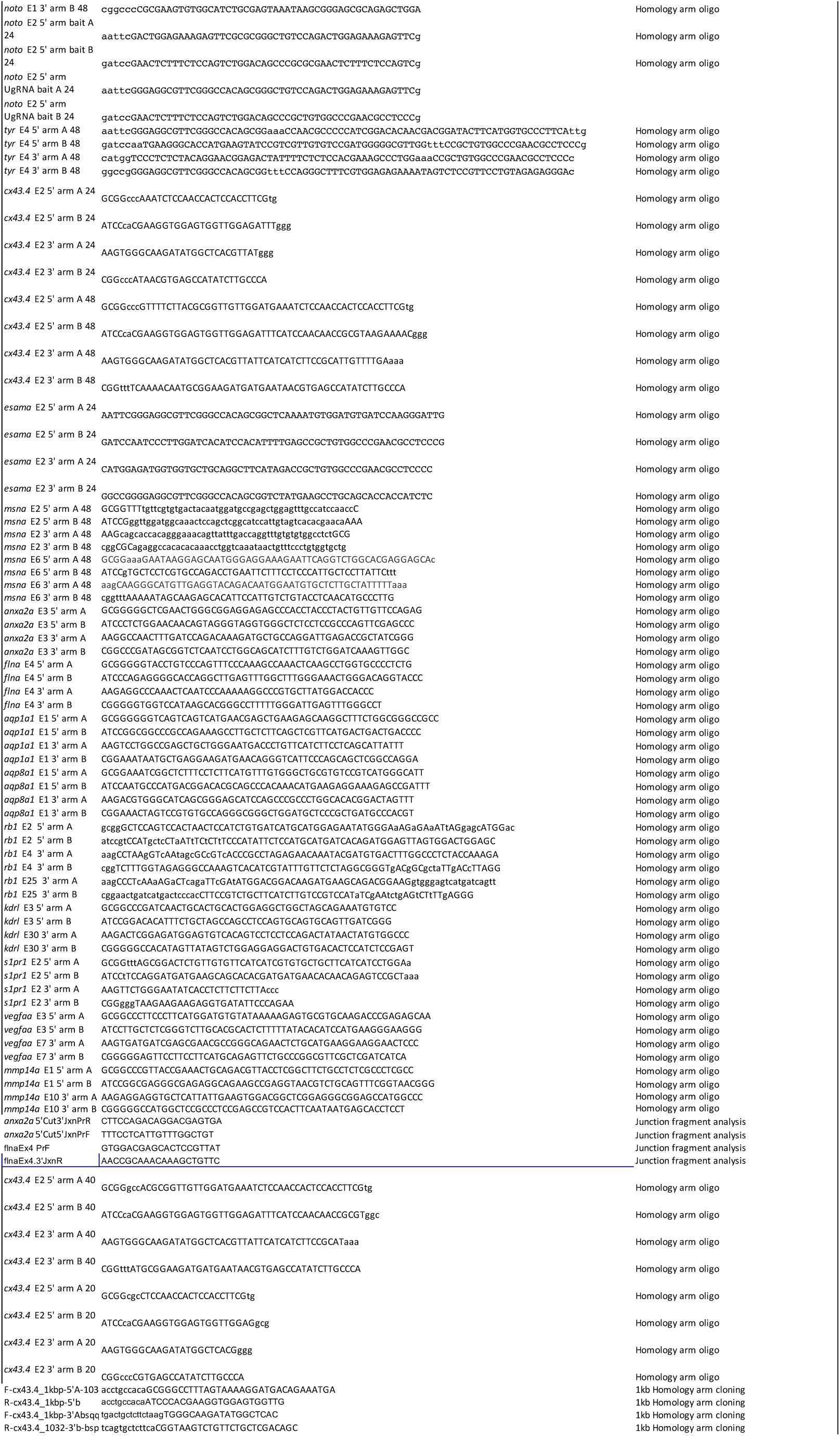
Primes and oligos

## Gene Targeting Protocol for Integrations with pGTag Vectors using CRISPR/Cas9

Targeting strategy (Figure 1):

A. Selection of a CRISPR/spCas9 target site downstream of the first AUG in the gene of interest
B. Synthesize sgRNA and spCas9 mRNA
C. Injection of sgRNA and spCas9 mRNA
D. Testing for indel production/mutagenesis
E. Design short homology arms
F. One Pot Cloning of Homology Arms into pGTag Vectors
G. Injection of GeneWeld reagents (spCas9 mRNA, Universal sgRNA (UgRNA), genomic sgRNA and pGTag homology vector) into 1-cell zebrafish embryos
H. Examine embryos for fluorescence and junction fragments

**Figure 1.**
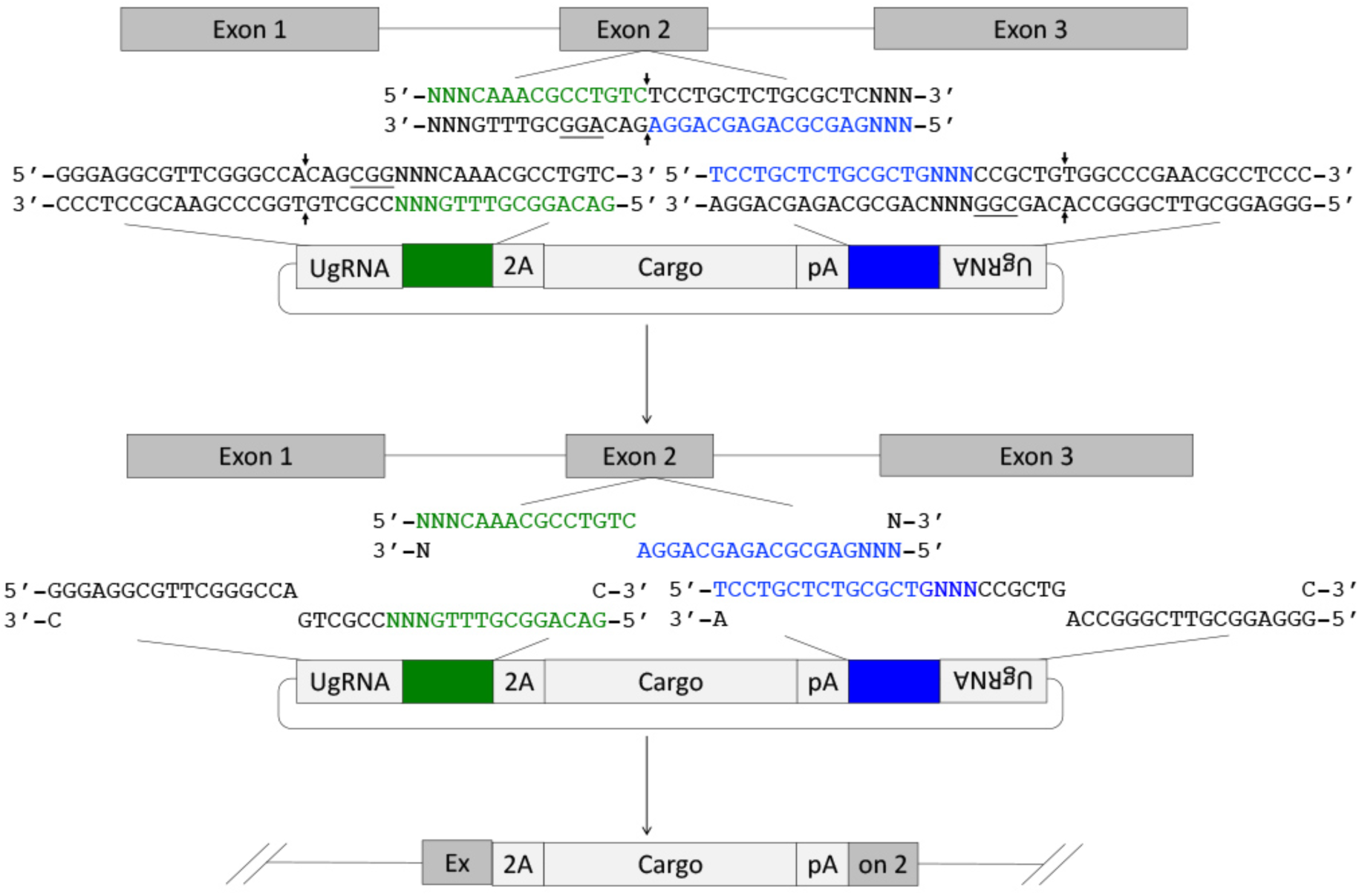
Targeting integration of the pGTag vectors into the 5’ region of a gene. Upon CRISPR/Cas9 targeting and cutting of both the genome (with a sgRNA) and plasmid donor (with UgRNA), the genomic and plasmid DNA likely undergo end resection mediated by the MRN complex and *ExoI*, resulting in annealing of complementary homology arms. This promotes precise homology-directed integration of cargo DNA at the CRISPR/Cas9 double-strand break.

### A. Selection of a CRISPR/spCas9 target site downstream of the first AUG in the gene of interest

1. To select a CRISPR/Cas9 target site in a 5’ exon, find and download the targeted gene’s genomic and coding sequences.

a. At <ensemble.org> Search for the gene name of interest for the species of interest and open the Transcript page.
b. In the left-hand side bar click on “Exons” to find the first coding exon and initiation ATG. If there are alternative transcripts for the gene, make sure there are not alternative initiation ATGs. If there are alternative start codons, target the first 5’ exon that is conserved in all transcripts to generate a strong allele.
c. Download the transcript and 5’ exon to be targeted as separate sequence files.
d. Using ApE: <http://biologylabs.utah.edu/jorgensen/wayned/ape/> annotate the coding sequence with the exons.

2. Use CRISPRScan (http://www.crisprscan.org/) (Moreno-Mateos et al., 2015) to efficiently identify target sites and generate oligos for sgRNA synthesis for the target gene.

a. Select the “Predict gRNAs” on the lower right-hand side of the home page of the CRISPRScan website.
b. Paste the 5’ exon sequence into the indicated box. If the exon is very large, start with a small amount of sequence. Ideally exon sequence of ~200 bp near the desired target site. Do not design CRISPRs to intron/exon borders. If there are problems with the copy and pasting of exon sequence, first paste the sequence into a new ape file, save, then copy and paste from the new file.
c. Select “Zebrafish (Danio rerio)” as the species
d. Select “Cas9 – nGG” as the enzyme.
e. Select “In vitro T7 promoter”.
f. Click on “Get sgRNAs.” Examine the output. The generated targets are ranked by CRISPRScan from high to low. Select a target site (the 20 bp that are capitalized in the oligo column) from those given by CRISPRScan using the following criteria (The best gRNAs will have all of these):

i. An exact match to the genomic locus., When an oligo is clicked on the page will display additional information to the right. In the section called “Site Type” any mismatches in the oligo are displayed. Exact matches including 5’GG- are ideal for in vitro transcription and 100% genomic target match.
ii. The target is in the desired location of the gene.
iii. The Target is on the reverse (template) strand. Reverse strand guides are more favorable, but either will work
iv. A high CRISPRScan score, and a lower CFD score. However, lower score sgRNA targets may work fine.
g. Annotate the selected target sequence in the transcript sequence files.
h. For sgRNA synthesis the entire oligo sequence from CRISPRscan containing the selected target will need to be synthesized. This oligo is represented as “Oligo A” in Figure 2.

**Figure 2.**
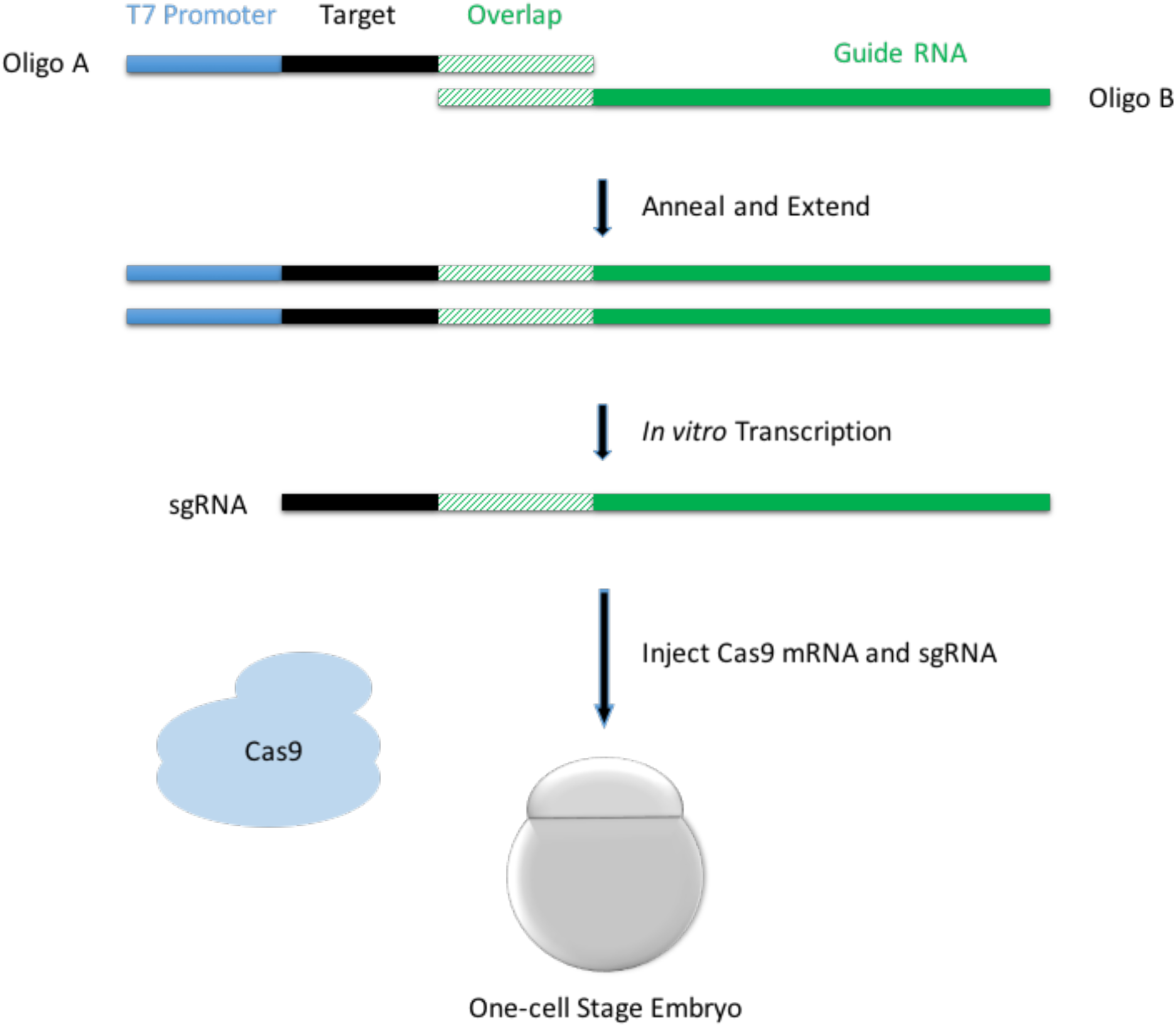
Cloning-free gRNA synthesis. Oligo A is composed of the T7 promoter at the 5’ end, target sequence for gRNA, and gRNA overlap sequence for gRNA synthesis. CRISPRScan provides direct output for Oligo A. The strategy for gRNA production using Oligo A is based on (Varshney et al., 2015).

3. **Alternative to CRISPRScan:** Designing “CRISPR Oligo A” from a genomic target sequence. Skip this section if Oligo A was designed with CRISPRScan.

If the target sequence was identified using tools other than CRISPRScan, Oligo A can be designed manually. (Note: CRISPRScan will use a shorter overlap region but this does not affect template production). Add T7 and Overlap sequences (see Figure 2) to the 20 bp of target sequence without the PAM. Oligo A for the targeted gene will look like the example below:

5’-TAATACGACTCACTATAGGNNNNNNNNNNNNNNNNNNGTTTTAGAGCTAGAAATAGC-3’

The sequences in blue (first 17 characters) are the T7 promoter, the grey GG are part of the T7 promoter and ideally are part of the target sequence (see below), the Ns are the target sequence, and the sequence in green (last 20 characters) are the overlap region to synthesize the non-variable part of the sgRNA. The T7 promoter works optimally with the two grey GGs, however, these GGs will be transcribed by T7 and thus become a part of the sgRNA. Target sequences that contain the GGs may work better, but there are differing reports in the literature on the importance of this (Moreno-Mateos et al., 2015). If possible, select a target that starts with GG. Refer to Moreno-Mateos et al., 2015 for other gRNA architectures with variations on the 5’GG motif.

a. If the target sequence did not have two Gs at the beginning, additional G’s will need to be added to the start of the target sequence for efficient transcription as outlined below: *The lower case ‘g’ is an extra ‘G’ not in the genomic sequence; the upper-case G *is* in the genomic. Lower case gs will not base pair with the genomic target.

i. without GG: ggN NNN NNN NNN NNN NNN NNN N (22 bp) – 2 bases are added,
ii. with one G: gGN NNN NNN NNN NNN NNN NNN (21 bp) – one base is added, G is part of the target sequence.
iii. with two G: GGN NNN NNN NNN NNN NNN NN (20 bp) – no bases are added, GG is part of the target sequence. Oligo A is made by taking this target sequence with 5’GG and pasting it into a clean file.
b. Copy and paste the T7 promoter sequence to the 5’ end of the target sequence: TAATACGACTCACTATA
c. Copy and paste the Overlap sequence to the 3’ end of the target sequence: GTTTTAGAGCTAGAAATAGC
d. Check the sequences to ensure they are correct and that the PAM is <u>NOT</u> present in this oligo.

4. Oligo B design (Figure 2) contains the conserved guide RNA sequence: All Oligo Bs will be the same and can be ordered in large quantities.

5’-GATCCGCACCGACTCGGTGCCACTTTTTCAAGTTGATAACGGACTAGCCTTATTTTAACTTGCTATTTCTAGCTCTAAAAC-3’

5. To increase yield of the sgRNA synthesis the primers “T7 primer” (5’-TAATACGACTCACTATA-3’) and “3’gRNA primer” (5’-GATCCGCACCGACTCGGTG-3’) are also required.

6. For checking for mutagenesis at the target site, design ~20 bp DNA primers for PCR amplification to amplify at least 130 bp of DNA surrounding the target site. Mutagenesis is estimated through comparison of PCR products from injected and uninjected embryos, by visualizing small insertions and/or deletions (Indels) using electrophoresis, or by sequencing.

a. Primer 3 is used for primer design: http://biotools.umassmed.edu/bioapps/primer3_www.cgi
b. Paste DNA sequence surrounding the target site into the web interface. It is recommended to use 160 – 300 bp of exon sequence centered on the cut site for primer design. Intron sequence can be used, but this often contains polymorphisms that can lead to amplification failure.
c. Locate the target sequence, including the PAM sequence (italicied below), and predict the cut site (3 bp into the target sequence from the PAM represented here by the ‘x’). Mark the targeted exon sequence approximately 65-150 bp on both sides of the cut site by putting [square brackets] around it. Primer3 will design primers outside this sequence. This design allows the primers to be used for both checking of mutagenesis and for junction fragment analysis when checking for integration. Example: CGGCCTCGGGATCCACCGGCC[AGAATCGATATACTACGATGAACAGAGCAAATTTGTGTGTAATAC<u>GGTCGCCACCATGGCCTxCCT*CGG*</u>TTTGCTACGATGCATTTGCACCACTCTCTCATGTCCGGTTCTGGG]AGGACGTCATCAAGGAGTTCATGCGCTTCAAGGTGCGCATGGAGGGCTCCGTGAAC
d. Set the “Primer Size” variables to Min = 130, Opt = 170, and Max = 300. Everything else can be left at the defaults.
e. Click on “Pick Primers”
f. Select primers from the output. Note the “product size” expected and the “tm” or melting temperature of each primer/pair. Smaller product sizes are easier to visualize mutagenesis.

### B. Synthesize the sgRNA

General guidelines and good laboratory practices for working with DNA and RNA. DNA, RNA and the enzymes are sensitive to contamination from dust and skin. Following these guidelines will prevent the degradation of the DNA and RNA you are trying to make:

- Be clean. Clean the workbench, pipetmen, racks, and centrifuges with RNase Away or something equivalent.
- Wear gloves and change when contaminated. Contamination will occur when gloves contact hair, face, skin, or the floor.
- Keep everything on ice unless the protocol indicates otherwise.
- Centrifuge components to the bottom of the tube before use, after mixing, after use, and after incubation steps.
- Do not vortex enzymes. Gently flick the tube or pipet up and down to mix samples.
- Avoid touching the walls of the tube when pipetting.
- Use a new pipette tip for each new dip.
- Dispense solutions from a pipet to the bottom of the tube, or into the liquid at the bottom of the tube when setting up reactions.
- Only remove 1.5 ml centrifuge tube and PCR tubes from their package while wearing gloves. Reseal the tube package after tubes are removed.

#### Assembly of CRISPR Oligos A + B into a Transcription Template

1. For synthesis of the gRNA from Oligo A and B, make a 100 μM freezer stock and 1 μM working stock for each oligo. All oligos are described in Section A starting on page 7.
2. Centrifuge ordered oligos briefly <u>before</u> opening, to move all dried DNA flakes to the bottom of the tube.
3. Add a volume (x μL) of RNase-free water to make a 100 μM stock. The tubes should be labeled with the gene name as well as the number of nmol in the tube. The amount of water to be added will need to calculated based on the nanomoles of material contained within.
4. Vortex for 30 seconds.
5. Centrifuge briefly.
6. Make a 100-fold dilution of each 100 μM stock Oligo A and B in separate 1.5 ml tubes.

a. Label one 1.5 mL centrifuge tube per Oligo A with name of oligo, date, and “1 μM” to indicate working stocks.

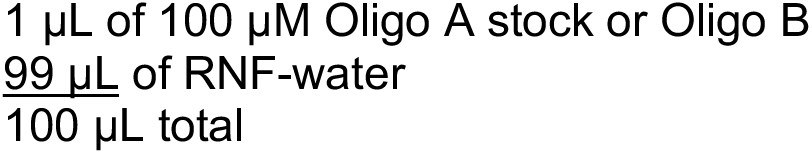
b. Vortex.
c. Briefly centrifuge.
d. Store all stocks in freezer at −20 °C for long-term storage.
7. Set up the following reaction in PCR tubes. The next two steps will generate a short segment of DNA (gDNA or guideDNA Template) which will be used as a template for synthesis of RNA:

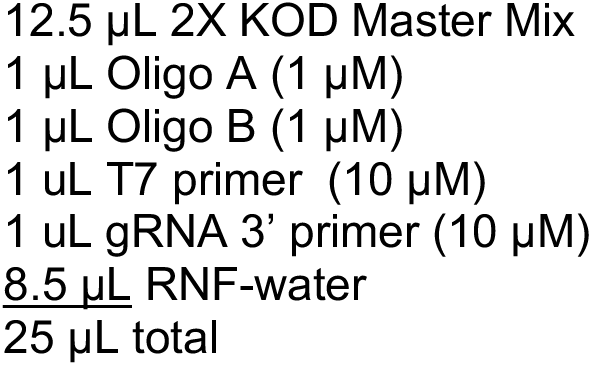
8. Run PCR under the following conditions:

Denature at 98°C for 2 minutes
Denature at 98 °C for 30 sec.
Anneal at 50 °C 30 sec.
Extend at 70 °C 30 sec.
Go to (step 2) nine times.
Extend at 70 °C 2 min
Hold 4 °C forever.
9. Run 1.2% agarose gel in 1X TAE to check that the template was synthesized:

a. Remove 3 μL of the reaction and place in a 1.5 ml tube.
b. Mix in 1 μL of 6x loading buffer.
c. Load all 4 μL of the sample on the gel. Run the gel at 125 V for 30 minutes. Be sure to load a molecular weight marker.
d. Check on the transilluminator and image the gel.
e. A single 120 bp band should be detected when 3 μL is loaded on gel.

#### In vitro transcription (IVT) using the gRNA template

1. Use the Ambion T7 Megascript Kit for transcription reagents, but follow the instructions below.
2. Thaw the T7 10X Reaction Buffer and RNF-water at room temperature, and thaw the ribonucleotides solutions on ice.
3. Vortex the T7 10X Reaction Buffer to make sure all DTT is solubilized. No white flecks should be visible.
4. Microcentrifuge all reagents briefly before opening to prevent loss of reagents and/or contamination by materials that may be present around the rim of the tube(s).
5. Keep the T7 Enzyme Mix on ice or in a −20 °C block during assembly of the reaction.
6. Make a master mix for each reaction. Assemble the reaction at room temperature on the bench. Add reagents from largest to smallest volume, adding the 10X Reaction Buffer second to last and the T7 Enzyme Mix last. *Note:* Components in the transcription buffer can lead to precipitation of the template DNA if the reaction is assembled on ice. If the reaction precipitates, the synthesis reaction will not fully occur.
7. Reagent list:

μL of RNF-water
μL of gDNA template (100 to 500 ng total)
μL of NTP (1 µl of each; A, U, C, G)
μL of 10x transcription buffer – must be fully resuspended at room temp
μL of T7 polymerase enzyme mix
8. Incubate at 37 °C for 4 to 16 hours. Longer incubations result in considerably better yields.
9. Add 1 μL of Turbo DNAse and incubate for 15 min at 37 °C. This will digest the template DNA in the sample.
10. Optional quality control step: Run 2 μL of sample on a 1.2% gel in 1X TAE.

a. Clean the gel box, comb and tray with RNase Away, rinse with DI water.
b. Remove 2 µl of sample into a clean 1.5 ml (Keep RNA on ice!)
c. Add 3 μL of RNF-water and 5 uL of Ambion RNA loading buffer with formamide.
d. Vortex briefly.
e. Spin down samples briefly.
f. Run all of this mixture on a 1.2% agarose gel/1X TAE, at 100 V for 1 hour.
g. Image gel. 2 bands should be visible at ~100 and 120 bp.

#### Purification of guide RNA

1. Use the miRNeasy Qiagen kit for purification of gRNAs according to the manufacturer’s instructions.
2. After Purification verify presence of RNA by running a 1.2% gel in 1X TAE.
3. Clean the gel box, comb and tray with RNase Away, rinse with DI water. Run on a 1.5% agarose gel/1X TAE, at 100 V for 1 hour as above.
4. Image gel. 2 bands should be visible at ~100 and 120 bp.
5. Nanodrop the RNA sample to determine the concentration.
6. Store RNA at −20 °C.

#### Preparation of SpCas9 mRNA

1. Digest ~5-10 μg pT3TS-nCas9n plasmid with Xba1 (plasmid Addgene #46757 (Jao et al., 2013)).
2. Purify digested DNA with Qiagen PCR cleanup kit or Promega PureYield Plasmid Miniprep System. Elute in RNF-water.
3. Run 100-500 ng on 1.2% agarose gel in 1X TAE to confirm the plasmid is linearized.
4. Use 100 ng to 1 μg DNA as template for in vitro transcription reaction.
5. Use mMESSAGE mMACHINE T3 kit Life Technologies (AM1348) and follow the instructions in the manual.
6. Use the miRNeasy Qiagen kit for purification of nCas9n mRNA according to the manufacturer’s instructions.
7. Verify mRNA integrity by mixing 1 uL of purified Cas9, 4 μL of RNF water, 5 μL glyoxl dye (Ambion).
8. Heat mixture at 50 °C for 30 minutes, then place on ice.
9. Clean the gel box, comb and tray with RNase Away, rinse with DI water.
10. Run all 10 μL of RNA mixture on 1.2% agarose gel in 1X TAE at 100 V for 1 hour as above. One band should be visible at 4.5 kb.
11. Nanodrop the RNA sample to determine the concentration. Concentrations between 0.45 and 1 μg/μL are expected.
12. Aliquot and store RNA at −80 °C.

### C. Injection of sgRNA and spCas9 mRNA

The injections here are designed to deliver 25 pg of gRNA and 300 pg of Cas9 mRNA in 2 nL of fluid to embryos at one-cell stage.

Injection trays are cast with 1.2% agarose with 1X embryo media (Zebrafish Book; zfin.org) in polystyrene petri dishes (Fisher No. FB0875713). Injection trays can be used multiple times and stored at 4*C for up to three weeks between use.

1. Trays are pre-warmed to 28.5 °C prior to injection by placing them in the 28.5 °C incubator. Try to mitigate tray cooling while not in use.
2. Glass needles are pulled from Kwik-Fil borosilicate glass capillaries (No. 1B100-4) on a Flaming/Brown Micropipette puller (Model P-97). Injection samples are made to contain the following diluted in RNF water or injection buffer (final concentration: 12.5 mM HEPES pH 7.5, 25 mM Potassium Acetate, 37.5 mM Potassium Chloride, 0.0125 % glycerol, 0.025 mM DTT ph 7.5)

a. ng/μL of genomic gRNA
b. 150 ng/μL of mRNA for Cas9
3. Needles are loaded with 1.5 to 2.5 μL of sample, and then loaded onto a micro-manipulator attached to a micro injector (Harvard Apparatus PLI - 90) set to 30-40 PSI with an injection time of 200 msec.
4. Needles are calibrated by breaking the end of the tip off with sterile tweezers, ejecting 10 times to produce a droplet of fluid, collecting the droplet into a 1 μL capillary tube (Drummond No. 1-000-0010), and measuring the distance from the end of the capillary to the meniscus of the droplet. This distance is converted to volume (where 1 mm = 30 nL) and adjusted to achieve an effective injection volume of 2 nL. Volumes are adjusted by changing the injection time. There is a linear relationship between volume and time at a set pressure. Avoid injection times less than 100msec and over 400 msec.
5. One cell embryos that have been collected from mating cages are pipetted from collection petri dishes to the wells on the injection tray.
6. Use the micro-manipulator and microscope to pierce the one-cell of embryos on the injection tray at an angle of 30° with the needle. Inject 2 nL of sample in the one-cell near the center of the cell-yolk boundary.
7. After embryos have been injected, wash them from the injection tray into a clean petri dish with embryo media.
8. Keep 20 - 40 embryos separate as uninjected controls. Treat and score the control embryos in the same way as the injected embryos.
9. At 3 - 5 hrs post injection remove any unfertilized or dead embryos from the dishes. This will prevent death of the still developing embryos.

### D. Testing for indel production/mutagenesis

#### Phenotypic scoring of embryos

1. The gRNA itself may be toxic to the developing embryos. Injection toxicity can be estimated by the number dead embryos from a round of injection compared to the un-injected control dish. Count and remove any brown/dead embryos from injected and un-injected dishes. If there are significantly more dead embryos in the injected dish then the guide may be toxic, impure, or very effective at disrupting a required gene. Reducing the amount of guide or Cas9 mRNA injected may help reduce toxicity.
2. Score and document embryonic phenotypes on days 1 - 4 post fertilization (dpf). Under a dissection microscope examine the un-injected controls and injected embryos, and sort the embryos into categories.
3. Scoring categories

- -Severe- These embryos have some parts that look like a control embryos, but are missing key features. Examples: embryos that lack their head, eyes, or tail, or embryos that have an unnaturally contorted shape or are asymmetric.
- -Mild- These embryos appear mostly normal, but have slight defects such as small eyes, pericardial edema, shortened trunk/tail, or curled/twisted tails.
- -Normal- appears normal and similar to controls.

#### Digestion of embryos for isolation of genomic DNA for mutagenesis analysis

Genomic DNA (GDNA) can be isolated from zebrafish embryos aged between 1 and 5 dpf using this protocol. Embryos can be analyzed as individuals or as pools (maximum 5) from the same injection.

1. Dechorionate embryos, if they have not emerged from the chorion.
2. It is recommended to screen a minimum of 3 embryos from each scoring category for mutagenesis. Place each embryo, including controls, into separate PCR tubes. Remove as much of the fish water as possible. If needed, spin briefly and remove additional water.
3. Add 20 μL of 50 mM NaOH <u>per embryo</u>.
4. Heat the embryos at 95°C in a thermocycler for 15 minutes.
5. Vortex samples for 10 seconds. Be sure that the tubes are sealed to prevent sample loss while vortexing.
6. Spin samples down and heat for an additional 15 min at 95 °C in a thermocycler.
7. Vortex samples and then spin the tubes down again. The embryos should be completely dissolved.

7. Neutralize the samples by adding 1 μL of 1 M Tris pH 8.0 per 10 μL NaOH. Mix by vortexing then spin down.

8. Genomic DNA should be kept at 4 °C while in use and stored at −20°C.

#### Analysis of CRISPR/Cas9 mutagenesis efficiency at targeted gene locus

1. Set up the following PCR reactions for each tube of digested embryos using the primers designed at the end of section A, page 10.

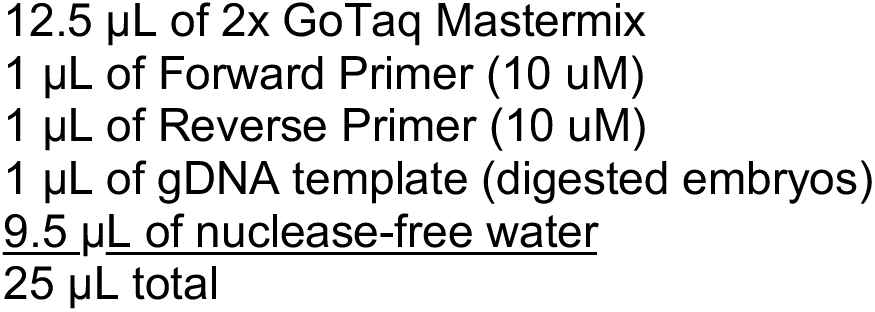
2. Vortex and briefly spin down the PCR reactions.
3. Run the following PCR program to amplify the targeted locus.

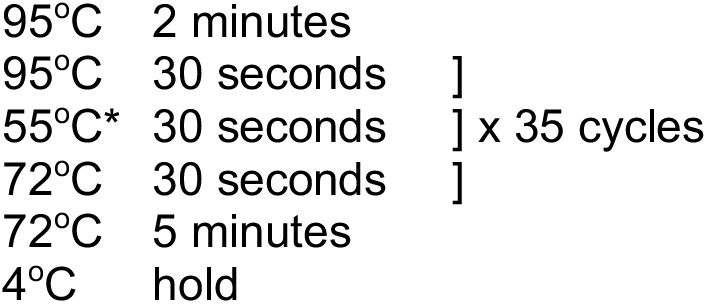

*if the primers were designed with higher or lower tm’s than the annealing temperature in line three, then that temperature will need to be adjusted to 2°C below the designed primer tm.
4. Run up to 7 μL of PCR product on a 3.0% agarose gel, 1X TAE, for 1 hr at 80-100V.
5. Analyze the gel for DNA bands that appear diffuse or different in size from the control lane. This indicates that the presence of indels in the gene of interest
6. Alternatively clone and sequence PCR products or sequence them directly to verify the presence of indels.

### E. Design short homology arms

Homology directed gene targeting allows the integration of exogenous DNA into the genome with precision to the base pair level. However, designing and cloning individual targeting vectors and homology arms for each gene of interest can be time consuming. The pGTag vector series provides versatility for ease of generation of knockout alleles (Figure 3). The vectors contain BfuAI and BspQI type II restriction enzymes for cloning of short homology arms (24 or 48 bp) using Golden Gate cloning. The pGTag vectors require in-frame integration for proper reporter gene function. The reporter gene consists of several parts. First, a 2A peptide sequence causes translational skipping, allowing the following protein to dissociate from the locus peptide. Second and third, eGFP, TagRFP, or Gal4VP16 coding sequences for the reporter protein have a choice of sequence for localization domains, including cytosolic (no) localization, a nuclear localization signal (NLS), or a membrane localization CAAX sequence. Finally translation is terminated by one of two different polyadenylation sequences (pA); a β-actin pA from zebrafish or the SV40pA.

For many genes, the signal from integration of the report protein is too weak to observe. In these cases the Gal4VP16 vector allows for amplification of the report to observable expression levels in F0s and subsequent generations. A 14XUAS/RFP Tol2 plasmid is provided to make a transgenic line for use with the Gal4VP16 vector.

Sequence maps for these plasmids can be downloaded at www.genesculpt.org/gtaghd/

**Figure 3.**
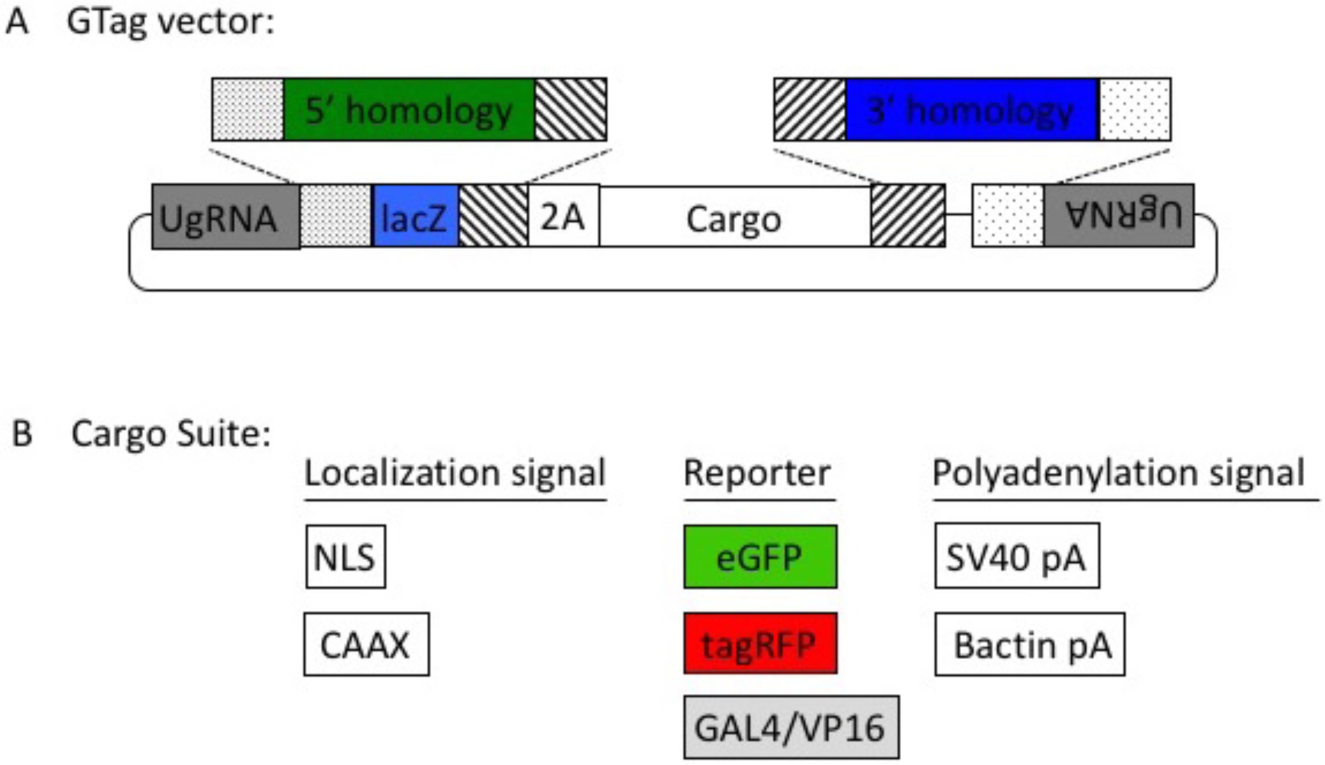
The pGTag vectors allow one step cloning of homology arms.

All vectors can be obtained through Addgene (www.addgene.org). Because the pGTag plasmids contain repeated sequences, they may be subject to recombination in certain strains of bacteria. **It is strongly recommended that they are propagated at 30°C to reduce the possibility recombination.**

The web tool, GTagHD www.genesculpt.org/gtaghd/, allows for quick design of cloning ready homology arm oligos for a gene of interest.

To use the tool, choose the “Submit Single Job” tab. Follow the instructions in the tab.

There should be 4 oligos (two pairs that will be annealed) generated that should be ordered for cloning. If there are any problems with the sequences and values that were entered, the web page will display the errors and give advice on how to fix them.

The following protocol describes how to design homology arm oligos manually:

*Note* In the following section when orientation words are used, they are used in the context of the reading frame of the genetic locus of interest. Example: A 5’ template strand CRISPR means that the target site for the CRISPR is on the template strand at the locus and is toward the 5’ end of the gene. Upstream homology domains are 5’ of the CRISPR cut and downstream homology domains are 3’ of the cut with respect to the gene being targeted. Also note: Upper case and lower case bases are not specially modified; they are typed the way they are as a visual marker of the different parts of the homology arms.

#### For the Upstream Homology Domain

1) Open the sequence file for the gene of interest and identify the CRISPR site. (In this example it is a Reverse CRISPR target in Yellow, the PAM is in Orange, coding sequence is in purple)

Copy the 48 bp 5’ of the CRISPR cut (the highlighted section below) into a new sequence file; this is the upstream homology.

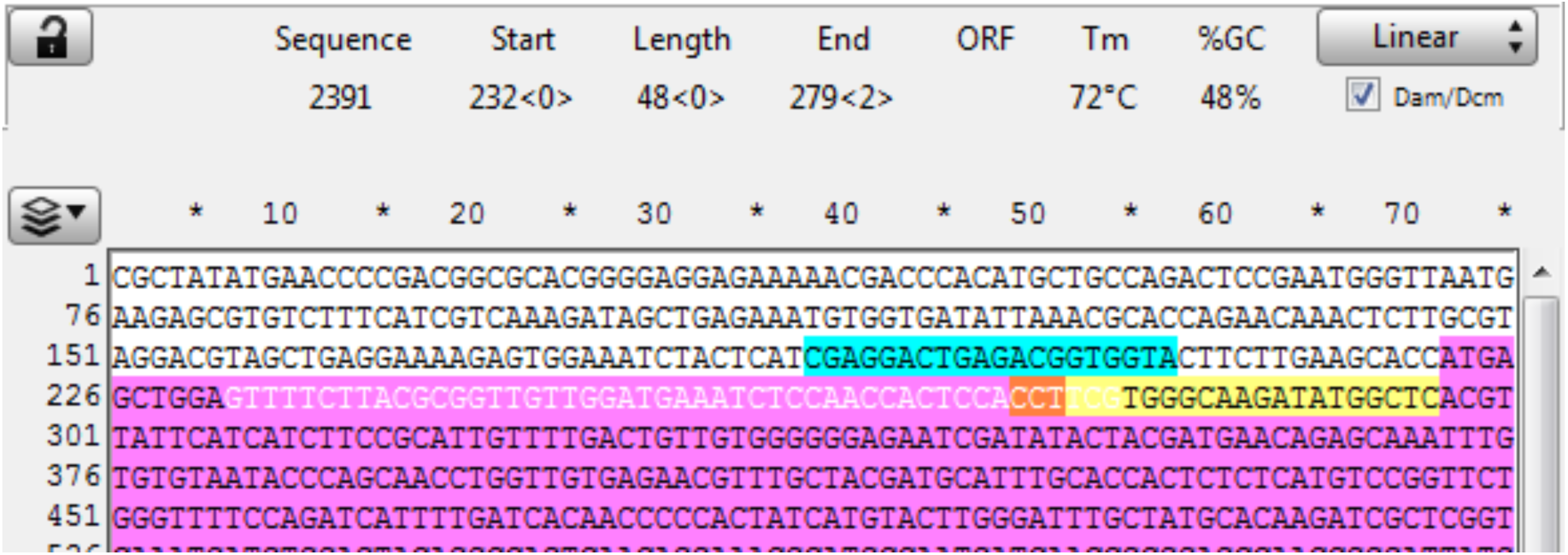

2) Observe the next three bases immediately upstream of the 48 bp of homology, and pick a base not present to be the 3 bp spacer between the homology and the Universal PAM in the vector. (Here the three bases are “GGA” so “ccc” was chosen for the spacer)

Add the spacer to the new file 5’ (in front) of the homology, see below. The spacer acts a non-homologous buffer between the homology and the eventual 6 bp flap from the universal guide sequence that will occur when the cassette is liberated and may improve intended integration rates over MMEJ events.

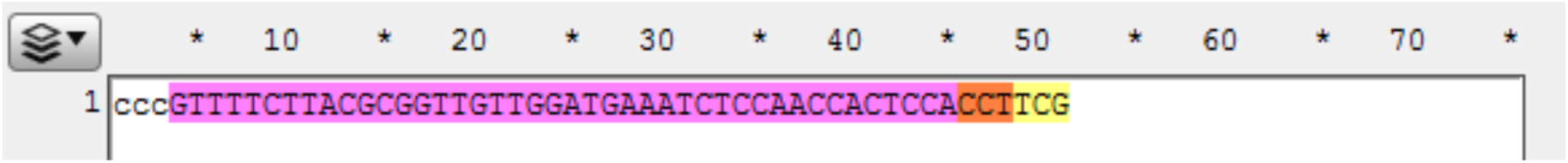

3) Determine where the last codon is in the homology. Here the 3’ G in the homology domain is the first base in the codon cut by this CRISPR target. Complete the codon by adding the remaining bases (called padding on GTagHD) for that codon from your sequence to ensure your integration event will be in frame.

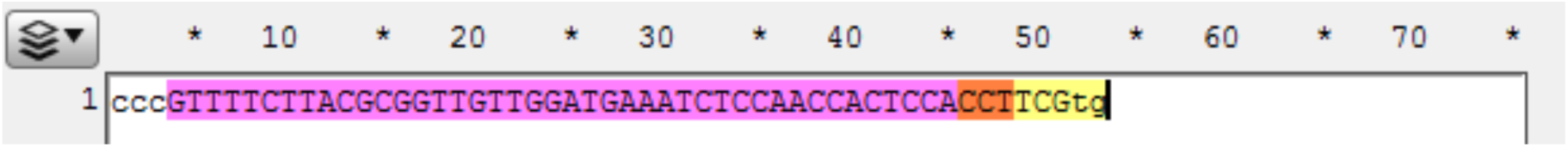

4) Add the BfuAI enzyme overhang sequences for cloning, to the ends of the homology domain. 5’-GCGG and 3’-GGAT. (Here both overhangs are added to prevent errors in copying sequence for the oligos in the next two steps.)

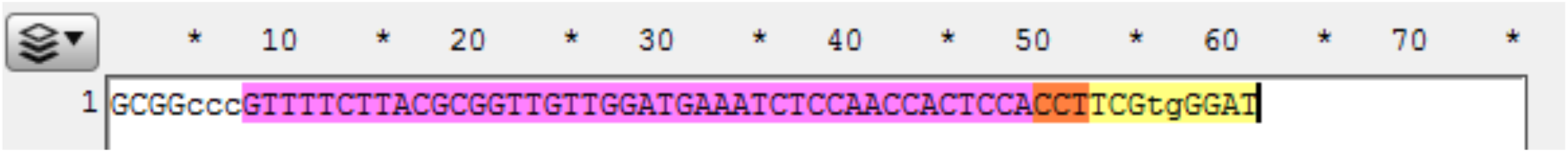

5) The Upstream Homology Oligo A will be this sequence from the beginning to the end of the last codon (see highlighted below). Copy and paste this sequence into a new file and save it. In this example this oligo sequence is 5’- GCGGcccGTTTTCTTACGCGGTTGTTGGATGAAATCTCCAACCACTCCACCTTCGtg-3’.

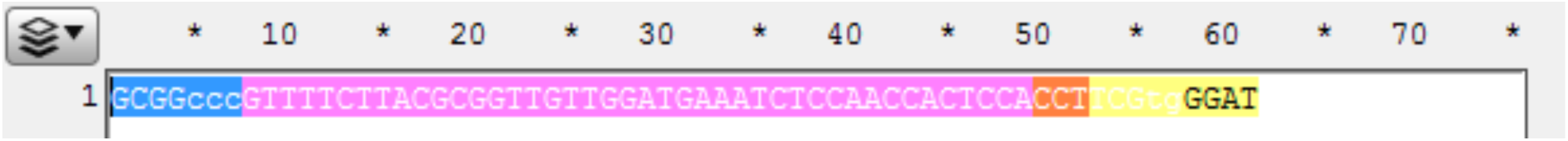

6) The Upstream Homology Oligo B will be the reverse compliment of this sequence from beginning of the spacer to the end of the sequence (see highlighted below). Copy the reverse compliment, paste it into a new file, and save it. In this example this oligo sequence is 5’- ATCCcaCGAAGGTGGAGTGGTTGGAGATTTCATCCAACAACCGCGTAAGAAAACggg-3’.

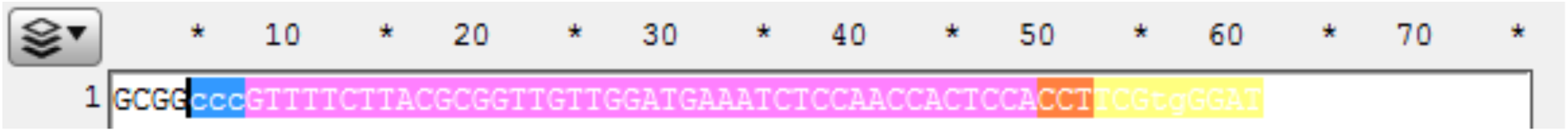

#### For the Downstream Homology Domain

7) Open sequence file for the gene of interest and identify the CRISPR site. (Reverse CRISPR target in Yellow, PAM in Orange, coding sequence is in purple) Copy the 48 bp 3’ of the CRISPR cut into a new sequence file; this is the downstream homology.

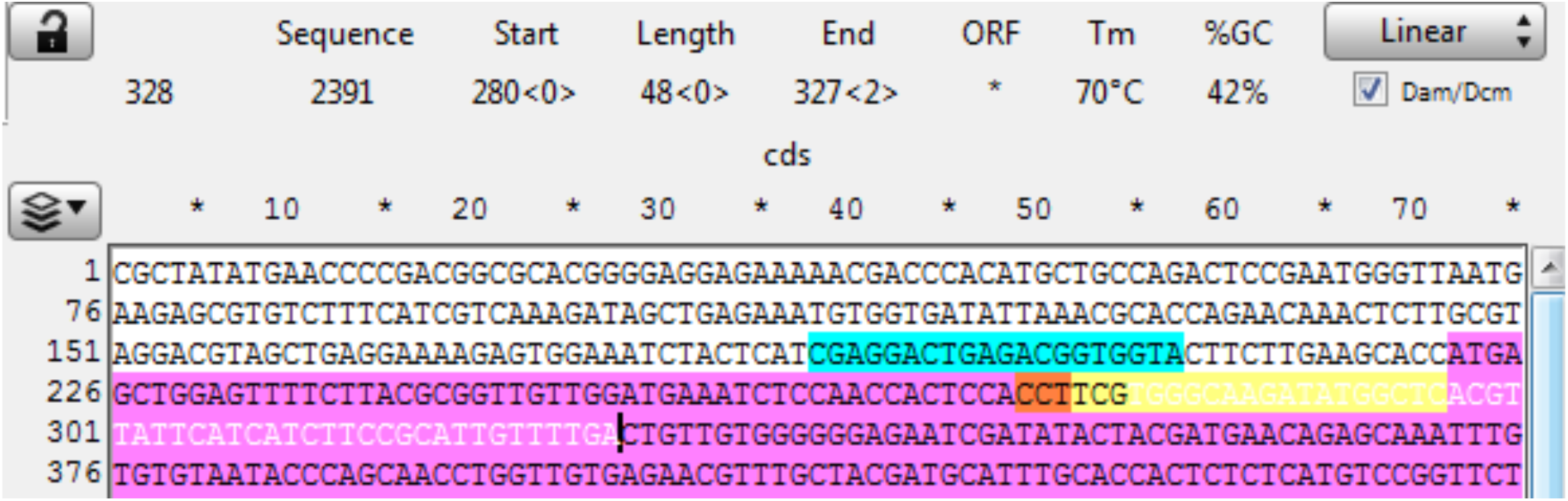

8) Observe the next three bases downstream of the 48 bp of homology, and pick a base not present to be the 3 bp spacer between the homology and the Universal PAM in the vector. (Here the bases are “CTG” so “aaa” was chosen for the spacer.)

Add the spacer to the new file 3’ of (after) the homology.

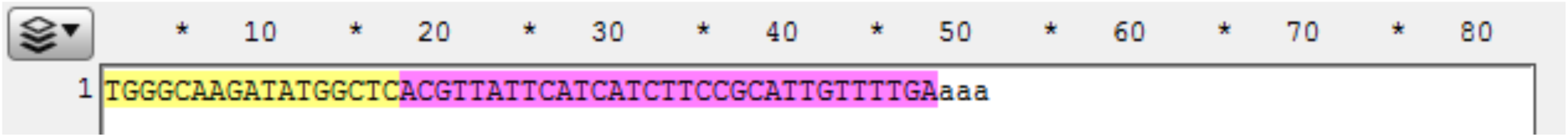

9) Add the BspQI enzyme overhang sequences for cloning, to the ends of the homology domain. 5’-AAG and 3’-CCG. (Here both overhangs are added to prevent errors in copying sequence for the oligos in the next two steps.)

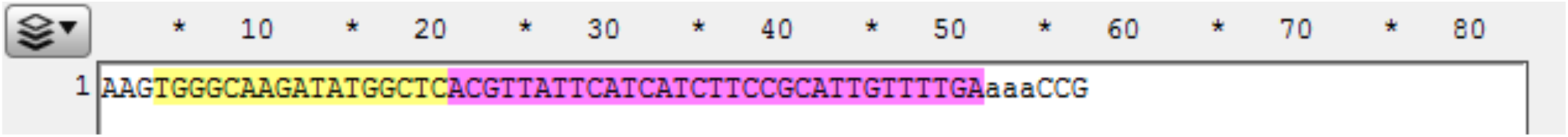

10) The Downstream Homology Oligo A will be this sequence from the beginning of the sequence to the end of the spacer (see highlighted below). In this example this oligo sequence is 5’-AAGTGGGCAAGATATGGCTCACGTTATTCATCATCTTCCGCATTGTTTTGAaaa-3’.

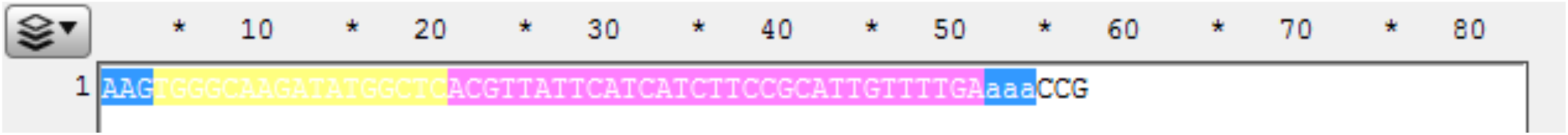

11) The Downstream Homology Oligo B (will be the reverse compliment of this sequence from the beginning of the homology to the end of the sequence (see highlighted below). In this example this oligo sequence is 5’- CGGtttTCAAAACAATGCGGAAGATGATGAATAACGTGAGCCATATCTTGCCCA-3’

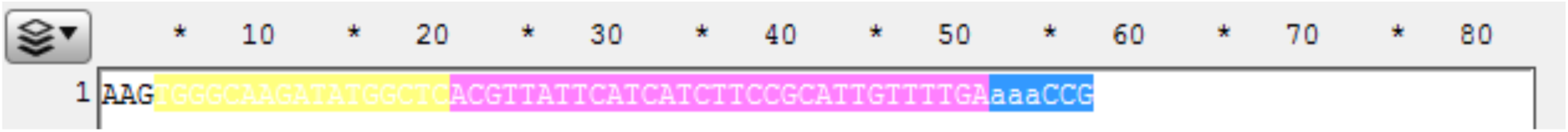

### F. One Pot Cloning of Homology Arms into pGTag Vectors

**Note if the homology arm oligos contain either the sequence “5’-ACCTGC-3’” or “5’-GAAGAGC-3’” (or their compliments) the cloning reaction will be less efficient.

**\*Note some sequences just don’t work very well.** Ligation is more efficient with annealed homology arms and the purified ~1.2 kb and ~2.4kb fragments from vectors that have been digested with BfuAI and BspQI. If problems are encountered, one homology arm can also be cloned sequentially.

1. Homology Arm Annealing Anneal upstream and downstream homology oligo pairs separately:

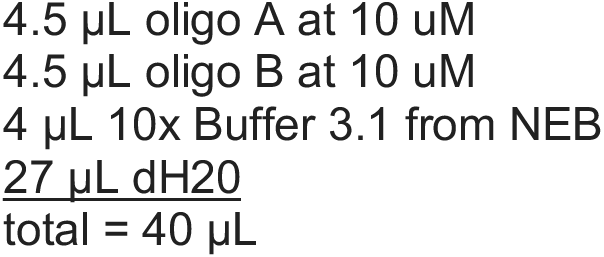

Incubate at 98°C for 4 min, 98°C 45 sec x 90 steps decrementing temp 1°C/cycle, 4°C hold (Alternatively heat in 95-98°C water for 5 minutes, and then remove the boiling beaker from the heat source and allow it to cool to room temp for 2 hours, before placing samples on ice.)
2. 1-Pot Digest Assemble the following:

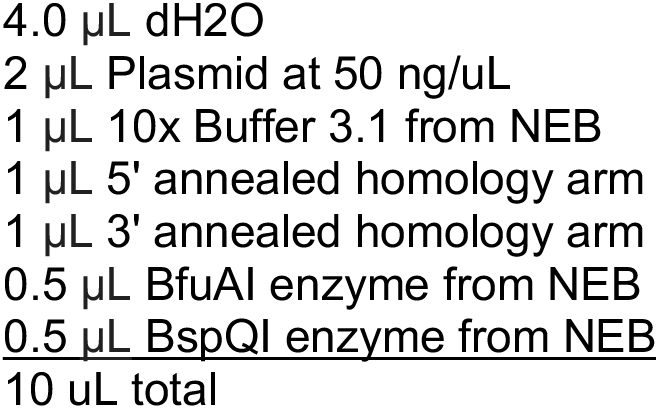

Incubate at 50°C for 1 hr, place on ice.
3. Ligation Add the following:

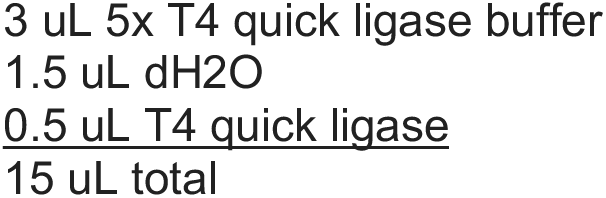

Incubate 8-10 min at room temperature (to overnight). Store at −20 °C,
4. Transformation

a. On ice, thaw 1 (one) vial competent cells (50 μL) for every 2 ligation reactions. (approx. 5 min). It is recommended to use NEB Stable Competant E. coli (C3040H) cells to limit recombination.
b. While cells are thawing, label the microcentrifuge tubes for each ligation and put on ice.
c. Once the cells are thawed, use a pipette to transfer 25 μL of the competent cells into each labeled tube.
d. Add 1.5 μL of a ligation reaction into competent cells to transform.

a. Amount of ligation reaction added should be less than 5% of volume of competent cells.
e. Mix by tapping the tube several times or gently mixing with the pipet tip.

a. Do NOT mix by pipetting, this will lyse the cells.
f. Incubate on ice for 5 to 20 minutes.
g. Heat shock the cells by submerging the portion of the tube containing the cells in a 42°C water bath for 40 - 50 seconds.
h. Incubate cells on ice for 2 minutes.
i. Add 125 μL of room temperature LB to each transformation.
j. Incubate cells at 30°C for 1-1.5 hour(s) in a shaking incubator.
k. While the transformed cells are recovering, spread 40 μL of X-Gal solution, and 40 μL IPTG 0.8 M on LB Kanamycin selection plates.

a. X-Gal is lethal to cells while wet, it is recommended to first label the plates and then place them in a 30°C incubator to dry.
l. After recovery and the X-Gal is dry, Plate 150 μL of each transformation on the corresponding correctly labeled plate.
m. Incubate plates overnight at 30°C.
5. Growing colonies Pick 3 white colonies from each plate and grow in separate glass culture tubes with 3 mL LB/Kanamycin. Or to pre-screen colonies by colony PCR:

a. Pick up to 8 colonies with a pipet tip and resuspend them in separate aliquots of 5 μL dH2O. Place the tip in 3 ml of LB/Kan, label, and store at 4°C.
b. Make a master mix for your PCR reactions containing the following amounts times the number of colonies you picked.

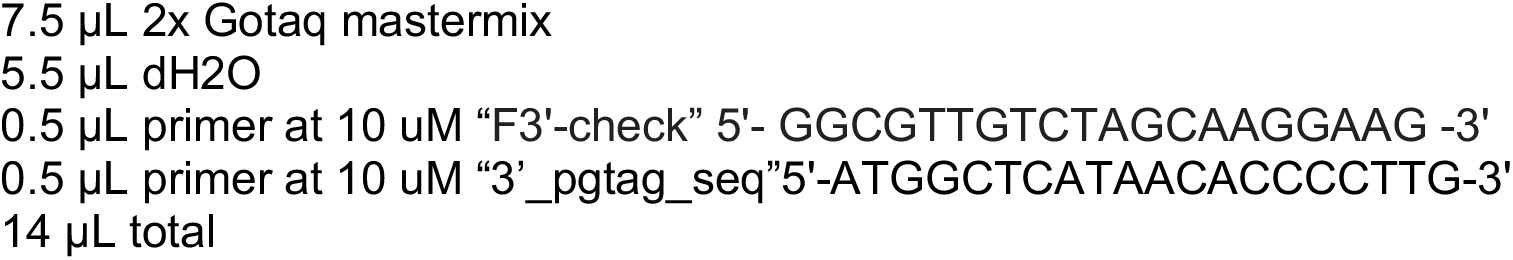
c. Aliquot 14 μL of mixed master mix into separate labeled PCR tubes.
d. Add 1 μL of colony to each reaction as template.
e. or 20 ng purified plasmid as control.
f. Cycle in a thermocycler

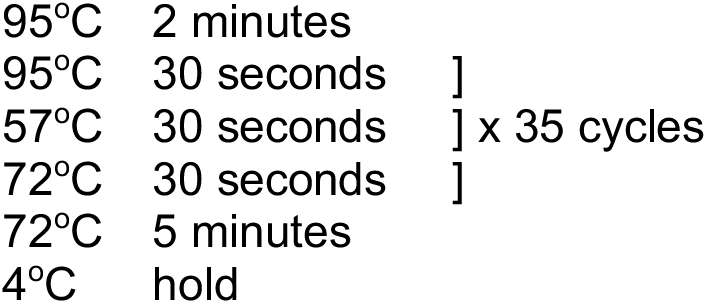
g. Run 5 μL of PCR product on a 1% agarose gel. You should get bands that are a different size than the control.
6. Mini Prep Cultures Follow Qiagen Protocol
7. Sequencing of Plasmids The 5’ homology arm can be sequenced by the 5’_pgtag_seq primer: 5’-GCATGGATGTTTTCCCAGTC-3’. The 3’ homology arm can be sequenced with the “3’_pgtag_seq”primer: 5’-ATGGCTCATAACACCCCTTG-3’.

### G. Injection of GeneWeld Reagents (spCas9 mRNA, Universal sgRNA (UgRNA), genomic sgRNA and pGTag homology vector) into 1-cell zebrafish embryos

#### Prepare and collect the following reagents for injection

1. Prepare nCas9n mRNA from pT3TS-nCas9n (Addgene #46757 from (Jao et al., 2013)) as described above (page 14).
2. Synthesize UgRNA and purify as described above (page 11) using the following oligo A: 5’-TAATACGACTCACTATAGGGAGGCGTTCGGGCCACAGGTTTTAGAGCTAGAAATAGC-3’ Corresponding to the universal target sequence: GGGAGGCGTTCGGGCCACAG Alternatively, the UgRNA can be directly ordered form IDT and resuspended in RNF water. 5’-GGGAGGCGUUCGGGCCACAGGUUUUAGAGCUAGAAAUAGCAAGUUAAAAUAAGG CUAGUCCGUUAUCAACUUGAAAAAGUGGCACCGAGUCGGUGCGGAUC-3’
3. The pGTag homology vectors should be purified a second time prior to microinjection under RNase free conditions with the Promega PureYield Plasmid Miniprep System beginning at the endotoxin removal wash and eluted in RNF water.

#### Embryo Injections for Integration of pGTag vectors

Injections are performed as previously described in 2 nl per embryo with the addition of the UgRNA and targeting pGTag DNA.

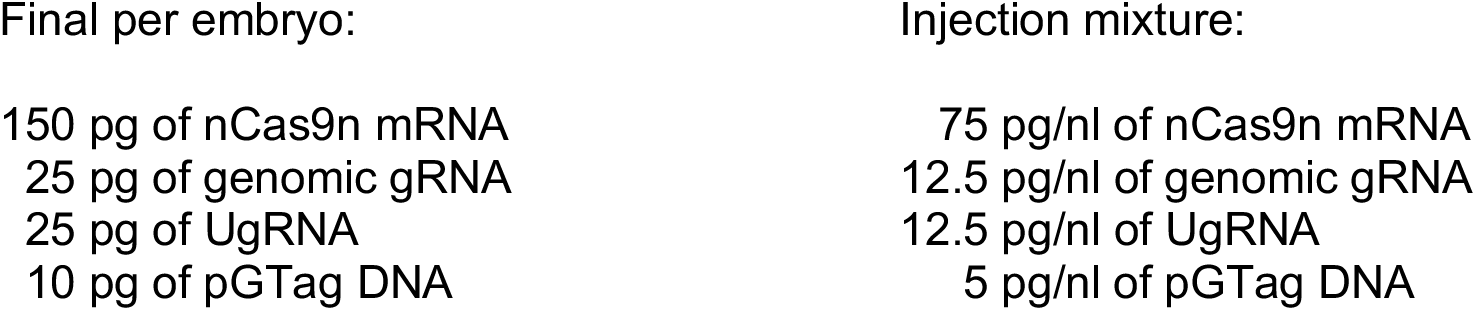

### H. Examine embryos for fluorescence and junction fragments

Embryos are examined for fluorescence under a Zeiss Discovery dissecting microscope with a 1X objective at 70-100X magnification. If weak signals are observed, embryos are manually dechorionated, and viewed on glass depression well slides. If no or weak signals were observed, Gal4VP16 integrations are attempted in a 14XUAS-RFP background. Embryos displaying widespread fluorescence in expression domains consistent with the targeted gene are examined for junction fragments or raised to adulthood for outcrossing.

F0 Junction fragment analysis between the genomic locus and the targeting vector is carried out by isolating DNA from embryos followed by PCR. The following primers are used for junction fragment analysis and must be paired with gene specific primers (5’ to 3’):

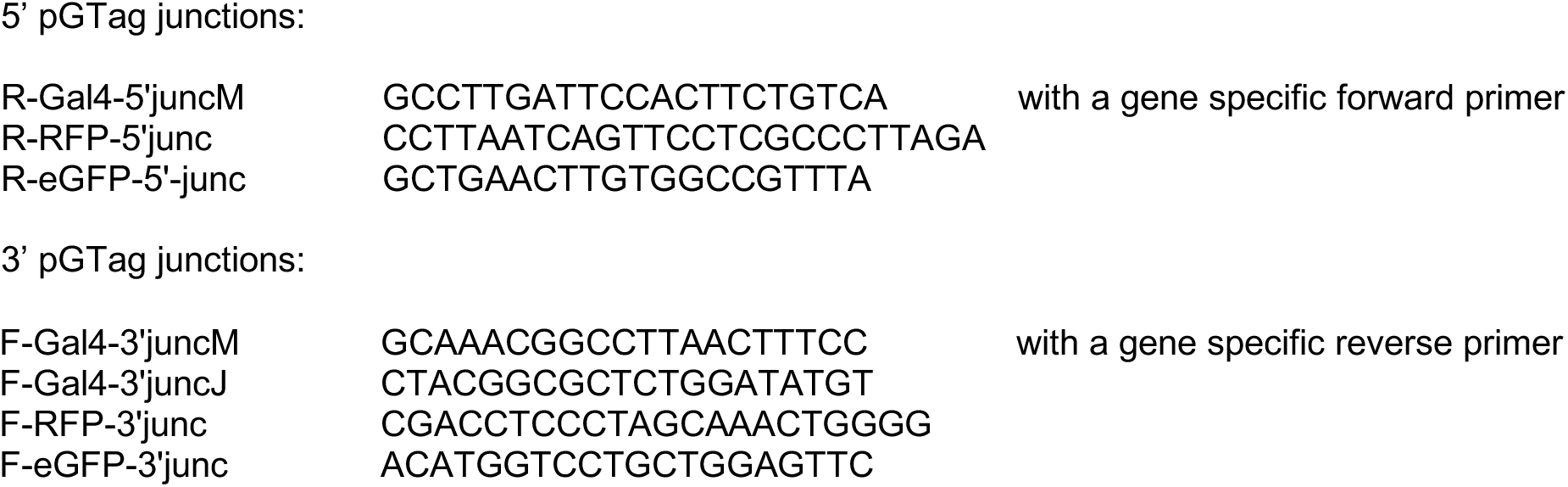

PCR amplification of junction fragments can be a result of artifacts (Won and Dawid, 2017), so it is important to carryout control amplifications with injected embryos that lack the genomic gRNA. F0 analysis by PCR of junction fragments is carried out to examine correct targeting. F-Gal4-3’juncM and F-Gal4-3’juncJ are two alternate primers for amplification of junction fragments from the Gal4 cassette due to gene specific mis-priming depending on the target loci.

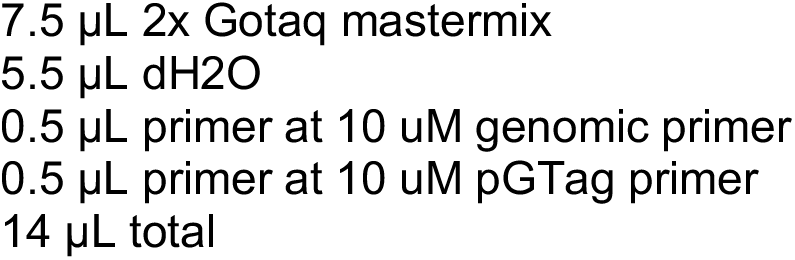

1. Aliquot 14 μL of mixed master mix into separate labeled PCR tubes.
2. Add 1 μL of genomic DNA to each reaction as template.
3. Cycle in a thermocycler with the following steps:

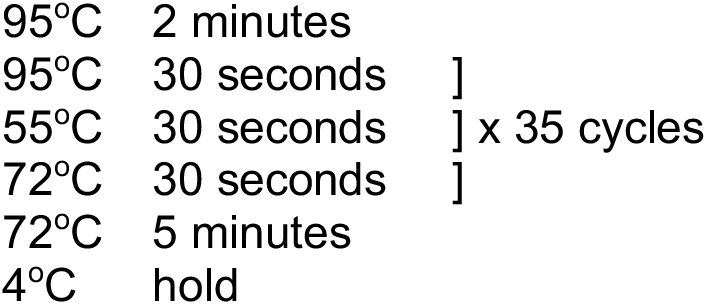
4. Run 5 μL of PCR product on a 1.2 % agarose gel in 1XTAE. Putative junction fragments should give bands that are of predicted size.

F0 animals that are positive for the reporter gene are raised to adults then outcrossed and examined for fluorescence as above. The Gal4VP16 system can lead to silencing resulting in mosaic patterns in F1 embryos. F1 embryos displaying fluorescence are examined for junction fragments as above, raised to outcross to make F2 families or sacrificed at 3 weeks post fertilization for Southern-Blot analysis of integrations. F0 and F1 identified fish can be incrossed or backcrossed to get an initial impression of the homozyogous phenotypes. It is recommended that lines are continuously outcrossed once established.

## References

Aida, T., Nakade, S., Sakuma, T., Izu, Y., Oishi, A., Mochida, K., Ishikubo, H., Usami, T., Aizawa, H., Yamamoto, T., et al. (2016). Gene cassette knock-in in mammalian cells and zygotes by enhanced MMEJ. BMC Genomics 17, 979.

Ata, H., Ekstrom, T.L., Martinez-Galvez, G., Mann, C.M., Dvornikov, A.V., Schaefbauer, K.J., Ma, A.C., Dobbs, D., Clark, K.J., and Ekker, S.C. (2018). Robust activation of microhomology-mediated end joining for precision gene editing applications. PLoS Genet 14, e1007652.

Balciuniene, J., Nagelberg, D., Walsh, K.T., Camerota, D., Georlette, D., Biemar, F., Bellipanni, G., and Balciunas, D. (2013). Efficient disruption of Zebrafish genes using a Gal4-containing gene trap. BMC Genomics 14, 619.

Bedell, V.M., Wang, Y., Campbell, J.M., Poshusta, T.L., Starker, C.G., Krug, R.G., 2nd, Tan, W., Penheiter, S.G., Ma, A.C., Leung, A.Y., et al. (2012). In vivo genome editing using a high-efficiency TALEN system. Nature 491, 114–118.

Beumer, K.J., Trautman, J.K., Bozas, A., Liu, J.L., Rutter, J., Gall, J.G., and Carroll, D. (2008). Efficient gene targeting in Drosophila by direct embryo injection with zinc-finger nucleases. Proc Natl Acad Sci U S A 105, 19821–19826.

Carlson, D.F., Tan, W., Lillico, S.G., Stverakova, D., Proudfoot, C., Christian, M., Voytas, D.F., Long, C.R., Whitelaw, C.B., and Fahrenkrug, S.C. (2012). Efficient TALEN-mediated gene knockout in livestock. Proc Natl Acad Sci U S A 109, 17382–17387.

Ceccaldi, R., Rondinelli, B., and D’Andrea, A.D. (2016). Repair Pathway Choices and Consequences at the Double-Strand Break. Trends Cell Biol 26, 52–64.

Clark, K.J., Balciunas, D., Pogoda, H.M., Ding, Y., Westcot, S.E., Bedell, V.M., Greenwood, T.M., Urban, M.D., Skuster, K.J., Petzold, A.M., et al. (2011). In vivo protein trapping produces a functional expression codex of the vertebrate proteome. Nat Methods 8, 506–515.

Clarke, R., Heler, R., MacDougall, M.S., Yeo, N.C., Chavez, A., Regan, M., Hanakahi, L., Church, G.M., Marraffini, L.A., and Merrill, B.J. (2018). Enhanced Bacterial Immunity and Mammalian Genome Editing via RNA-Polymerase-Mediated Dislodging of Cas9 from Double-Strand DNA Breaks. Mol Cell 71, 42–55 e48.

Conway, A.B., Lynch, T.W., Zhang, Y., Fortin, G.S., Fung, C.W., Symington, L.S., and Rice, P.A. (2004). Crystal structure of a Rad51 filament. Nat Struct Mol Biol 11, 791–796.

Geurts, A.M., Cost, G.J., Freyvert, Y., Zeitler, B., Miller, J.C., Choi, V.M., Jenkins, S.S., Wood, A., Cui, X., Meng, X., et al. (2009). Knockout rats via embryo microinjection of zinc-finger nucleases. Science 325, 433.

Grzesiuk, E., and Carroll, D. (1987). Recombination of DNAs in Xenopus oocytes based on short homologous overlaps. Nucleic Acids Res 15, 971–985.

Halene, S., Wang, L., Cooper, R.M., Bockstoce, D.C., Robbins, P.B., and Kohn, D.B. (1999). Improved expression in hematopoietic and lymphoid cells in mice after transplantation of bone marrow transduced with a modified retroviral vector. Blood 94, 3349–3357.

Hasty, P., Rivera-Perez, J., Chang, C., and Bradley, A. (1991). Target frequency and integration pattern for insertion and replacement vectors in embryonic stem cells. Mol Cell Biol 11, 4509–4517.

He, M.D., Zhang, F.H., Wang, H.L., Wang, H.P., Zhu, Z.Y., and Sun, Y.H. (2015). Efficient ligase 3-dependent microhomology-mediated end joining repair of DNA double-strand breaks in zebrafish embryos. Mutat Res 780, 86–96.

Hisano, Y., Sakuma, T., Nakade, S., Ohga, R., Ota, S., Okamoto, H., Yamamoto, T., and Kawahara, A. (2015). Precise in-frame integration of exogenous DNA mediated by CRISPR/Cas9 system in zebrafish. Sci Rep 5, 8841.

Hoshijima, K., Jurynec, M.J., and Grunwald, D.J. (2016). Precise Editing of the Zebrafish Genome Made Simple and Efficient. Dev Cell 36, 654–667.

Ihry, R.J., Worringer, K.A., Salick, M.R., Frias, E., Ho, D., Theriault, K., Kommineni, S., Chen, J., Sondey, M., Ye, C., et al. (2018). p53 inhibits CRISPR-Cas9 engineering in human pluripotent stem cells. Nat Med 24, 939–946.

Kent, T., Chandramouly, G., McDevitt, S.M., Ozdemir, A.Y., and Pomerantz, R.T. (2015). Mechanism of microhomology-mediated end-joining promoted by human DNA polymerase theta. Nat Struct Mol Biol 22, 230–237.

Li, X.L., Li, G.H., Fu, J., Fu, Y.W., Zhang, L., Chen, W., Arakaki, C., Zhang, J.P., Wen, W., Zhao, M., et al. (2018). Highly efficient genome editing via CRISPR-Cas9 in human pluripotent stem cells is achieved by transient BCL-XL overexpression. Nucleic Acids Res.

McGrail, M., Hatler, J.M., Kuang, X., Liao, H.K., Nannapaneni, K., Watt, K.E., Uhl, J.D., Largaespada, D.A., Vollbrecht, E., Scheetz, T.E., et al. (2011). Somatic mutagenesis with a Sleeping Beauty transposon system leads to solid tumor formation in zebrafish. PLoS One 6, e18826.

Mehta, A., Beach, A., and Haber, J.E. (2017). Homology Requirements and Competition between Gene Conversion and Break-Induced Replication during Double-Strand Break Repair. Mol Cell 65, 515–526 e513.

Melby, A.E., Warga, R.M., and Kimmel, C.B. (1996). Specification of cell fates at the dorsal margin of the zebrafish gastrula. Development 122, 2225–2237.

Moreno-Mateos, M.A., Vejnar, C.E., Beaudoin, J.D., Fernandez, J.P., Mis, E.K., Khokha, M.K., and Giraldez, A.J. (2015). CRISPRscan: designing highly efficient sgRNAs for CRISPR-Cas9 targeting in vivo. Nat Methods 12, 982–988.

Nakade, S., Tsubota, T., Sakane, Y., Kume, S., Sakamoto, N., Obara, M., Daimon, T., Sezutsu, H., Yamamoto, T., Sakuma, T., et al. (2014). Microhomology-mediated end-joining-dependent integration of donor DNA in cells and animals using TALENs and CRISPR/Cas9. Nat Commun 5, 5560.

Ogura, E., Okuda, Y., Kondoh, H., and Kamachi, Y. (2009). Adaptation of GAL4 activators for GAL4 enhancer trapping in zebrafish. Dev Dyn 238, 641–655.

Orr-Weaver, T.L., Szostak, J.W., and Rothstein, R.J. (1981). Yeast transformation: a model system for the study of recombination. Proc Natl Acad Sci U S A 78, 6354–6358.

Rong, Y.S., and Golic, K.G. (2000). Gene targeting by homologous recombination in Drosophila. Science 288, 2013–2018.

Shin, J., Chen, J., and Solnica-Krezel, L. (2014). Efficient homologous recombination-mediated genome engineering in zebrafish using TALE nucleases. Development 141, 3807–3818.

Singleton, M.R., Wentzell, L.M., Liu, Y., West, S.C., and Wigley, D.B. (2002). Structure of the single-strand annealing domain of human RAD52 protein. Proc Natl Acad Sci U S A 99, 13492–13497.

Talbot, W.S., Trevarrow, B., Halpern, M.E., Melby, A.E., Farr, G., Postlethwait, J.H., Jowett, T., Kimmel, C.B., and Kimelman, D. (1995). A homeobox gene essential for zebrafish notochord development. Nature 378, 150–157.

Varshney, G.K., Pei, W., LaFave, M.C., Idol, J., Xu, L., Gallardo, V., Carrington, B., Bishop, K., Jones, M., Li, M., et al. (2015). High-throughput gene targeting and phenotyping in zebrafish using CRISPR/Cas9. Genome Res 25, 1030–1042.

Wang, Y., Kaiser, M.S., Larson, J.D., Nasevicius, A., Clark, K.J., Wadman, S.A., Roberg-Perez, S.E., Ekker, S.C., Hackett, P.B., McGrail, M., et al. (2010). Moesin1 and Ve-cadherin are required in endothelial cells during in vivo tubulogenesis. Development 137, 3119–3128.

Yang, H., Wang, H., Shivalila, C.S., Cheng, A.W., Shi, L., and Jaenisch, R. (2013). One-step generation of mice carrying reporter and conditional alleles by CRISPR/Cas-mediated genome engineering. Cell 154, 1370–1379.

Zhang, J.P., Li, X.L., Li, G.H., Chen, W., Arakaki, C., Botimer, G.D., Baylink, D., Zhang, L., Wen, W., Fu, Y.W., et al. (2017). Efficient precise knockin with a double cut HDR donor after CRISPR/Cas9-mediated double-stranded DNA cleavage. Genome Biol 18, 35.

Zhang, Y., Zhang, Z., and Ge, W. (2018). An efficient platform for generating somatic point mutations with germline transmission in the zebrafish by CRISPR/Cas9-mediated gene editing. J Biol Chem 293, 6611–6622.

Zu, Y., Tong, X., Wang, Z., Liu, D., Pan, R., Li, Z., Hu, Y., Luo, Z., Huang, P., Wu, Q., et al. (2013). TALEN-mediated precise genome modification by homologous recombination in zebrafish. Nat Methods 10, 329–331.

## References

Jao, L.E., Wente, S.R., and Chen, W. (2013). Efficient multiplex biallelic zebrafish genome editing using a CRISPR nuclease system. Proc Natl Acad Sci U S A 110, 13904–13909.

Won, M., and Dawid, I.B. (2017). PCR artifact in testing for homologous recombination in genomic editing in zebrafish. PLoS One 12, e0172802.

